# A view of the pan-genome of domesticated cowpea (*Vigna unguiculata* [L.] Walp.)

**DOI:** 10.1101/2022.08.22.504811

**Authors:** Qihua Liang, María Muñoz-Amatriaín, Shengqiang Shu, Sassoum Lo, Xinyi Wu, Joseph W. Carlson, Patrick Davidson, David M. Goodstein, Jeremy Phillips, Nadia M. Janis, Elaine J. Lee, Chenxi Liang, Peter L. Morrell, Andrew D. Farmer, Pei Xu, Timothy J. Close, Stefano Lonardi

**Affiliations:** Department of Computer Science and Engineering, University of California, Riverside, CA, 92521, USA; Department of Botany & Plant Sciences, University of California, Riverside, CA, 92521, USA; US Department of Energy Joint Genome Institute, Walnut Creek, CA, 94598, USA; State Key Laboratory for Managing Biotic and Chemical Threats to the Quality and Safety of Agro-products, Institute of Vegetables, Zhejiang Academy of Agricultural Sciences, 198 Shiqiao Road, Hangzhou 310021, China; Department of Agronomy and Plant Genetics, University of Minnesota, St. Paul, MN, 55108, USA; National Center for Genome Resources, Santa Fe, NM, 87505, USA; Key Lab of Specialty Agri-Product Quality and Hazard Controlling Technology of Zhejiang Province, College of Life Sciences, China Jiliang University, Hangzhou, China

**Keywords:** Cowpea, *Vigna unguiculata*, pan-genome, genetic variation, SNPs, presence/absence variants (PAV), inversions, IT97K-499-35, Suvita-2, Sanzi, CB5-2, UCR779, TZ30, ZN016

## Abstract

Cowpea, *Vigna unguiculata L*. Walp., is a diploid warm-season legume of critical importance as both food and fodder in sub-Saharan Africa. This species is also grown in Northern Africa, Europe, Latin America, North America, and East to Southeast Asia. To capture the genomic diversity of domesticates of this important legume, *de novo* genome assemblies were produced for representatives of six sub-populations of cultivated cowpea identified previously from genotyping of several hundred diverse accessions. In the most complete assembly (IT97K-499-35), 26,026 core and 4,963 noncore genes were identified, with 35,436 pan genes when considering all seven accessions. GO-terms associated with response to stress and defense response were highly enriched among the noncore genes, while core genes were enriched in terms related to transcription factor activity, and transport and metabolic processes. Over 5 million SNPs relative to each assembly and over 40 structural variants >1 Mb in size were identified by comparing genomes. Vu10 was the chromosome with the highest frequency of SNPs, and Vu04 had the most structural variants. Noncore genes harbor a larger proportion of potentially disruptive variants than core genes, including missense, stop gain, and frameshift mutations; this suggests that noncore genes substantially contribute to diversity within domesticated cowpea.

**Article Summary:** This study reports annotated genome assemblies of six cowpea accessions. Together with the previously reported annotated genome of IT97K-499-35, these constitute a pan-genome resource representing six subpopulations of domesticated cowpea. Annotations include genes, variant calls for SNPs and short indels, larger presence or absence variants, and inversions. Noncore genes are enriched for loci involved in stress response and harbor many genic variants with potential effects on coding sequence.

## Introduction

Individuals within a species vary in their genomic composition. The genome of any individual does not include the full complement of genes contained within the species. A pan-genome includes genes core to the species (shared among all individuals) and those absent from one or more individuals (noncore, dispensable, or variable genes). This pan-genome concept started to be applied to plants by Morgante *et al*. (2007) but began in bacterial species (reviewed by Golicz *et al*., 2020). Due to the complexity of plant genomes, the first studies exploring gene presence-absence variation (PAV) in plants used reduced-representation approaches, including array comparative genomic hybridization (CGH) and sequencing of transcriptomes (e.g., Springer *et al*. 2009, Muñoz-Amatriaín *et al*. 2013; Hirsch *et al*. 2014). Once sequencing of multiple plant genomes became feasible, several pan-genomes of variable degrees of completeness were generated, and it was soon understood that PAV is prevalent in plants and that the pan-genome of any plant species is larger than the genome of any individual accession (reviewed by Lei *et al*. 2021). Moreover, many of the genes absent in reference accessions have functions of potential adaptive or agronomic importance, such as time to flowering, and response to abiotic and biotic stresses (Gordon *et al*. 2017; Montenegro *et al*. 2017; Bayer *et al*. 2020), making the construction of a pan-genome a crucial task for crops of global importance.

Cowpea is a diploid (*2n* = 22) member of the family Fabaceae tribe Phaseoleae, closely related to mung bean, common bean, soybean, and several other warm-season legumes. Cowpea was domesticated in Africa, but its cultivation has spread throughout most of the globe (Herniter *et al*., 2020). The inherent resilience of the species to drought and high temperatures (Hall 2004), together with its nutritional value as a reliable source of plant-based protein and folic acid, position cowpea favorably as a component of sustainable agriculture in the context of global climate change. Most cowpea production and consumption presently occur in sub-Saharan Africa, especially in the Sudano-Sahelian Zone, with production mainly by smallholder farmers, often as an intercrop with maize, sorghum, or millet (Boukar *et al*., 2019). Tender green seeds are often consumed during the growing season, and immature pods are eaten as a vegetable, especially in East and Southeast Asia. In addition, fresh leaves are sometimes consumed, and dry haulms are harvested and sold as fodder for livestock. Spreading varieties are also utilized as cover crops to prevent soil erosion and weed control.

A reference genome sequence of cowpea cv. IT97K-499-35 was previously generated (Lonardi *et al*., 2019). Preliminary sequence comparisons using whole genome shotgun (WGS) data of 36 accessions suggested that extensive SNP and structural variation exists within domesticated cowpea (Lonardi *et al*., 2019). Cowpea also displays a wide range of phenotypic variation, and genetic assignment approaches have identified six subpopulations within cultivated cowpea germplasm (Muñoz-Amatriaín *et al*., 2021). These observations support the need to develop cowpea pan-genome resources based on diverse cowpea accessions.

This study reports *de novo* assemblies of six cultivated cowpea accessions. Each accession was annotated using transcriptome sequences from the accession along with *ab initio* methods. These genome sequences, together with the previously reported sequence of IT97K-499-45 (Lonardi *et al*., 2019), constitute a pan-genome resource for domesticated cowpea. Using annotations for the seven genomes, including genes, along with variant calls for SNPs and short indels, and larger structural variants, the following questions were addressed: (i) What proportion of genes are core and noncore, and do core and noncore genes differ in size or functional class? (ii) What proportion of large-effect variants are created by single nucleotide variants versus structural variants (including indels), and do the proportions of large-effect variants differ among core and noncore genes? (iii) To what extent are gene content and gene order consistent across accessions within the species *V. unguiculata* and across species within the genus *Vigna* and the tribe Phaseoleae? The results suggest that both extensive structure differences among individual accessions and the nature of variation in noncore genes are important considerations in efforts to identify genetic variation with adaptive potential.

## Materials & Methods

### Cowpea accessions selected for sequencing (Supplemental Table S01)

Accessions chosen for sequencing and *de novo* assembly represented the six subpopulations of domesticated cowpea described in Muñoz-Amatriaín et al. (2021), as indicated in Figure 1. The intention of choosing accessions that cover each subpopulation was to maximize the discovery of genetic variations relevant to cultivated cowpea using a small number of samples. As shown by Gordon *et al*. (2017) in *Brachypodium distachyon*, the addition of individuals from subpopulations not previously sampled contributes much more to increasing the pan-genome size than adding closely related individuals.

**Figure 1.**
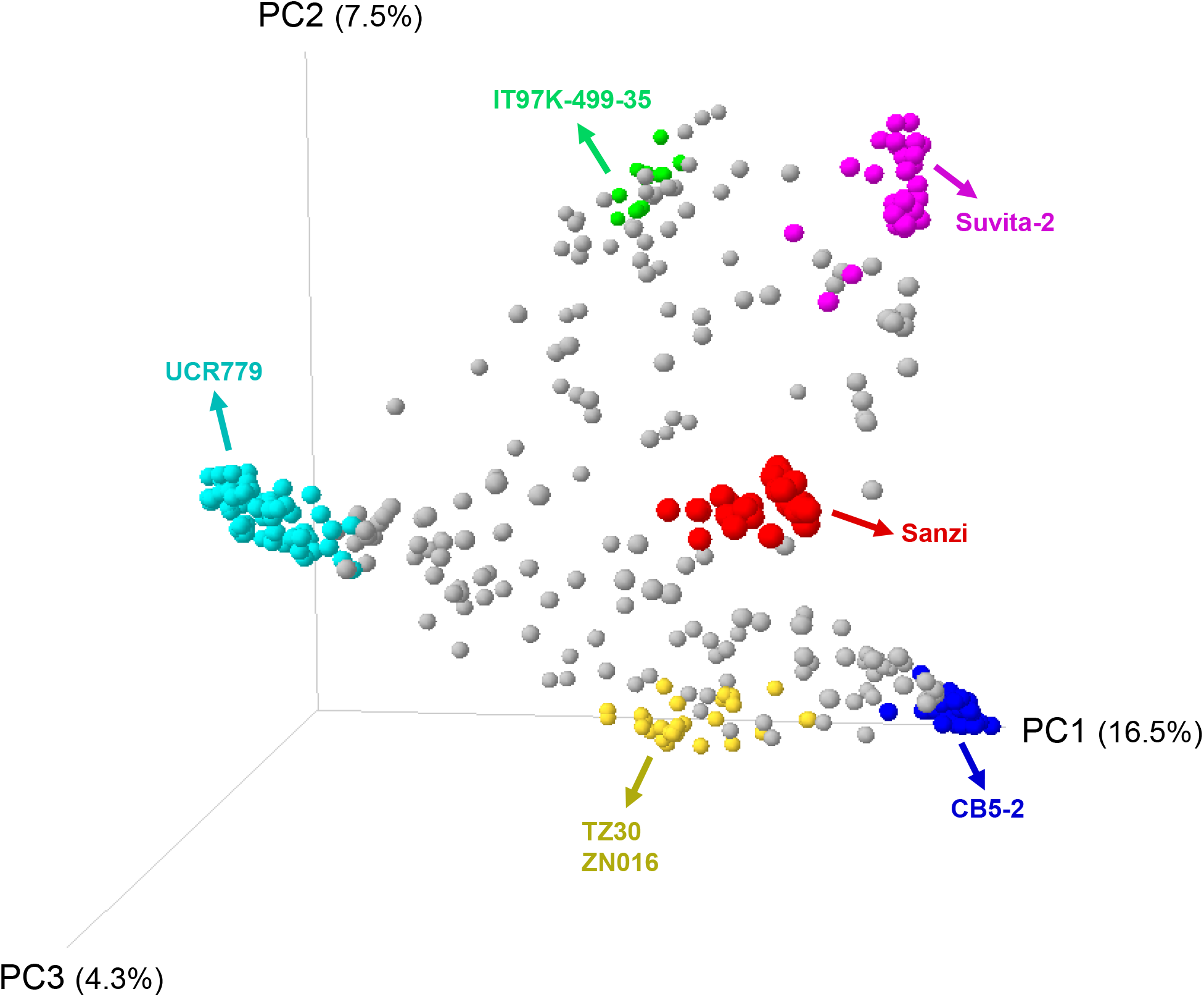
Principal component analysis of the UCR Minicore, indicating the accessions selected for sequencing and the subpopulation they belong to. Accessions in the plot are colored by the result of STRUCTURE for *K*=*6*, as shown in Muñoz-Amatriaín *et al*. (2021).

IT97K-499-35 is a blackeye variety with resistance to the parasitic plants Striga and Alectra, developed at the International Institute of Tropical Agriculture in Ibadan, Nigeria (Singh *et al*., 2006) and provided by Michael Timko (U Virginia, Charlottesville, Virginia, USA) to the University of California Riverside (UCR) in 2006. The sequence assembly and annotation of IT97K-499-35 were described in Lonardi *et al*. (2019). CB5-2 is a fully inbred isolate closely related to CB5, the predominant Blackeye of the US Southwest for several decades. CB5 (Blackeye 8415) was bred by WW Mackie at the University of California (Mackie, 1946) to add resistances to Fusarium wilt and nematodes to a California Blackeye landrace, and provided to UCR by K Foster, University of California, Davis, in 1981. Suvita-2, also known as Gorom Local (IITA accession TVu-15553, US NPGR PI 583259), is somewhat resistant to bruchids and certain races of Striga and is relatively drought tolerant. This landrace was collected from a local market by VD Aggarwal at the Institut de l’Environnement et de Recherches Agricoles (INERA) in Burkina Faso (Aggarwal *et al*.,1984) and provided to UCR by VD Aggarwal in 1983. Sanzi is an early flowering, small-seeded landrace from Ghana with resistance to flower bud thrips (Boukar *et al*., 2013), provided by KO Marfo, Nyankpala Agricultural Experiment Station, Tamale, Ghana to UCR in 1988. UCR779 (PI 583014) is a landrace from Botswana (de Mooy, 1985; Ehlers *et al*., 2002) that was provided to UCR as B019-A in 1987 by CJ de Mooy of Colorado State University. Yardlong bean or asparagus bean (cv.-gr. Sesquipedalis), the vegetable type of cowpea, is widely grown in Asian countries for the consumption of tender long pods. TZ30 is an elite Chinese variety with a pod length of around 60 cm. ZN016 is a landrace originating from southeastern China with a pod length of about 35 cm and showing resistance to multiple major diseases of cowpea. TZ30 and ZN016 were used previously as parents of a mapping population to study the inheritance of pod length (Xu *et al*., 2017).

### DNA sequencing and *de novo* assembly of seven cowpea accessions

The annotated genome (v1.0) of African variety IT97K-499-35 was assembled from Pacific Biosciences (Menlo Park, California, USA) long reads, two Bionano Genomics (San Diego, California, USA) optical maps and ten genetic linkage maps as described previously (Lonardi *et al*., 2019). The six additional *de novo* assemblies were produced by Dovetail Genomics (Scotts Valley, California, USA) using Illumina (San Diego, California, USA) short reads (150×2). DNA was extracted by Dovetail Genomics from seedling tissue of CB5-2, TZ30, and ZN016, and seeds of CB5-2, Suvita-2, Sanzi, and UCR779. Meraculous (Chapman *et al*., 2011) was used to assemble the reads, then sequences from Dovetail Chicago® and Dovetail Hi-C® libraries were added (using their proprietary pipeline) to resolve misassemblies and increase contiguity. These assemblies were further refined using ALLMAPS (Tang *et al*., 2015). This analysis used ten previously reported genetic linkage maps to relate assemblies to the standard orientations and numbering of the eleven cowpea chromosomes, as described in Lonardi *et al*. (2019) for IT97K-499-35. See “Data Availability Statement” for access to raw data and assemblies.

### Calling of SNPs, indels, and structural variants

SNPs and indels were called using each reference genome versus the reads from the six other accessions. Reads of each accession described above for genome assemblies, plus short-read sequences produced by 10X Genomics from IT97K-49-35, were mapped to all assemblies using BWA (Li *et al*., 2009). SNPs and indels were called using the GATK 4.2.0 pipeline in GVCF mode for each accession. All the per-sample GVCFs were gathered in joint genotyping to produce a set of joint-called SNPs and indels. Both per-sample SNPs and joint-called SNPs were filtered with the same parameters of ‘QD < 2.0 || FS > 60.0 || MQ < 40.0 || MQRankSum < −12.5 || ReadPosRankSum < −8.0 || SOR > 4.0’. Indels were filtered with ‘QD < 2.0 || FS > 200.0 || ReadPosRankSum < −20.0 || SOR > 10.0’.

Each pair of individual genomes was aligned using minimap2 (Li, 2018), producing Q) 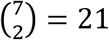 alignment files. Structural variants, including inversions and translocations, were identified from the alignment files using SyRI (Goel *et al*., 2019). Figures were produced using PlotSR (Goel *et al*., 2022). Depth analyses were carried out using Mosdepth (Pedersen & Quinlan 2018). The average nucleotide diversity within and between populations was calculated from a VCF file using Pixy (Korunes *et al*., 2021).

### Annotation of genes and repeats

All genomes were annotated using the JGI plant genome annotation pipelines (Shu *et al*., 2014), integrated gene call (IGC), and gene model improvement (GMI). Both IGC and GMI are evidence-based gene call pipelines. In IGC, a gene locus was defined by peptide alignments of related organism homologous peptides and with alignments of within-organism transcriptome assemblies. Genes were predicted by homology-based gene prediction programs FGENESH+ (Salamov and Solovyev, 2000), FGENESH_EST, and GenomeScan (Yeh *et al*., 2001), and a JGI in-house homology-constrained transcriptome assembly ORF finder. Homologous proteomes included *Arabidopsis thaliana* and those from common bean (*Phaseolus vulgaris*), soybean (*Glycine max*), barrel medic (*Medicago truncatula*), poplar (*Populus trichocarpa*), rice (*Oryza sativa*), grape (*Vitis vinifera*) and Swiss-Prot. For transcript-based annotations of the six new assemblies, RNA for RNA-seq was extracted using Qiagen RNeasy Plant (Hilden, Germany) from each accession from well-hydrated and drought-stressed young seedling root and leaves, immature flower buds, and pods five days after pollination, and from developing seeds of Suvita-2, TZ30 and ZN016 (not CB5-2, Sanzi or UCR779) 13 days after pollination. RNA quality was assessed, and concentrations were determined using an Agilent 2100 BioAnalyzer (Santa Clara, California, USA) and the Agilent RNA 6000 Nano Kit. The RNA-seq short reads from each accession were assembled using a JGI in-house genome-guided assembler, PERTRAN (Shu *et al*., 2013), using each genome assembly. Each short-read-based assembly and UNIGENE sequences (P12_UNIGENES.fa from harvest.ucr.edu) were fed into PASA (Haas *et al*., 2003) to produce transcriptome assemblies. The best gene per locus (based on evidence) was defined using PASA from alignment of transcriptome assemblies for splicing correctness, alternative transcripts, and UTR addition. The PASA genes were filtered to obtain the final gene set, including an automated repeat coding sequence (CDS) overlap filter, a manual low-quality gene filter, and an automatic filter from transposable element (TE) protein domain assignments. This process was repeated once with one additional homology seeding of non-self, high-confidence gene models.

### Determination of core and noncore genes among seven accessions

Core and noncore genes were determined by running the GET_HOMOLOGUES-EST tool (https://github.com/eead-csic-compbio/get_homologues) on the primary transcripts of the seven cowpea accessions provided in nucleotide and protein formats. GET_HOMOLOGUES-EST was run in orthoMCL-mode, as suggested by the authors for pan-genome analyses (Contreras-Moreira *et al*., 2017). The other GET_HOMOLOGUES-EST options “**-M -c -z -t 0 –A -L**” were used to obtain orthoMCL gene clusters, which had genes in 1-7 accessions. The term “core” means that a matching gene was identified in all seven accessions and “noncore” means that a matching copy gene was identified in less than all seven accessions.

GO-term enrichment analyses were performed in agriGO v2.0 (Tian *et al*., 2017) for core and noncore genes using GO terms available from the Legume Information System (https://www.legumeinfo.org/). Given the large number of GO terms in both the core and noncore gene sets, GO slims (Onsongo *et al*., 2008) were extracted and used for Figure 3. The full list of core and noncore genes, with GO and other annotations, is available from the Google Drive noted in the Data Availability Statement.

### Annotation of variants in core and noncore genes

To test if variants in noncore genes have been subject to reduced selective constraint, Variant Effect Predictor (VeP) (McLaren *et al*., 2016) was used to annotate variants identified in the primary transcripts of core and noncore genes. Gene annotations for IT97K-499-35 were used to identify intervals that overlap core and noncore genes, and filtering of the VCF file used BEDtools intersect (Quinlan & Hall, 2010) with variants called relative to the IT97K-499-35 assembly using the six other assemblies. Scripts used for these analyses are at https://github.com/MorrellLAB/Cowpea_Pangenome. VeP was run separately for SNPs and indels, reporting classes of variants with potentially large effects, including missense, stop gains, start or stop changes, and frameshifts. The numbers of synonymous changes and in-frame indels are also reported.

### Relative size of core and noncore genes

The physical sizes of core and noncore genes were compared in the total annotated length and the length of the coding portion of the primary transcript of each gene. The length of each gene was extracted from the general feature format (GFF) annotations. The CDS length was calculated based on the primary transcript identified in Phytozome annotations (https://phytozome-next.jgi.doe.gov/cowpeapan/info/Vunguiculata_v1_2). The full list of core and noncore genes, with gene and CDS sizes indicted, is available from the Google Drive noted in the Data Availability Statement.

### Nucleotide sequence diversity in cowpea

Tajima’s (1983) estimate of θ = 4Neμ was used to determine the level of sequence diversity in the pangenome accessions. “Callable” regions were identified based on coverage estimates in mosdepth (Pederson & Quinlan, 2018), with “callable” regions defined as those with coverage between 5x and 400x. This estimate was derived from a sample with ~200X average coverage. The callable regions were used to create a BED file used for filtering genomic regions. This approach is intended to avoid variant calls in regions with inadequate sequence depth or regions where very high coverage may indicate non-unique mapping of sequence reads. The callable regions and the VCF file of filtered variants mapped to the IT97K-499-35 reference were used with pixy (Korunes & Samuk, 2020), a tool designed to deal with missing data in genome-level resequencing datasets.

### Physical locations of SNPs from genotyping platforms

The physical positions of SNPs in the Illumina iSelect Cowpea Consortium Array (Muñoz-Amatriaín *et al*., 2017), whose positions in the IT97K-499-35 genome were provided in Lonardi *et al*. (2019), were mapped using BWA MEM (Li *et al*., 2009) within each of the seven assemblies using the contextual sequence that flanked each variant. The resulting alignment file was processed with SAMtools (Li *et al*., 2009) and SNP_Utils (https://github.com/MorrellLAB/SNP_Utils) to report positions in a VCF file. The positions of iSelect SNPs relative to all seven genome assemblies are provided in Supplemental Table S02, and an updated summary map for the 51,128 iSelect SNPs is in Supplemental Table S03. The positions identified for iSelect SNPs relative to the IT97K-499-35 assembly were used to annotate the variants. The annotation used variant effect predictor (VeP) (McLaren *et al*., 2016) with the GFF file provided by Phytozome (https://genome.jgi.doe.gov/portal/) and SNP positions in VCF files (https://github.com/MorrellLAB/cowpea_annotation/blob/main/Results/IT97K-499-35_v1.0/iSelect_cowpea.vcf; see Data Availability Statement).

### Synteny analysis among genome assemblies

To assess the conservation of gene content and ordering between genome assemblies from diverse species, MCScanX (Wang *et al*., 2012) was run for every genome pair, using default settings and homologous gene pairings derived from gene family assignments defined as the best match of the longest protein product with an E-value of 1e-10 or better from hmmsearch (Eddy 2011) applied to the legfed_v1_0 families (Stai *et al*., 2019).

## Results and Discussion

### Development of six *de novo* assemblies and pan-genome construction

Summary statistics for the seven assemblies (assembly characteristics, repetitive content, genes, BUSCO completeness) are reported in Table 1. More detailed statistics of the intermediate assembly steps are reported in Supplemental Table S04. The contiguity of the new six assemblies, as indicated by their N50s, is comparable to the PacBio assembly for IT97K-499-35 despite being based on short-read sequences. In all six new assemblies, each of the eleven chromosomes of cowpea is represented by a single scaffold. These six assembled genomes are similar to each other in size, ranging from 447.58 Mb to 453.97 Mb, with a mean of 449.91 Mb. IT97K-499-35 had a ~15% larger (more complete) assembled size (519.44 Mb) than these six accessions, with the difference attributable to long-read sequencing and optical mapping providing a more complete assembly. Assemblies of the six additional accessions share the same percentage of repetitive content of about 45-46% (Table 1 and Supplemental Figure S1). The IT97K-499-35 assembly has a somewhat higher repetitive content than the assemblies of these six accessions. This may be attributable to more complete resolution of unique positions of repetitive sequences within long sequence reads than is possible from only short reads. A difference between the sequencing methods in the resolution of repetitive sequences is evident in centromeric regions, which are typically abundant in repetitive sequences, where some chromosomes of the six newly sequenced accessions appear to be missing from the assemblies. Centromeric regions were defined based on a 455-bp tandem repeat previously identified by fluorescence in situ hybridization (Iwata-Otsubo *et al*., 2016). Supplemental Table S05 shows the coordinates of the putative centromeric regions in IT97K-499-35 for all eleven chromosomes for a total span of 20.18 Mb, in CB5-2 on five chromosomes for a total span of 5.6 Mb, in Sanzi on one chromosome for a total span of 0.59 Mb, in ZN016 on four chromosomes for a total of 7.13 Mb and TZ30 on one chromosome for 1.32 Mb. The tandem repeat was not found in any assembled chromosome of Suvita-2 or UCR779, nor in the other chromosome assemblies where coordinates are not listed.

**Table 1.**
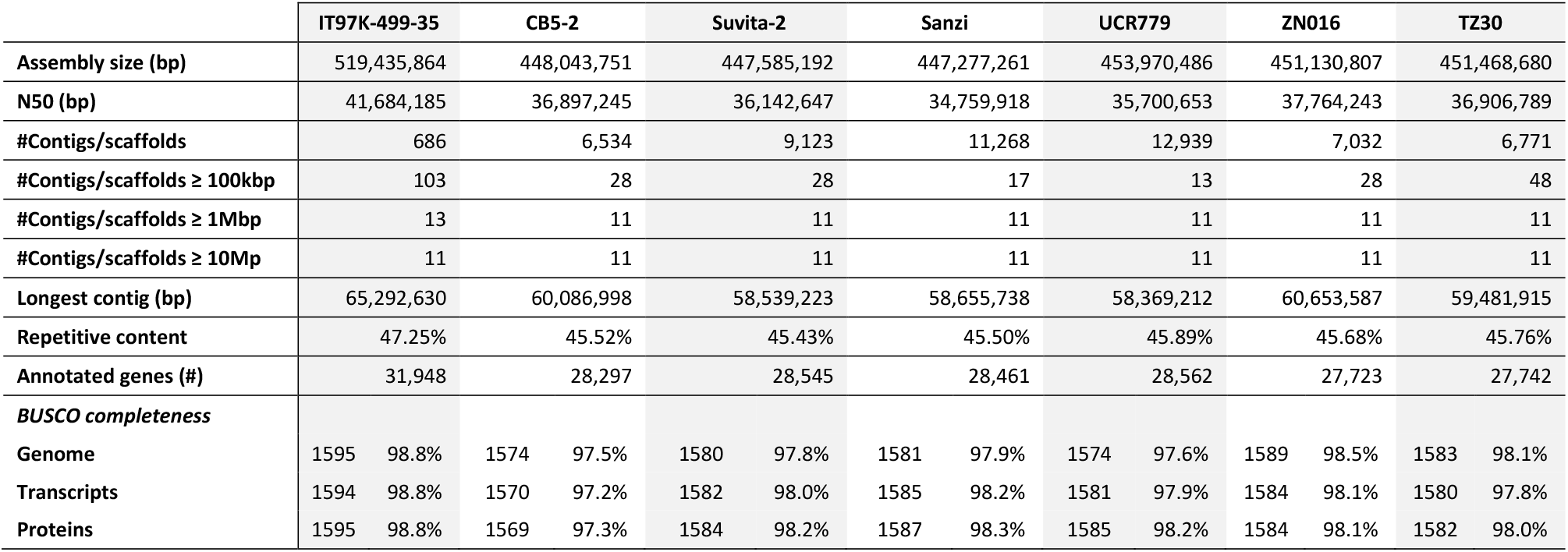
Summary of assembly statistics, repetitive content, gene content and BUSCO4 completeness for the seven genomes.

RNA was prepared from each accession to support gene annotation, and the same annotation protocol was applied to each accession (see Materials & Methods). This is important when comparing genomes at the gene level, as it reduces the technical variability that can otherwise obfuscate the interpretation of results (Lei *et al*. 2021). The number of genes annotated in the six new assemblies ranged from 27,723 to 28,562, with a mean of 28,222 (Table 1). IT97K-499-35 had ~13% more annotated genes, with a total of 31,948, reflecting deeper transcriptome sequencing and, to some extent, the more complete assembly of its genome. Supplemental Table S06 summarizes the number of alternative transcripts, exon statistics, gene model support, and ontology annotations (Panther, PFam, KOG, KEGG, and E.C.). The number of alternative transcripts in the six new assemblies ranged from 15,088 to 17,115. Again, IT97K-499-35 had a higher number of alternative transcripts, a total of 22,536, than the other six accessions. The average number of exons was 5.4 in each of the six new assemblies and 5.2 in IT97K-4899-35, with a median length ranging from 162 to 169 bp. Gene and repeat density were computed in 1Mb non-overlapping sliding windows along each chromosome and each accession (Supplemental Figure S1). All chromosomes have a higher gene density in their more recombinationally active regions, while repeat density peaks in the low-recombination centromeric and pericentromeric regions (see also Supplemental Figure S8 in Lonardi *et al*., 2019). All seven accessions have similar gene and repeat density, and high BUSCO v4 completeness at the genome, transcript, and protein levels (Supplemental Table S07), with somewhat higher numbers for IT97K-499-35 than the six new assemblies.

As stated above (Materials and Methods), genes annotated in the seven genomes were classified as core if a matching gene was present in all accessions and noncore if absent in one or more of the seven accessions. In IT97K-499-35, a total of 26,026 core genes (in 24,476 core clusters) and 4,963 noncore genes (in 4,285 noncore clusters) were identified (Supplemental Table S08). When considering all seven accessions itemized in Supplemental Table S08, a total of 26,494 core genes and 9,042 noncore genes (in 8,157 noncore clusters) were identified, resulting in a total of 35,536 pan genes in 32,633 pan gene clusters.

To determine if adding accessions significantly changed the numbers and proportions of core and noncore genes, we took advantage of the analysis results produced by GET_HOMOLOGUES-EST. GET_HOMOLOGUES-EST produces pan or core genome growth simulations by adding accessions in random order, using twenty permutations. Figure 2 shows the growth of core and pan genomes for an increasing number of accessions. A fitted Tettelin function (Tettelin *et al*., 2005) is plotted in green. As expected, the number of pan genes increases as additional accessions are “added” to the pan-genome, while the number of core genes decreases. However, the fact that the core gene plot is flattening considerably (approaching an asymptotic limit) for six and seven accessions indicates that most core genes have been identified with these seven diverse accessions. In contrast, the pan-genome plot has not flattened, indicating that there may be many more noncore genes not included among these seven accessions. Figure 2 provides an estimated 29,659 pan gene clusters and an estimated 24,439 core gene clusters as the output of GET_HOMOLOGUES-EST from 20 random samplings. Roughly, it appears that the pan-genome defined by the seven cultivated cowpea accessions is comprised of about 80% core genes, constituting nearly the entire set of core genes in cultivated cowpea, and 20% noncore genes. Clearly, more noncore genes would be revealed with a larger number of accessions.

**Figure 2.**
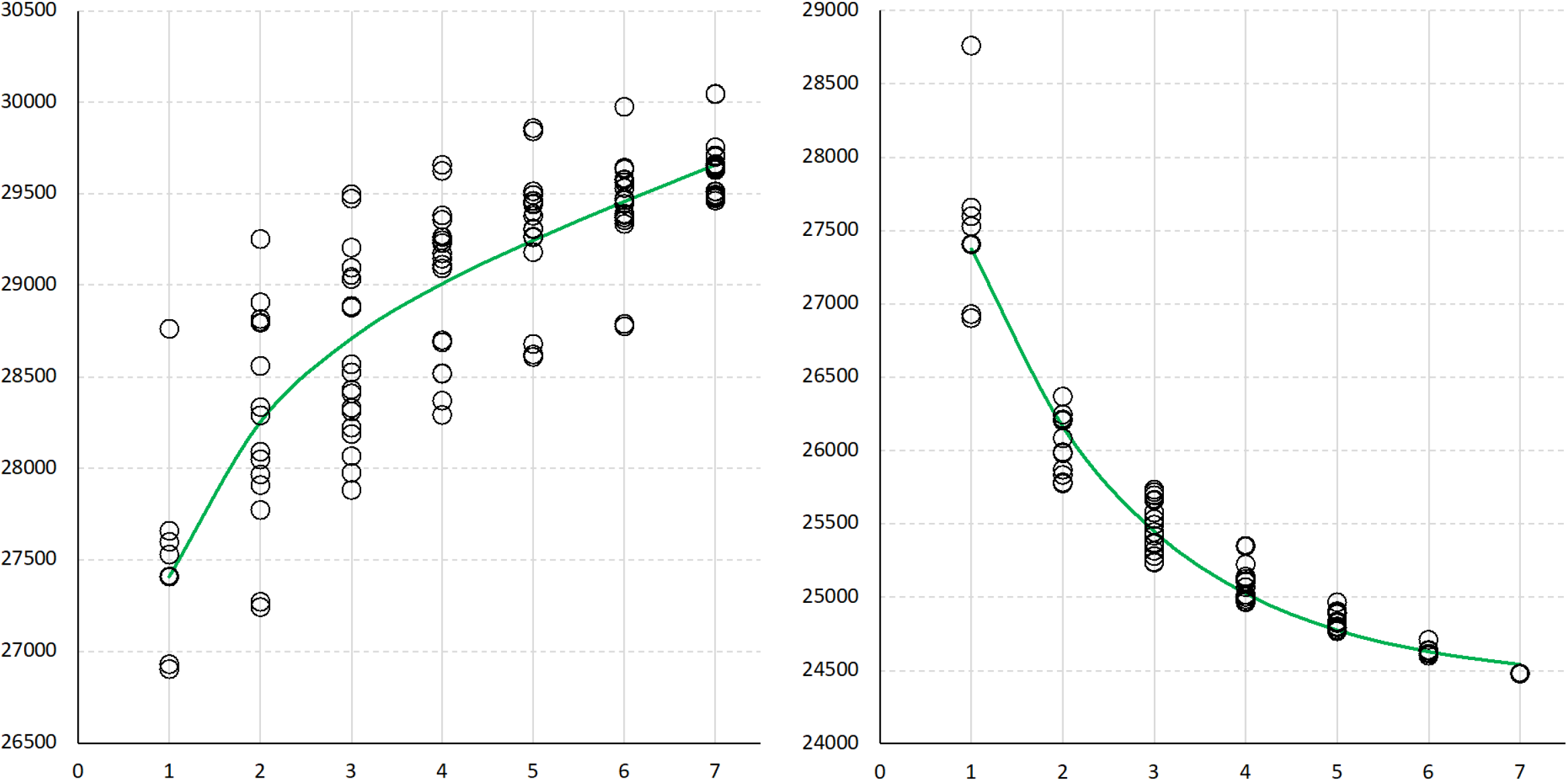
The number of genes identified in the pan-genome (left) and core genome (right) as new accessions are added. Green curves are fitted Tettelin functions.

**Figure 3.**
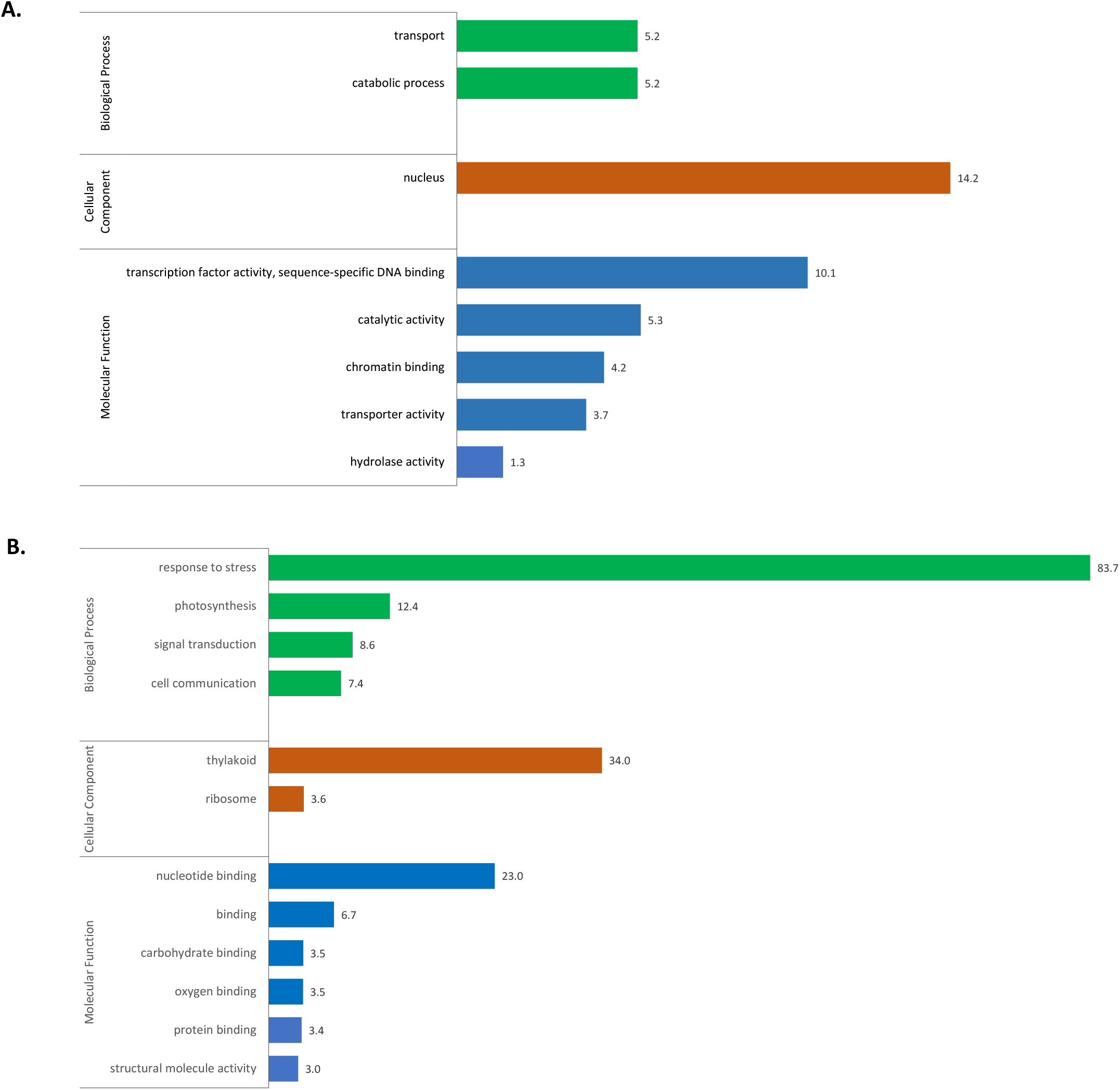
Gene Ontology (GO) term enrichment analysis. Significantly enriched GO terms for core (A) and noncore genes (B) are shown for GO-Slim categories belonging to Biological Process, Cellular Component, and Molecular Function aspects (in different colors). -log_10_ of FDR-adjusted p-values (q-values) are shown on the right of each bar.

A GO term enrichment analysis was performed for genes within the two components of the pan-genome (core and noncore) using agriGO v2 (Tian *et al*., 2017). Many GO terms for all three ontology aspects (biological process, cellular component, and molecular function) were significantly enriched in both core and noncore genes (Supplemental Table S09). Given the high number of significant GO terms, GO Slim terms (Onsongo *et al*., 2008) were extracted and used for Figure 3. Terms enriched in the core genes were related to transport and some metabolic processes and molecular functions involving DNA-binding transcription factor activity (Figure 3; Supplemental Table S09). This supports the idea that the core genome contains genes that perform essential cellular functions that are highly conserved at the species level. The output was quite different for the noncore genes, with very high enrichment of the GO term “response to stress” (Figure 3), in particular “defense response” (−log_10q_ = 123.7; Supplemental Table S09). This is consistent with previous research showing that the “dispensable” genome encodes genes involved in defense response and other beneficial functions for some individuals (Golicz *et al*., 2016; Gordon *et al*., 2017; Montenegro *et al*., 2017).

### Genetic variation analysis

In addition to identifying gene PAVs (presence-absence variants), the seven assemblies were used to identify other types of variation. Variants were detected using two different software pipelines, depending on their size. SNPs and indels of length up to 300 nucleotides, both considered small variants, were detected using GATK (see Materials & Methods). Larger structural variations, including deletions, duplications, inversions, and translocations, were detected using SyRI (Goel *et al*., 2019). Across all “callable” regions of the genome, average θ_*π*_ = 0.0111 (± 0.0549). At the pseudomolecule level, average diversity was highest on Vu05, with θ_*π*_ = 0.0155 (± 0.0723), and lowest on Vu10, with θ_*π*_ = 0.0095 (± 0.0447) (Supplemental Table S10). A mean diversity of ~1% is higher than many grain crops, such as barley (Morrell et al., 2014; Schmid et al., 2018) and roughly comparable to maize (Tittes et al., 2021). The observed diversity in the cowpea pangenome sample is above average for herbaceous plants (Miller & Gross, 2011; Leffler et al., 2012; Corbett-Detig et al., 2015).

For SNPs and indels, the genome of each accession was used in turn as the “reference,” mapping the reads for each of the six other accessions against that genome. For each, the six SNP sets produced by GATK were merged by taking the union of the SNPs based on their location (i.e., a SNP in two accessions was counted only once if it appeared in the same genomic position). Supplemental Table S11 summarizes the number of SNPs detected, where the reference genome is listed on each row. For instance, using Suvita-2 as the reference, 1,489,850 SNPs were detected using mapped reads from CB5-2, compared to 2,625,678 SNPs using the reads from UCR779. Combining the SNPs by counting all distinct SNPs in the union of the six sets of SNPs, the number of SNPs for Suvita-2 was 5,292,933.

When UCR779 was used as the reference, a much higher number of SNPs was detected in every pairwise comparison, indicating that UCR779 is the most divergent among these seven accessions. Conversely, CB5-2 (a California cultivar) has fewer SNPs in pairwise comparisons to TZ30 or ZN016 (both from China) than in pairwise comparisons to other accessions. This suggests that CB5-2 is more similar to these two accessions than to the other four accessions. This is consistent with genetic assignment analyses reported by Muñoz-Amatriaín *et al*. (2021) and historical considerations discussed in Herniter *et al*. (2020). Supplemental Table S12 provides a similar analysis for indels, where again, UCR779 stands out as the most different among the seven accessions. Summary statistics for SNPs and indels for each chromosome and each accession can be found in the file “SNPs_indels_stats.xlsx,” available from the Google Drive indicated in the Data Availability Statement below.

GATK requires a minimum coverage of 5X to call SNPs. Coverage analysis with Mosdepth indicated that the average read coverage of IT97K-499-35 is very high (e.g., about ~190X when mapping CB5-2 reads to IT97K-499-35), thus a very high fraction of IT97K-499-35 chromosomes was covered by at least five reads. The lowest was Vu10 with 85.1%, the highest was Vu07 with 98.6%, and the overall percentage of SNPs in IT97K-499-35 that were in a “callable” region (i.e., with coverage 5x-400x) was 88.96%. The frequency of SNPs, as the number of unique SNPs identified (Supplemental Table S11) divided by the size of the assembled genome (Table 1), ranges from one in 139 to one in 309 bp, and the indel frequency (Supplemental Table S12) ranges from one in 486 to one in 529 bp. Circos plots for SNP density (SNPs per Mb) on each chromosome using each accession as the reference are in Supplemental Figure S2 (A-G), where it is evident, for example, that Vu04 and Vu10 have the highest SNP frequency. In contrast, Vu05 and Vu09 have the lowest. This was observed previously when mapping nearly one million SNPs on the IT97K-499-35 reference genome (Lonardi *et al*., 2019). Also, when using UCR779 as the reference (Supplemental Figure S2-E), the number of SNPs on Vu04 and Vu10 is significantly higher than when any other accession is used as the reference, again consistent with UCR779 being the most different among the seven accessions.

Structural variations were identified using SyRI (Goel *et al*., 2019) from the alignment of each pair of individual genomes and visualized using PlotSR (Goel *et al*., 2022) (Figure 4). The visualization shows a relatively large number of apparent structural rearrangements between the seven cowpea genomes, which are more abundant in the centromeric and pericentromeric regions of all chromosomes. Vu04 is the chromosome with the highest abundance of structural variants (Figure 4). A summary of all the structural variants identified in all pairs of accessions is reported in Supplementary Table S13. The table shows that Suvita-2 versus UCR779 had the largest number of inversions (2,008) and translocations (1,822). This intuitively makes sense since these two accessions belong to two different genetic subpopulations separated by the first principal component (Figure 1).

**Figure 4.**
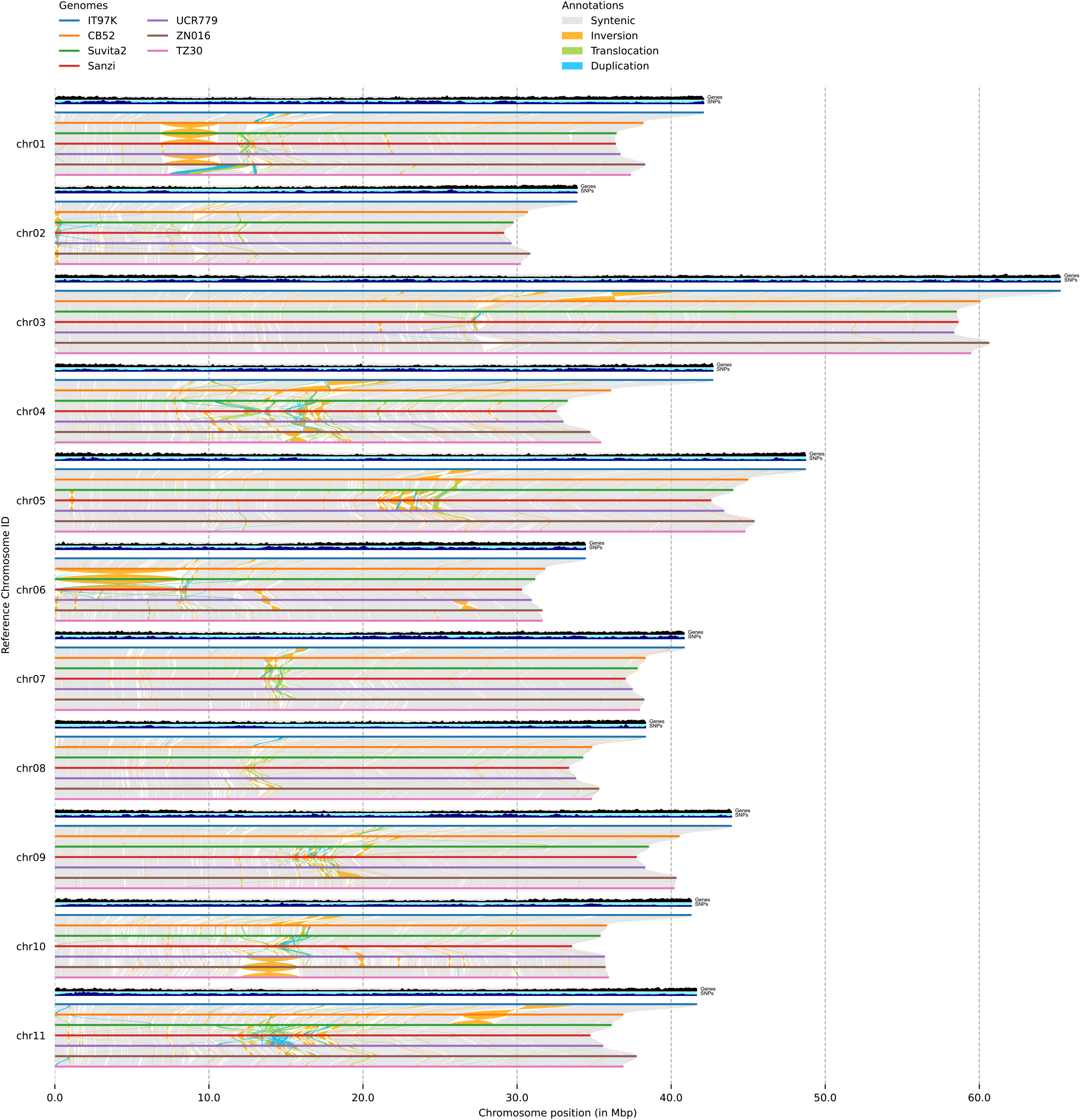
Representation of structural variations (of any size) detected by SyRI from the output of whole-genome pairwise alignments between the seven cowpea accessions. The black track indicates gene density in the reference genome IT97K-499-35, while the blue track indicates SNP density in the reference genome IT97K-499-35.

Inversions are a common type of rearrangement with important consequences for cross-over frequency and distribution, as they suppress recombination in heterozygotes (Kirkpatrick, 2010). While inversion can be important to maintaining locally adaptive variants (Kirkpatrick & Barton, 2006), crossover inhibition can impede plant breeding efforts. Table 2 summarizes the genomic coordinates of all inversions larger than 1 Mbp. For example, the first column of Table 2, corresponding to IT97K-499-35, shows 27 inversions that were identified by comparing the reference genome against the other six accessions. The same inversion can appear in multiple sub-tables. For instance, the ~4.2 Mb inversion on chromosome 3 previously described in (Lonardi *et al*., 2019) occurs in the same orientation in six accessions and the opposite orientation only in IT97K-499-35, so it is listed six times in the column for IT97K-499-35.

**Table 2.**
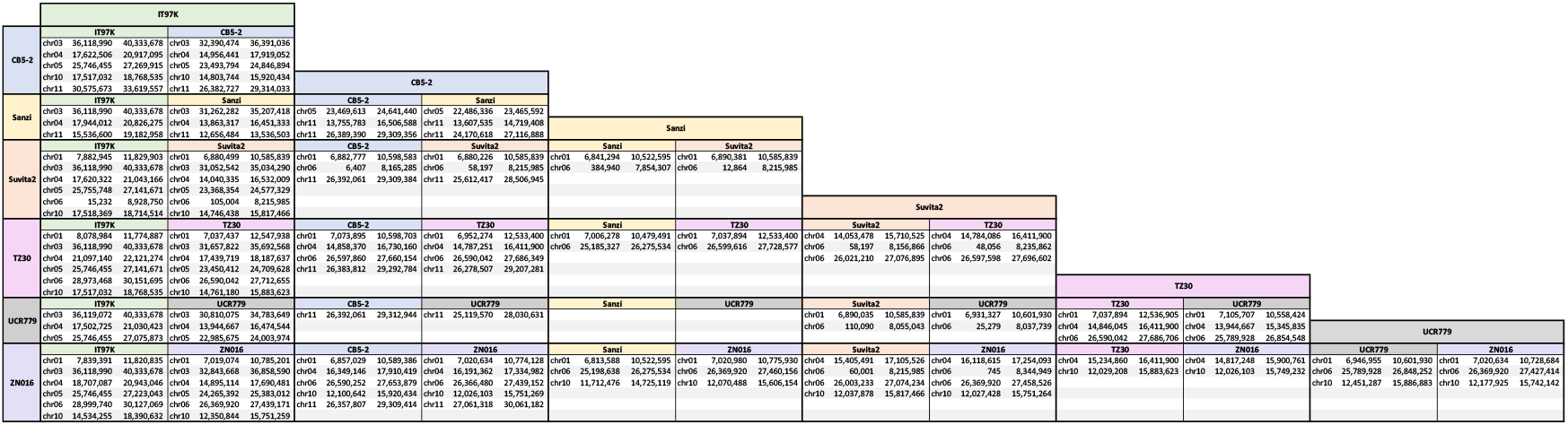
Genomic coordinates of all inversions of size > 1 Mbp detected by comparing the seven cowpea genomes pairwise. IT97K-499-35 is abbreviated as IT97K.

Similarly, the inversions on Vu04 and Vu05 are detected against five accessions. The ~9.0 Mb inversion on Vu06 is the largest inversion found by SyRI, and its orientation is unique to Suvita-2. However, this inversion appears to be due to an assembly imperfection. It is reported as unoriented in the ALLMAPS output (Supplemental Table S14), and comparisons between optical maps derived from Suvita-2 and another cowpea accession not included here indicate a non-inverted orientation in Suvita-2 (unpublished). Also, as shown in Lonardi *et al*. (2019) and Supplemental Figure S3, this entire region has a very low recombination rate and comprises nearly the entire short arm of acrocentric chromosome 6 (Iwata-Otsubo *et al*., 2016). These factors can account for a spurious orientation assignment for this region in the Suvita-2 Vu06 assembly.

The positions of the largest inversions shown in Figure 4 are provided in Table 2, e.g., the inversions on Vu03 in IT97K-499-35 reported by Lonardi *et al*. (2019), and the inversion on Vu06 in Suvita-2 likely due to a mis-assembly, as discussed above. It should be noted that regions with apparently low synteny within several chromosomes are low-recombination centromeric and pericentromeric regions (Lonardi *et al*., 2019), which are notoriously hard to assemble due to their high repetitive content and hard to orient due to a paucity of mapped and recombinationally ordered SNPs. In these regions, it is expected to find compressed contigs, gaps, and misassemblies, any of which might be flagged as apparent structural variations. The number of false-positive structural variations can likely be reduced by increasing the completeness of the assemblies within these regions using long-read sequencing and optical mapping. Supplemental Figure S4 (A-U) shows all 21 SyRi+PlotSR alignments between all pairs of cowpea accessions.

### Further characterization of core and noncore genes

Partitioning SNPs into those found in core versus noncore genes in IT97K-499-35 resulted in 702,073-SNPs in core genes and 239,100 SNPs in noncore genes. The indel comparison involves 161,900 indels in core genes and 39,845 in noncore genes. The numbers of variants with potential consequences are summarized in Figure 5 and Supplementary Table S15. Counting both SNPs and indels, there are 80,693 potentially benign variants among core genes (3.10 per gene) and 36,519 in noncore genes (7.36 per gene), which is a 2.37-fold higher frequency in noncore versus core genes. Likewise, potentially harmful variants, including missense, stop gained, start or stop change, and frameshift total 95,465 among core genes (3.67 per gene) and 75,048 in noncore genes (15.12 per gene),which is a 4.12-fold higher incidence in noncore versus core genes. Among these, noncore genes have a much higher incidence of frameshift variants (1.48 per gene) than do core genes (0.23 per gene), this being a 6.43-fold difference. In each of these comparisons, noncore genes contribute proportionally a larger number of variants than do core genes, whether benign or potentially harmful.

**Figure 5.**
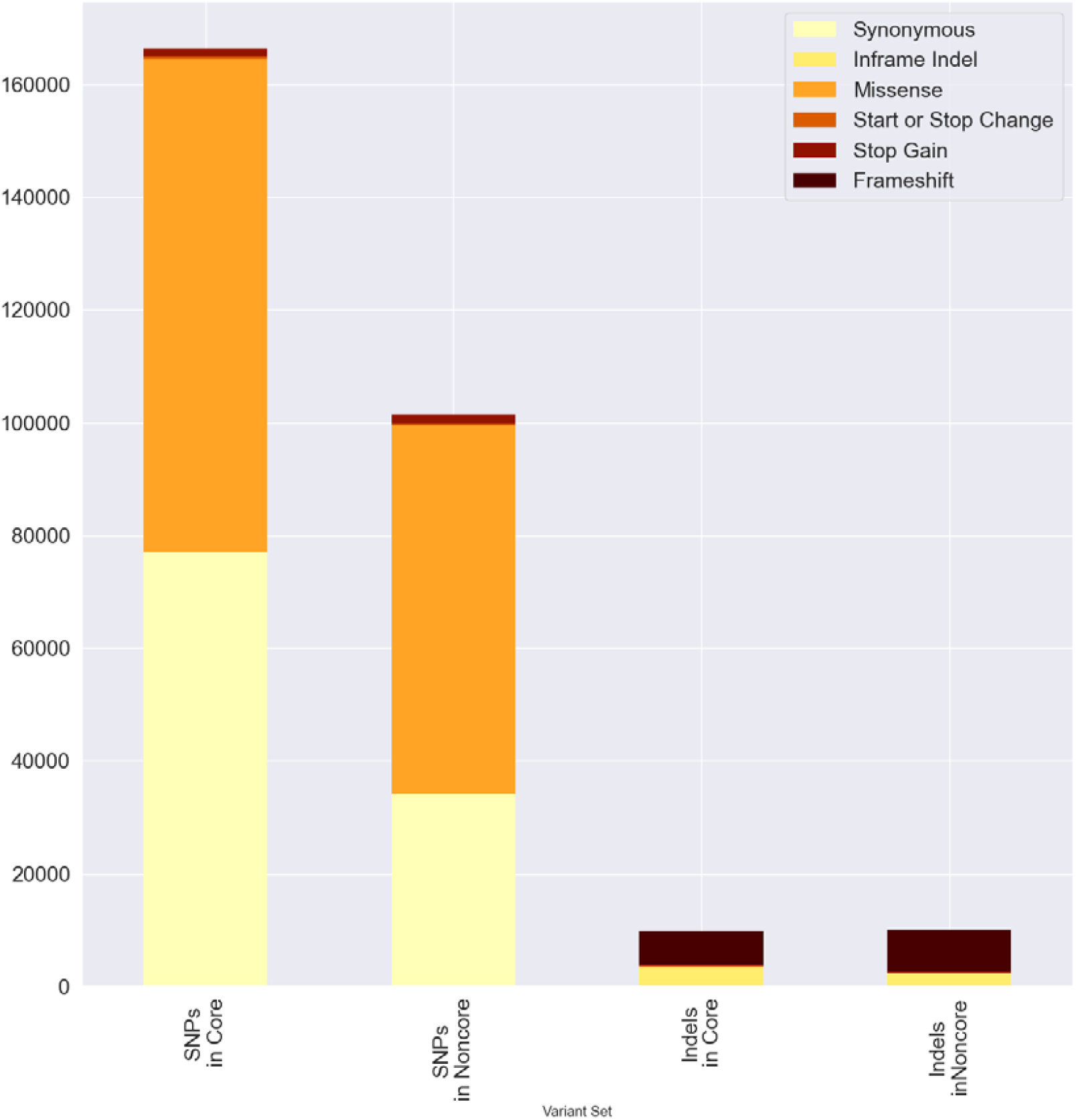
Variant effect predictor (VeP) annotations for SNPs and indels found in the core and noncore genes present in IT97K-499-35. Values on the y-axis are the absolute number of variants in each variant class.

Based on the gene annotations, core gene primary transcripts are longer than noncore gene primary transcripts, with a mean length of 4,226.08 (± 4,047.234) for IT97K-499-35 core genes versus 2,341.32 bp (± 3,190.67) for IT97K-499-35 noncore genes (with median lengths of 3,292 and 1,347 bp, respectively). This difference is significant based on a non-parametric, two-sample Wilcoxon rank sum test, with p-value < 2.2 e^-16^. For IT97K-499-35, primary transcripts from core genes cover 110.9 Mb of the genome, while primary transcripts from noncore genes cover 11.6 Mb. These differences in lengths could result from either longer coding regions or longer or more abundant introns within the primary transcripts. When considering only the coding sequence (CDS) for each IT97K-499-35 gene, the mean length of the CDS in core genes is greater than in noncore genes, with a mean of 1,319.14 (± 960.61) for core versus 792.97 bp (± 915.98) for noncore (with median lengths of 1,113 and 426 bp). Based on the Wilcoxon test, the difference in length of the coding sequence is significant, with a p-value < 2.2e^-16^. The explanation for this CDS length difference is unknown.

### Presence-absence variation of genes controlling black seed coat color

To facilitate the community’s use of the cowpea pan-genome, all the genomes and their annotations have been included as resources in the Legume Information System (LIS; www.legumeinfo.org; Dash *et al*., 2016). As an example of a use case for pan-genomics, the Genome Context Viewer (GCV) is an application that enables dynamic comparison of genomes based on their gene content, using assignments of genes to families as the basis for computation and visualization of conserved gene order and structural variation with potential impact on function, e.g., copy number variation (CNV) and presence-absence variation (PAV) (Cleary and Farmer, 2018). Figure 6A shows the results of a query centered on a region from the reference cowpea genome that features a cluster of tandemly duplicated MYB transcription factor genes in which presence-absence variation was previously determined to be associated with seed coat pigmentation (Herniter et al., 2018). The colors of the genes in this “beads on a string” representation reflect the gene family assignments; here, the brown triangles in the center of the region represent the MYB genes with varying copy numbers in the different cowpea accessions, with a maximum of five copies in the reference accession to as few as a single copy in UCR779. Outside the CNV region, there is strong conservation of gene content, with one other region showing some evidence of reordering among the cowpea accessions. The viewer facilitates comparison not only within but across species, and one can see evidence of similar CNV in the corresponding region of several *Phaseolus* spp. genomes (Schmutz *et al*. 2014, Moghaddam *et al*. 2021), as well as an inversion of the segment containing the genes relative to cowpea, soybean (Valliyodan *et al*. 2019) and other *Vigna* species (Sakai *et al*. 2015, Kang *et al*. 2014). Two corresponding homoeologous regions evidence the most recent whole genome duplication in soybean. The region serves as a breakpoint for the syntenic block in Gm09, which, taken together with the other structural variation, suggests that the expansion of gene copy number here has had consequences for the stability of the chromosome in these regions over evolutionary time (Hastings et al., 2009).

**Figure 6A.**
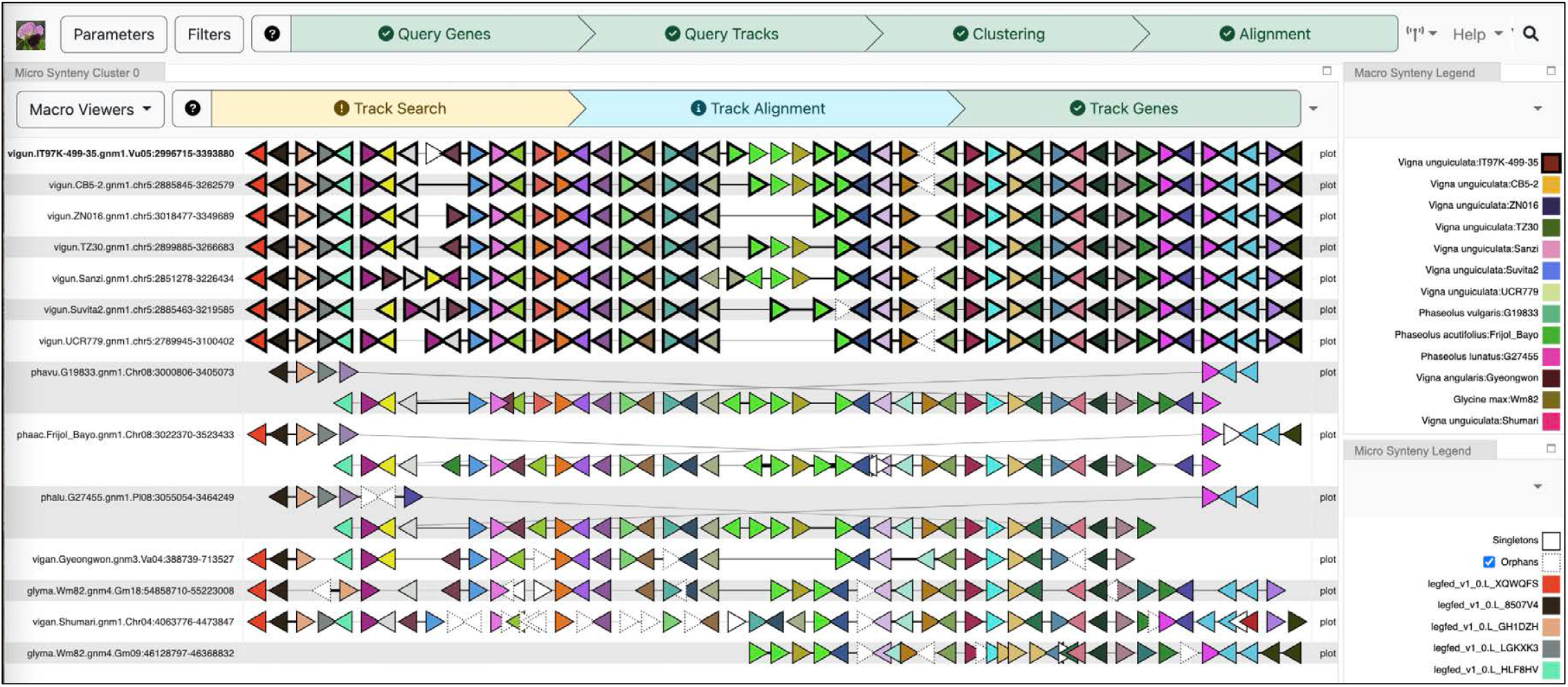
Conservation of gene content within and across species. A region depicting gene content conservation and variability among cowpea genomes and other representative Phaseoleae species. Triangular glyphs represent order and orientation of genes, with color representing gene family memberships. (https://vigna.legumeinfo.org/tools/gcv)

Although the GCV view shows good evidence for CNV, there are some limitations to what may be inferred from that alone. First, since the viewer only has access to gene family assignment information, it cannot determine which elements among those in tandem arrays have the highest sequence similarity and provide insight into which copies have been deleted. Second, because it relies on the surrounding genomic context of each gene to place it into correspondence, it will have limited capability for finding genes that are present in the assembly but are largely isolated on small scaffolds that were not incorporated into the main pseudomolecules. Another tool at LIS that provides a complementary view based on the underlying sequence identity of the different copies of the expanded gene family is shown in Figure 6B. Here, the InterMine (Kalderamis *et al*., 2014) instance for cowpea (https://mines.legumeinfo.org/cowpeamine/begin.do) was used to collect all protein sequences for cowpea genes assigned to the given family. A dynamic tree construction procedure invoked based on hmmalign-derived (http://www.csb.yale.edu/userguides/seq/hmmer/docs/node18.html; Eddy, 2011) additions of these genes to the multiple sequence alignment for the founding members of the family. The resulting tree (a subtree of which is shown) allows the user to determine the best correspondences of the copies in each genome and pulls in two additional genes on unanchored contigs that likely belong to the region.

**Figure 6B.**
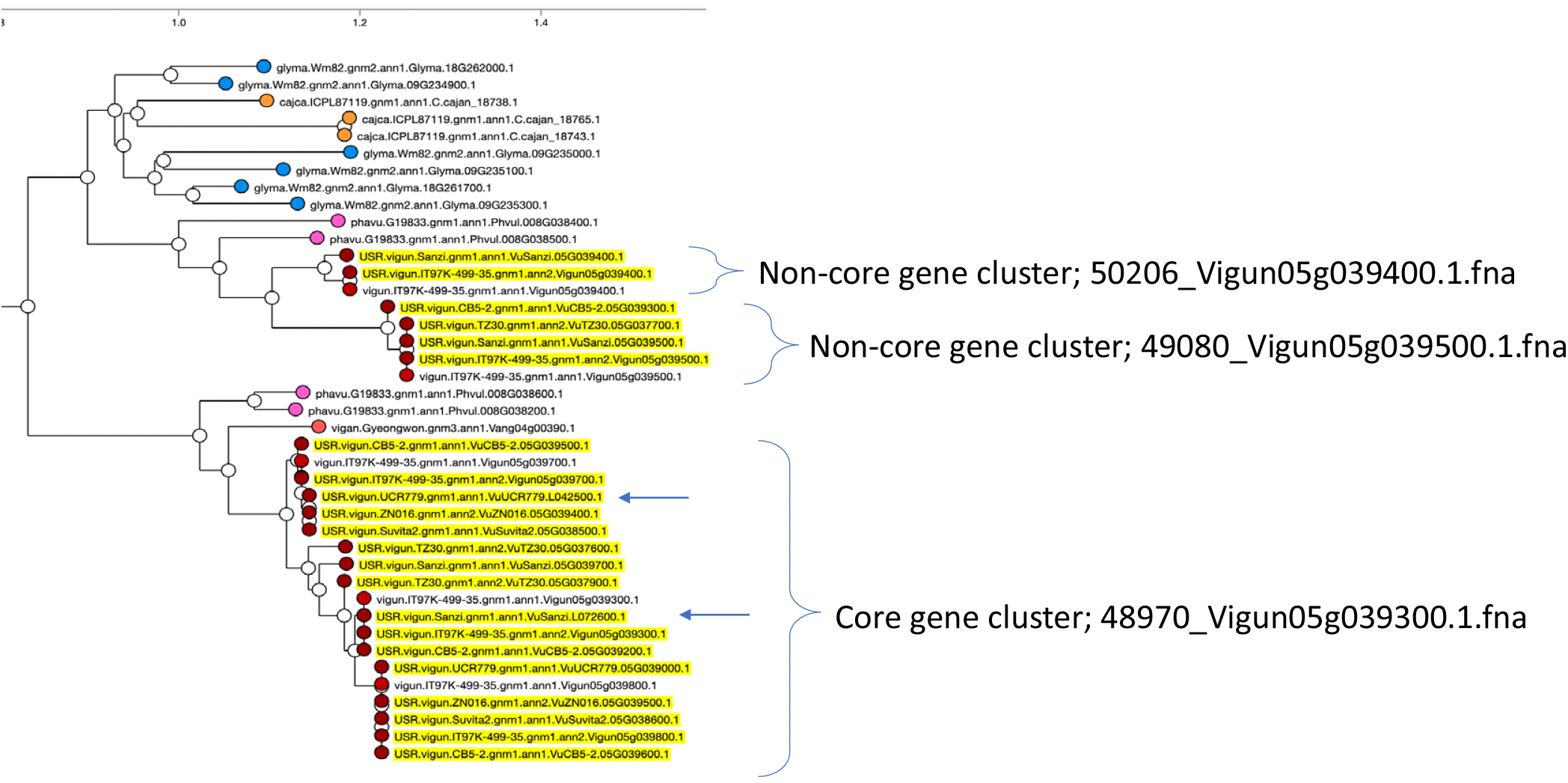
Conservation of gene content within and across species. All cowpea proteins assigned to the family whose members exhibit copy number variation in Figure 6A are shown augmenting a dynamically recomputed gene tree at the Legume Information System, with genes from unanchored contigs not present in the chromosomes aligned in 6A indicated with arrows (https://mines.legumeinfo.org/cowpeamine).

### Pangenome core genes and cross-species synteny

To explore the question of how within-species gene content conservation compares with gene content shared between species in other species and genera, we used the LIS gene family assignments to define homology pairings between all members of each gene family, then used the resulting data to determine collinearity blocks among all pairwise comparisons of the cowpea genomes, as well as to soybean and representative genomes from *Vigna* and *Phaseolus* spp. The counts of genes participating in at least one collinear block were tallied for each genome in each pairwise comparison. As expected, intra-specific comparisons between cowpea accessions yield higher numbers of conserved collinear genes than inter-specific comparisons. On the other hand, there is no appreciable difference in the extent of conserved collinearity when comparing cowpea genomes to other species within the *Vigna* genus versus species from *Phaseolus* or *Glycine* genera (Supplemental Figure S5). Because soybean has an additional whole genome duplication relative to all other species in the comparison, the total number of soybean genes found in collinear blocks is higher than in other comparisons. Comparisons between all species and the *Vigna radiata* version 6 genome (Kang *et al*. 2014) show fewer conserved collinear genes, but this is presumably due to missing data in that assembly, given that all other interspecific comparisons are similar.

## Data Availability Statement

The genome assemblies and annotations described in this manuscript are available from CowpeaPan (https://phytozome-next.jgi.doe.gov/cowpeapan). Raw DNA and RNA sequence data from IT97K-499-35 and whole genome shotgun DNA sequences for 36 diverse cowpea accessions used for SNP discovery in Muñoz-Amatriaín *et al*. (2017) are available from the National Center for Biotechnology Information (NCBI) as SRA accessions SRS3721827, SRP077082, SAMN071606186 through SAMN071606198, SAMN07194302 through SAMN07194309, and SAMN07194882 through SAMN07194909, as stated in Lonardi *et al*. (2019). Raw DNA and RNA sequence data from the six additional accessions providing de novo assemblies in this report, and sequences produced by 10X Genomics from IT97K-49-35, are available as BioProject PRJNA836573 from the National Center for Biotechnology Information (NCBI). More complete annotation files, assemblies and SNPs are also available via the https://drive.google.com/drive/folders/1iQaLW4SLmN2lP7q4k3uovHK3SvsxGbVi?usp=sharing Google shared drive link.

## Acknowledgments

The authors thank Staff Research Associate Yi-Ning Guo (UCR) for RNA preparation; Programmer Steve Wanamaker (UCR) for informatics assistance; Jeffrey Ehlers (Bill & Melinda Gates Foundation), Phillip Roberts (UCR) and Anthony Hall (UCR) for discussions on the accessions for sequencing; Bao Lam Huynh (UCR) and Mitchell Lucas (UCR) for assistance with greenhouse operations for pure seed production; Richard Hayes (Department of Energy Joint Genome Institute, Berkeley, California, USA) for assistance with Phytozome; Samuel Hokin (National Center for Genome Resources, Santa Fe, New Mexico, USA) for data curation for loading data into CowpeaMine; and Bruno Contreras-Moreira (Spanish National Research Council) for helpful discussions on the use of GET_HOMOLOGUES-EST. The University of Minnesota Supercomputing Institute also provided hardware and software support.

## Figure and Table Captions

**Figure 1. Principal component analysis of the UCR Minicore, indicating the accessions selected for sequencing and the subpopulation they belong to.** Accessions in the plot are colored by the result of STRUCTURE for *K*=*6*, as shown in Muñoz-Amatriaín *et al*. (2021).

**Figure 2. The number of genes identified in the pan-genome (pan genes) and core genome (core genes) as new accessions are added.** Green curves are fitted Tettelin functions.

**Figure 3. Gene Ontology (GO) term enrichment analysis.** Significantly enriched GO terms for core (A) and noncore genes (B) are shown for GO-Slim categories belonging to Biological Process, Cellular Component, and Molecular Function aspects (in different colors). -log10 of FDR-adjusted p-values (q-values) are shown on the right of each bar.

**Figure 4. Representation of structural variations (of any size) detected by SyRI from the output of whole-genome pairwise alignments between the seven cowpea accessions.** The black track indicates gene density in the reference genome IT97K-499-35, while the blue track indicates SNP density in the reference genome IT97K-499-35.

**Figure 5. Variant effect predictor (VeP) annotations for SNPs and indels found in the core and noncore genes present in IT97K-499-35.** Values on the y-axis are the absolute number of variants in each variant class.

**Figure 6. Conservation of gene content within and across species.** (A) A region depicting gene content conservation and variability among cowpea genomes and other representative Phaseoleae species. Triangular glyphs represent order and orientation of genes, with color representing gene family memberships. (https://vigna.legumeinfo.org/tools/gcv) (B) All cowpea proteins assigned to the family whose members exhibit copy number variation in (A) are shown augmenting a dynamically recomputed gene tree at the Legume Information System, with genes from unanchored contigs not present in the chromosomes aligned in (A) indicated with arrows (https://mines.legumeinfo.org/cowpeamine).

**Table 1. Summary of assembly statistics, repetitive content, gene content, and BUSCO completeness for the seven genomes.**

**Table 2. Genomic coordinates of all inversions of size > 1 Mbp detected by comparing the seven cowpea genomes pairwise.** IT97K- 499-35 is abbreviated as IT97K.

**Supplemental Figure S1. Gene and repeat density.** Accessions are represented by different shades of red (genes) and blue (repeats). The reference genome of IT97K-499-35 is the longest curve, i.e., the one that extends furthest to the right of each graph.

**Supplemental Figure S2. SNP density (number of SNPs per Mb) using each genome as the “reference.”** (A) IT97K-499-35 (IT97K), centromeres are marked with orange in the innermost circle, (B) CB5-2, (C) Suvita-2, (D) Sanzi, (E) UCR779, (F) ZN016, (G) TZ30.

**Supplemental Figure S3. Output of ALLMAPS for chromosome Vu06 for Suvita-2.** Ten genetic maps were used to orient the five Dovetail contigs in Vu06 (two of which are larger than 1Mb – see Supplemental Table S14). The first four were arbitrarily oriented by ALLMAPS due to low recombination in that region, as shown on the graphs on the right, which plot cM position (y-axis) as a function of physical position (x-axis). In particular, the 8.2 Mb contig represented in gray in the bottom left figure is a region of very low recombination frequency and was likely oriented incorrectly.

**Supplemental Figure S4. Structural variants (of any size) detected by SyRI between any pairs of genomes in this study.** (A) CB5-2 vs Sanzi, (B) CB5-2 vs Suvita-2, (C) CB5-2 vs TZ30, (D) CB5-2 vs UCR779, (E) CB5-2 vs ZN016, (F) IT97K-499-35 vs CB5-2, (G) IT97K-499-35 vs Sanzi, (H) IT97K-499-35 vs Suvita-2, (I) IT97K-499-35 vs TZ30, (J) IT97K-499-35 vs UCR779, (K) IT97K-499-35 vs ZN016, (L) Sanzi vs TZ30, (M) Sanzi vs UCR779, (N) Sanzi vs ZN016, (O) Suvita-2 vs Sanzi, (P) Suvita-2 vs TZ30, (Q) Suvita-2 vs UCR779, (R) Suvita-2 vs ZN016, (S) UCR779 vs TZ30, (T) UCR779 vs ZN016, (U) ZN016 vs TZ30.

**Supplemental Figure S5**. **Macrosynteny views.** (A) Macrosynteny view with blocks representing regions in the IT97K-499-35 reference cowpea genome with conserved gene order relative to each of the genomes shown as tracks below. The region from the microsynteny view of Figure 6A is shown with a vertical gray bar, and the set of chromosomes displayed is restricted to those showing synteny in that region (i.e., the non-cowpea chromosomes have an apparent lack of synteny downstream because of genomic rearrangements that have moved corresponding content to other chromosomes than those shown). Various inversions are seen as blocks with orientations opposing those of their neighboring blocks. Gaps in otherwise syntenic regions indicate regions where gene content diversity outweighs conserved content through presence-absence and copy-number variation. (B) Counts of genes participating in conserved collinear blocks for all pairwise genome comparisons among the cowpea pangenome members and across representative genomes from several genera in the Phaseoleae tribe. Self-comparisons are included to illustrate within-species conservation of duplicated content from ancient whole genome duplication (WGD) events shared by subfamily Faboideae species and the more recent WGD in the *Glycine max* genome.

**Supplemental Table S01. Cowpea accessions used in this work.**

**Supplemental Table S02. Cowpea iSelect SNP positions on each of the seven genome assemblies.** The allele, chromosome and position in the assembled genome are indicated for each accession (columns D-X). For IT97K-499-35 (97K) the orientation of the sequence used for the cowpea iSelect array (Muñoz-Amatriaín et al. 2017) “forward” strand is indicated in column B and the two possible alleles for iSelect assay “forward” strand are in column C.

**Supplemental Table S03. Cowpea iSelect arrray SNP positions and alleles relative to the IT97K-499-35 sequence of Lonardi et al. 2019 (columns B-F,I,J) and to the iSelect “Forward Strand” of Muñoz-Amatriaín et al. 2017 and Muñoz-Amatriaín et al. 2021 (columns G&H).** Other columns indicate reasons for exclusion of data from 2,316 SNPs: blastn alignment ambiguities (columns K-M,S), poor technical performance on array (column N), monomorphic across all DNA samples (column O), excess heterozygote and or no-call (columns P-R).

**Supplemental Table S04. Statistics of the six new assemblies at each step of the Dovetail assembly pipeline.**

**Supplemental Table S05. Putative centromeric region coordinates (all numbers are bp).**

**Supplemental Table S06. Gene annotation statistics.**

**Supplemental Table S07. BUSCO v4 completeness results.**

**Supplemental Table S08. Core and noncore genes identified from sequencing the seven cowpea genomes, tabulated by gene cluster.**

**Supplemental Table S09. Enrichment analysis of GO Terms for core and noncore genes performed in AgriGO v2.** Only significantly enriched GO terms (FDR < 0.05) are shown for the three different ontology aspects.

**Supplemental Table S10. Average diversity at the chromosome (pseudomolecule) level relative to the IT97K-4899-35 assembly.** Values reported are θπ and the standard deviation for “callable regions.”

**Supplemental Table S11. Number of SNPs when considering each accession as the “reference” genome and the resulting union of unique SNPs (merged GVCF) for each accession.**

**Supplemental Table S12. Number of indels of size 1 to 300 bp when considering each accession as the “reference” genome and the union set of all indels (merged GVCF) for each accession.**

**Supplemental Table S13. Genomic coordinates of all structural variants detected via SyRI by comparing the seven cowpea genomes pairwise.** IT97K-499-35 is abbreviated as IT97K.

**Supplemental Table S14. Largest inversions, using each of the Dovetail assemblies as reference.** Left table: ALLMAPS’ orientation of assembled contigs based on markers’ position on the genetic maps (“?” indicates a contig that was arbitrarily oriented). Right tables: Large (>1Mb) inversions detected by SyRI, and whether they are within an oriented ALLMAPS contig.

**Supplemental Table S15. Summary of nucleotide sequence variants in core and noncore genes with potential consequences on coding sequence as identified by Variant Effect Predictor (VeP).** SNPs and indels were analyzed separately. These values are shown in Figure 5. Predictions are based on annotations from the IT97K-499-35 genome assembly.

## Literature Cited

Aggarwal VD, Muleba N, Drabo I, Souma J, Mbewe M (1984) Inheritance of *Striga gesnerioides* resistance in cowpea. In: Proceedings of 3rd International Symposium on Parasitic Weeds (Parker C, Musselman LJ, Polhill RM, Wilson AK eds.) ICARDA, Aleppo, Syria, pp. 143–147

Bayer PE, Golicz AA, Scheben A, Batley J, Edwards D (2020) Plant pan-genomes are the new reference. Nature Plants 6:914–920.

Boukar O, Belko N, Chamarthi S, Togola A, Batieno J, Owusu E, Haruna M, Diallo S, Umar ML, Olufajo O, Fatokun C (2019) Cowpea (*Vigna unguiculata*): Genetics, genomics and breeding. Plant Breeding 138:415–424, doi.org/10.1111/pbr.12589

Boukar O, Bhattacharjee R, Fatokun C, Kumar PL, Gueye B (2013) Cowpea. In: Genetic and Genomic Resources of Grain Legume Improvement (Singh M, Upadhyaya HD, Bisht IS eds.), Chapter 6, pp. 137–156, Elsevier, London, UK. ISBN 978-0-12-397935-3

Cao J, Schneeberger K, Ossowski S, Günther T, Bender S, Fitz, J, Koenig D, Lanz C, Stegle O, Lippert C, Wang X, Ott F, Müller J, Alonso-Blanco C, Borgwardt K, Schmid KJ, Weigel D (2011) Whole-genome sequencing of multiple *Arabidopsis thaliana* populations. Nature Genetics 43:956–963, doi.org/10.1038/ng.911

Chapman JA, Ho I, Sunkara S, Luo S, Schroth GP, Rokhsar DS (2011) Meraculous: *de novo* genome assembly with short paired-end reads. PLOS ONE 6:e23501

Cleary A, Farmer A (2018) Genome Context Viewer: visual exploration of multiple annotated genomes using microsynteny. Bioinformatics 34:1562–1564, doi.org/10.1093/bioinformatics/btx757

Contreras-Moreira B, Cantalapiedra CP, Garcia Pereira MJ, Gordon S, Vogel JP, Igartua E, Casas AM, Vinuesa P (2017) Analysis of plant pan-genomes and transcriptomes with GET_HOMOLOGUES-EST, a clustering solution for sequences of the same species. Frontiers in Plant Science 8:184, doi.org/10.3389/fpls.2017.00184

Corbett-Detig RB, Hartl DL, Sackton TB (2015) Natural selection constrains neutral diversity across a wide range of species. PLOS Biology 13:e1002112, doi: 10.1371/journal.pbio.1002112

Darling AE, Mau B, Perna NT (2010) progressiveMauve: multiple genome alignment with gene gain, loss and rearrangement. PLOS ONE 5:e11147, doi.org/10.1371/journal.pone.0011147

Dash S, Campbell JD, Cannon EK, Cleary AM, Huang W, Kalberer SR, Karingula V, Rice AG, Singh J, Umale PE, Weeks NT, Wilkey AP, Farmer AD, Cannon SB. (2016) Legume information system (LegumeInfo.org): a key component of a set of federated data resources for the legume family. Nucleic Acids Research 44:D1181–1188, doi: 10.1093/nar/gkv1159

de Mooy BE (1985) Germplasm evaluation of Botswana cowpea (Vigna unguiculata [L.] Walp.) landraces. M.S. Thesis, Michigan State University, Lansing, Michigan, USA.

Eddy SR (2011) Accelerated Profile HMM Searches. PLoS Computational Biology 7:e1002195, doi: 10.1371/journal.pcbi.1002195.

Ehlers JD, Fery RL, Hall AE (2002) Cowpea breeding in the USA: new varieties and improved germplasm. *In* Challenges and Opportunities for Enhancing Sustainable Cowpea Production. Proceedings of the World Cowpea Conference III held at the International Institute of Tropical Agriculture (IITA), Ibadan, Nigeria, 4-8 September 2000, (Eds. Fatokun CS, Tarawali SA, Singh BB, Kormawa PM, Tamò M), Chapter 1.6, pp. 62–77, IITA, Ibadan, Nigeria, ISBN 978-131-190-8

Goel M, Schneeberger K (2022) plotsr: visualizing structural similarities and rearrangements between multiple genomes. Bioinformatics 38:2922–2926, doi.org/10.1093/bioinformatics/btac196

Goel M, Sun H, Jiao W-B, Schneeberger K (2019) SyRI: finding genomic rearrangements and local sequence differences from whole-genome assemblies. Genome Biology 20:277, doi:10.1186/s13059-019-1911-0

Golicz AA, Bayer PE, Bhalla PL, Batley J, Edwards D (2020) Pangenomics comes of age: from bacteria to plant and animal applications. Trends in Genetics 36:132–145

Golicz AA, Bayer PE, Barker GC, Edger PP, Kim HR, Martinez PA, Chan CKK, Severn-Ellis A, McCombie R, Parkin IAP, Paterson AH, Pires JC, Sharpe AG, Tang H, Teakle GR, Town CD, Bately J, Edwards D (2016) The pangenome of an agronomically important crop plant *Brassica oleracea*. Nature Communications 7:13390, doi.org/10.1038/ncomms13390

Gordon SP, Contreras-Moreira B, Woods DP, Des Marais DL, Burgess D, Shu S, Stritt C, Roulin AC, Schackwitz W, Tyler L, Martin J, Lipzen A, Dochy N, Phillips J, Barry K, Geuten K, Budak H, Juenger TE, Amasino R, Caicedo AL, Goodstein D, Davidson P, Mur LAJ, Figueroa M, Freeling M, Catalan P, Vogel JP (2017) Extensive gene content variation in the *Brachypodium distachyon* pan-genome correlates with population structure. Nature Communications 8:2184, doi.org/10.1038/s41467-017-02292-8

Haas BJ, Delcher AL, Mount SM, Wortman JR, Smith Jr RK, Hannick LI, Maiti R, Ronning CM, Rusch DB, Town CD, Salzberg SL, White O (2003) Improving the *Arabidopsis* genome annotation using maximal transcript alignment assemblies. Nucleic Acids Research 31:5654–5666, http://nar.oupjournals.org/cgi/content/full/31/19/5654

Hastings PJ, Lupski JR, Rosenberg SM, Ira G (2009) Mechanisms of change in gene copy number. Nature Review Genetics 10:551–564, doi: 10.1038/nrg2593. PMID: 19597530; PMCID: PMC2864001.

Hall AE (2004) Breeding for adaptation to drought and heat in cowpea. European Journal of Agronomy 21:447–454

Herniter IA, Muñoz-Amatriaín M, Close TJ (2020) Genetic, textual, and archeological evidence of the historical global spread of cowpea (*Vigna unguiculata* [L.] Walp.). Legume Science 3:57

Herniter IA, Muñoz-Amatriaín M, Lo S, Guo Y-N, Close TJ (2018) Identification of candidate genes controlling black seed coat and pod tip color in G3 8:3347–3355

Hirsch CN, Foerster JM, Johnson JM, Sekhon RS, Muttoni G, Vaillancourt B, Peñagaricano F, Lindquist E, Pedraza MA, Barry K, de Leon N, Kaeppler SM, Buell CR (2014) Insights into the maize pan-genome and pan-transcriptome. The Plant Cell 26:121–135

Iwata-Otsubo A, Lin J-Y, Gill N, Jackson SA (2016) Highly distinct chromosome structures in cowpea (*Vigna unguiculata*), as revealed by molecular cytogenetic analysis. Chromosome Research 24:197–216

Kalderimis A, Lyne R, Butano D, Contrino S, Lyne M, Heimbach J, Hu F, Smith R, Stĕpán R, Sullivan J, Micklem G (2014) InterMine: extensive web services for modern biology. Nucleic Acids Research 42:W468–472, doi: 10.1093/nar/gku301

Kang YJ, Kim SK, Kim MY, Lestari P, Kim KH, Ha BK, Jun TH, Hwang WJ, Lee T, Lee J, Shim S, Yoon MY, Jang YE, Han KS, Taeprayoon P, Yoon N, Somta P, Tanya P, Kim KS, Gwag JG, Moon JK, Lee YH, Park BS, Bombarely A, Doyle JJ, Jackson SA, Schafleitner R, Srinives P, Varshney RK, Lee SH (2014) Genome sequence of mungbean and insights into evolution within *Vigna* species. Nature Communications 5:5443, doi: 10.1038/ncomms6443

Kent WJ (2002) BLAT - the BLAST-like alignment tool. Genome Research 12:656–64

Kirkpatrick M (2010) How and why chromosome inversions evolve. PLOS Biology 8:e1000501

Kirkpatrick M, Barton N (2006) Chromosome inversions, local adaptation and speciation. Genetics 173:419–434

Korunes KL, Samuk K (2021) PIXY: Unbiased estimation of nucleotide diversity and divergence in the presence of missing data. Molecular Ecology Resources 21:1359–1368

Leffler EM, Bullaughey K, Matute DR, Meyer WK, Ségurel L, Venkat A, Andolfatto P, Przeworski M (2012) Revisiting an old riddle: what determines genetic diversity levels within species. PLOS Biology 10:e1001388

Lei L, Goltsman E, Goodstein D, Wu GA, Rokhsar DS, Vogel JP (2021) Plant pan-genomics comes of age. Annual Reviews of Plant Biology 72:411–435

Li H (2018) Minimap2: pairwise alignment for nucleotide sequences. Bioinformatics 34:3094–3100

Li H, Durbin R (2009) Fast and accurate short read alignment with Burrows-Wheeler transform. Bioinformatics 25:1754–1760

Li H, Handsaker B, Wysoker A, Fennell T, Ruan J, Homer N, Marth G, Abecasis G, Durbin R (2009) 1000 Genome Project Data Processing Subgroup, The Sequence Alignment/Map format and SAMtools. Bioinformatics 25:2078–2079

Li YH, Zhou G, Ma J, Jiang W, Jin LG, Zhang Z, Guo Y, Zhang J, Sui Y, Zheng L, Zhang SS, Zuo Q, Shi XH, Li YF, Zhang WK, Hu Y, Kong G, Hong HL, Tan B, Song J, Liu ZX, Wang Y, Ruan H, Yeung CKL, Liu J, Wang H, Zhang LJ, Guan RX, Wang KJ, Li WB, Chen SY, Chang RZ, Jiang Z, Jackson SA, Li R, Qiu LJ (2014) *De novo* assembly of soybean wild relatives for pan-genome analysis of diversity and agronomic traits. Nature Biotechnology 32:1045–1052, doi.org/10.1038/nbt.2979

Liang Q, Lonardi S (2021) Reference-agnostic representation and visualization of pan-genomes. BMC Bioinformatics 22:502

Lonardi S, Muñoz-Amatriaín M, Liang Q, Shu S, Wanamaker SI, Lo S, Tanskanen J, Schulman AH, Zhu T, Luo MC, Alhakami H, Ounit R, Hasan AM, Verdier J, Roberts PA, Santos JRP, Ndeve A, Doležel J, Vrána J, Hokin SA, Farmer AD, Cannon SB, Close TJ (2019) The genome of cowpea (*Vigna unguiculata* [L.] Walp.). The Plant Journal 98:767–782

Mackie WW (1946) Blackeye beans in California. Bulletin 696, University of California Agricultural Experiment Station

McLaren W, Gil L, Hunt SE, Riat HS, Ritchie GRS, Thormann A, Flicek P, Cunningham F (2016) The Ensembl variant effect predictor. Genome Biology 17:122

Miller AJ, Gross BL (2011) From forest to field: perennial fruit crop domestication. American Journal of Botany 98:1389–1414

Moghaddam SM, Oladzad A, Koh C, Ramsay L, Hart JP, Mamidi S, Hoopes G, Sreedasyam A, Wiersma A, Zhao D, Grimwood J, Hamilton JP, Jenkins J, Vaillancourt B, Wood JC, Schmutz J, Kagale S, Porch T, Bett KE, Buell CR, McClean PE (2021) The tepary bean genome provides insight into evolution and domestication under heat stress. Nature Communications 12:2638, doi: 10.1038/s41467-021-22858-x

Montenegro JD, Golicz AA, Bayer PE, Hurgobin B, Lee H, Chan C-KK., Visendi P, Lai K, Doležel J, Batley J, Edwards D (2017) The pangenome of hexaploid bread wheat. The Plant Journal 90:1007–1013, doi.org/10.1111/tpj.13515

Morgante M, De Paoli E, Radovic S (2007) Transposable elements and the plant pan-genomes. Current Opinions in Plant Biology 10:149–155

Morrell PL, Gonzales AM, Meyer KKT, Clegg MT (2014) Resequencing data indicate domestication’s modest effect on barley diversity: a cultigen with multiple origins. Journal of Heredity 105:253–264, doi: 10.1093/jhered/est083

Muñoz-Amatriaín M, Eichten SR, Wicker T, Richmond TA, Mascher M, Steuernagel B, Scholz U, Ariyadasa R, Spannagl M, Nussbaumer T, Mayer KF, Taudien S, Platzer M, Jeddeloh JA, Springer NM, Muehlbauer GJ, Stein N (2013) Distribution, functional impact, and origin mechanisms of copy number variation in the barley genome. Genome Biology 14:R58

Muñoz-Amatriaín M, Lo S, Herniter IA, Boukar O, Fatokun C, Carvalho M, Castro I, Guo Y-N, Huynh B-L, Roberts PA, Carnide V, Close TJ (2021) The UCR Minicore: a resource for cowpea research and breeding. Legume Science 3:e95, doi.org/10.1002/leg3

Muñoz-Amatriaín M, Mirebrahim H, Xu P, Wanamaker SI, Luo M, Alhakami H, Alpert M, Atokple I, Batieno BJ, Boukar O, Bozdag S, Cissé N, Drabo I, Ehlers JD, Farmer A, Fatokun C, Gu YQ, Guo Y, Huynh B, Jackson SA, Kusi F, Lawley CT, Lucas MR, Ma Y, Timko MP, Wu J, You F, Barkley NA, Roberts PA, Lonardi S, Close TJ (2017) Genome resources for climate-resilient cowpea, an essential crop for food security. The Plant Journal 89:1042–1054

Onsongo G, Xie H, Griffin TJ, Carlis (2008) Generating GO slim using relational database management systems to support proteomics analysis. 21^st^ IEEE International Symposium on Computer-Based Medical Systems, doi.org/10.1109/CBMS.2008.77

Pedersen BS, Quinlan AR (2018) Mosdepth: quick coverage calculation for genomes and exomes. Bioinformatics 34:867–868, doi.org/10.1093/bioinformatics/btx699

Quinlan AR, Hall IM (2010) BEDtools: a flexible suite of utilities for comparing genomic features. Bioinformatics 26:841–841

Sakai H, Naito K, Ogiso-Tanaka E, Takahashi Y, Iseki K, Muto C, Satou K, Teruya K, Shiroma A, Shimoji M, Hirano T, Itoh T, Kaga A, Tomooka N (2015) The power of single molecule real-time sequencing technology in the de novo assembly of a eukaryotic genome. Scientific Reports 5:16780, doi:10.1038/srep16780

Salamov AA, Solovyev VV (2000) *Ab initio* gene finding in *Drosophila* genomic DNA. Genome Research 10:516–522

Schmid K, Kilian B, Russell J (2018) Barley Domestication, Adaptation and Population Genomics. editors. In: Stein N, Muehlbauer G (eds.) The Barley Genome. Compendium of Plant Genomes. Springer, Cham. pp. 317–336.

Schmutz J, McClean PE, Mamidi S, Wu GA, Cannon SB, Grimwood J, Jenkins J, Shu S, Song Q, Chavarro C, Torres-Torres M, Geffroy V, Moghaddam SM, Gao D, Abernathy B, Barry K, Blair M, Brick MA, Chovatia M, Gepts P, Goodstein DM, Gonzales M, Hellsten U, Hyten DL, Jia G, Kelly JD, Kudrna D, Lee R, Richard MM, Miklas PN, Osorno JM, Rodrigues J, Thareau V, Urrea CA, Wang M, Yu Y, Zhang M, Wing RA, Cregan PB, Rokhsar DS, Jackson SA (2014) A reference genome for common bean and genome-wide analysis of dual domestications. Nature Genetics 46:707–713, doi:10.1038/ng.3008

Shu S, Goodstein D, Rokhsar D (2013) PERTRAN: genome-guided RNA-seq read assembler. https://www.osti.gov/biblio/1241180

Shu S, Rokhsar D, Goodstein D, Hayes D, Mitros T (2014) JGI plant genomics gene annotation pipeline. Lawrence Berkeley National Laboratory, Berkeley, CA, USA

Singh BB, Olufajo OO, Ishiyaku MF, Adeleke RA, Ajeigbe HA, Mohammed SG (2006) Registration of six improved germplasm lines of cowpea with combined resistance to *Striga gesnerioides* and *Alectra vogelii*. Crop Science 46:2332–2333

Springer NM, Ying K, Fu Y, Ji T, Yeh CT, Jia Y, Wu W, Richmond T, Kitzman J, Rosenbaum H, Iniguez AL, Barbazuk WB, Jeddeloh JA, Nettleton D, Schnable PS (2009) Maize inbreds exhibit high levels of copy number variation (CNV) and presence/absence variation (PAV) in genome content. PLoS Genetics 5:e1000734

Stai JS, Yadav A, Sinou C, Bruneau A, Doyle JJ, Fernández-Baca D, Cannon SB (2019) Cercis: a non-polyploid genomic relic within the generally polyploid legume family. Frontiers in Plant Science 10:345, doi:10.3389/fpls.2019.00345

Tajima F (1983) Evolutionary relationship of DNA sequences in finite populations. Genetics 105:437–460

Tang H, Zhang X, Miao C, Zhang J, Ming R, Schnable JC, Schnable PS, Lyons E, Lu J (2015) ALLMAPS: robust scaffold ordering based on multiple maps. Genome Biology 16:3, doi:10.1186/s13059-014-0573-1

Tettelin H, Masignani V, Cieslewicz MJ, Donati C, Medini D, Ward NL, Angiuoli SV, Crabtree J, Jones AL, Durkin AS, DeBoy RT, Davidsen TM, Mora M, Scarselli M, Immaculada Margarit y Ros I, Peterson JD, Hauser CR, Sundaram JP, Nelson WC, Madupu R, Brinkac LM, Dodson RJ, Rosovitz MJ, Sullivan SA, Daugherty SC, Haft DH, Selengut J, Gwinn ML, Zhou L, Zafar N, Khouri H, Radune D, Dimitrov G, Watkins K, O’Connor KJB, Smith S, Utterback TR, White O, Rubens CE, Grandi G, Madoff LC, Kasper DL, Telford JL, Wessels MR, Rappuoli R, Fraser CM (2005) Genome analysis of multiple pathogenic isolates of *Streptococcus agalactiae:* implications for the microbial “pan-genome”. Proc. Natl. Acad. Sci. USA 102:13950–13955, doi:10.1073/pnas.0506758102

Tian T, Liu Y, Yan H, You Q, Yi X, Du Z, Xu W, Su Z (2017) agriGO v2.0: a GO analysis toolkit for the agricultural community, 2017 update. Nucleic Acids Research 45:W122–W129, doi: 10.1093/nar/gkx382

Tittes S, Lorant A, McGinty S, Doebley JF, Holland JB, de Jesus Sánchez-González, Seetharam A, Tenaillon M, Ross-Ibarra J (2021) Not so local: the population genetics of convergent adaptation in maize and teosinte. bioRxiv, doi: 10.1101/2021.09.09.459637

Torkamaneh D, Lemay M-A, Belzile F (2021) The pan-genome of the cultivated soybean (PanSoy) reveals an extraordinarily conserved gene content. Plant Biotechnology Journal 19:1852–1862, https://doi.org/10.1111/pbi.13600

Valliyodan B, Cannon SB, Bayer PE, Shu S, Brown AV, Ren L, Jenkins J, Chung CY, Chan TF, Daum CG, Plott C, Hastie A, Baruch K, Barry KW, Huang W, Patil G, Varshney RK, Hu H, Batley J, Yuan Y, Song Q, Stupar RM, Goodstein DM, Stacey G, Lam HM, Jackson SA, Schmutz J, Grimwood J, Edwards D, Nguyen HT (2019) Construction and comparison of three reference-quality genome assemblies for soybean. The Plant Journal 100:1066–1082, doi: 10.1111/tpj.14500

Wang Y, Tang H, Debarry JD, Tan X, Li J, Wang X, Lee TH, Jin H, Marler B, Guo H, Kissinger JC, Paterson AH (2012) MCScanX: a toolkit for detection and evolutionary analysis of gene synteny and collinearity. Nucleic Acids Research 40:e49, doi: 10.1093/nar/gkr1293

Xu P, Wu X, Muñoz-Amatriaín M, Wang B, Wu X, Hu Y, Huynh BL, Close TJ, Roberts PA, Zhou W, Lu Z, Li G (2017) Genomic regions, cellular components and gene regulatory basis underlying pod length variations in cowpea (*V. unguiculata* L. Walp). Plant Biotechnology Journal 15:547–557, doi:10.1111/pbi.12639

Yeh R-F, Lim LP, Burge CB (2001) Computational inference of homologous gene structures in the human genome. Genome Research 11:803–816

Yu G, Wang L, Han Y, He Q (2012). clusterProfiler: an R package for comparing biological themes among gene clusters. OMICS: A Journal of Integrative Biology 16:284–287, doi:10.1089/omi.2011.0118

