## Supplemental Figure S1 for "A view of the pan-genome of domesticated cowpea (*Vigna unguiculata* [L.] Walp.)"

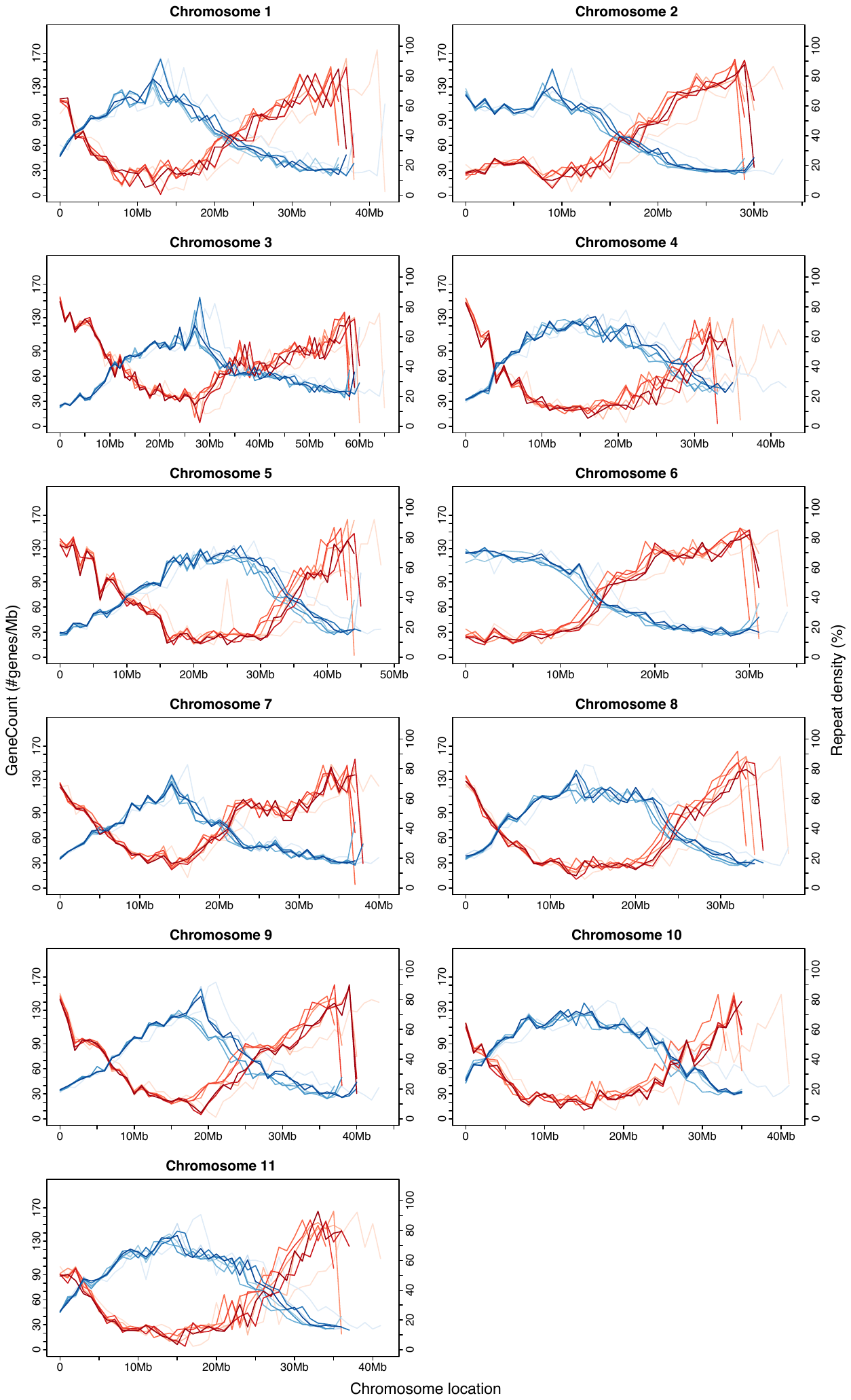

**Supplemental Figure S1.** **Gene and repeat density.** Accessions are represented by different shades of red (genes) and blue (repeats). The reference genome of IT97K-499-35 is the longest curve, i.e., the one that extends furthest to the right of each graph.
