## Supplemental Figure S2 for "A view of the pan-genome of domesticated cowpea (*Vigna unguiculata* [L.] Walp.)"

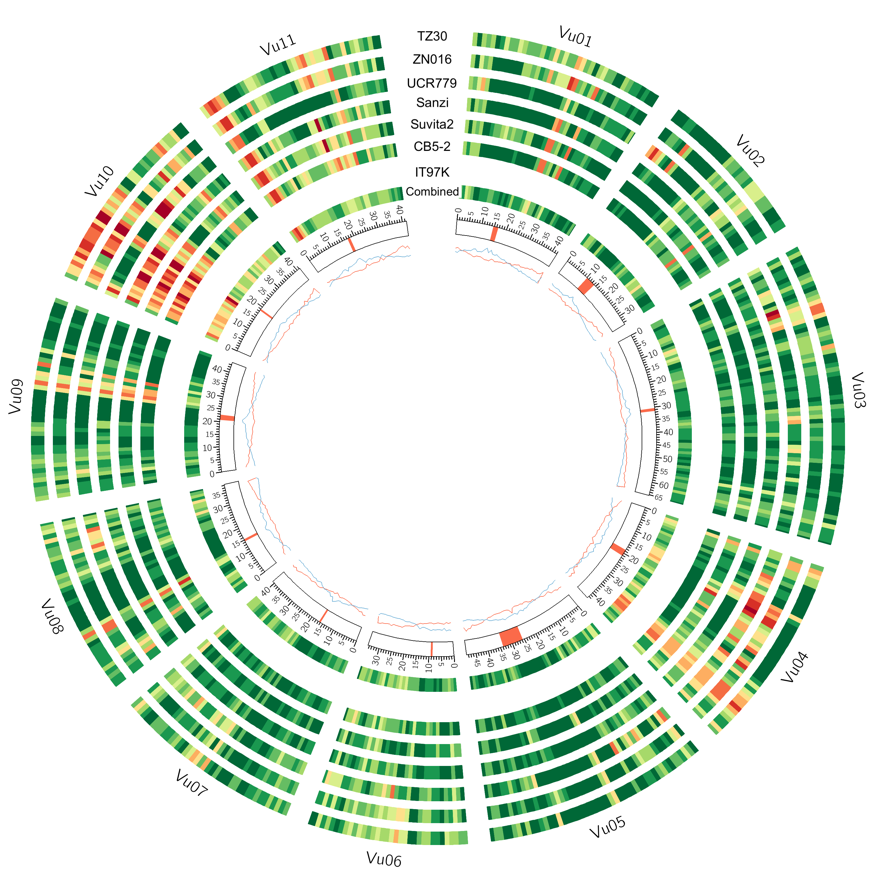

**Supplemental Figure S2A. SNP density (number of SNPs per Mb) using IT97K-499-35 as the “reference”.** Centromeres for IT97K-499-35 are marked with orange in the innermost circle.

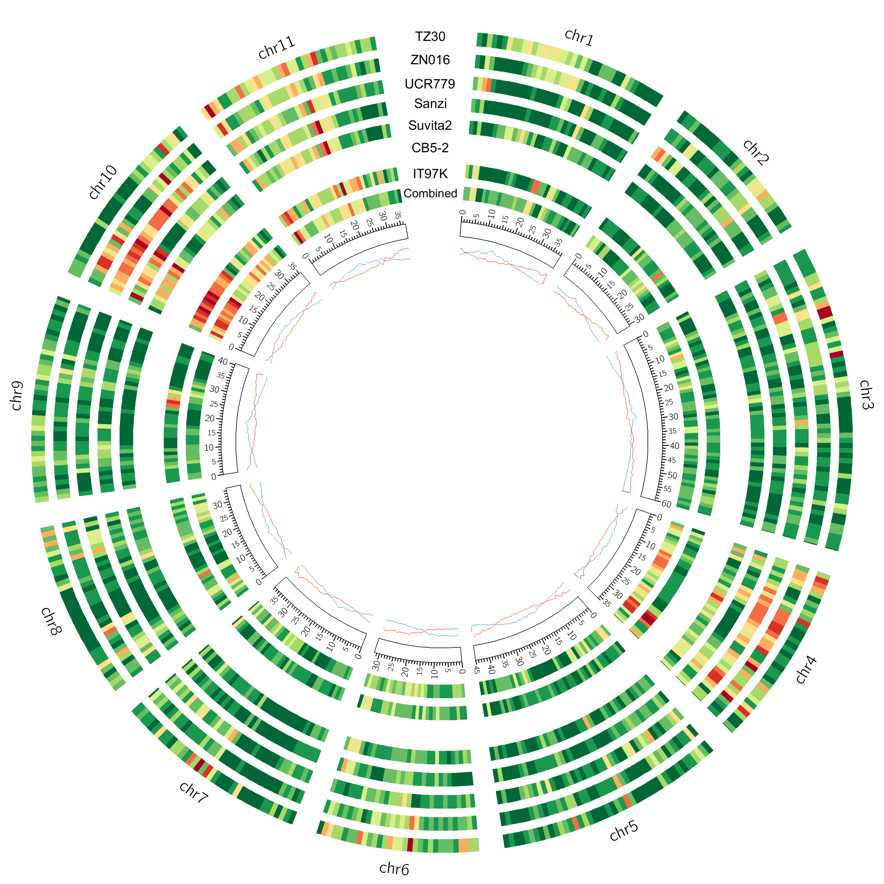

**Supplemental Figure S2B.** **SNP density (number of SNPs per Mb) using CB5-2 as the “reference”.**

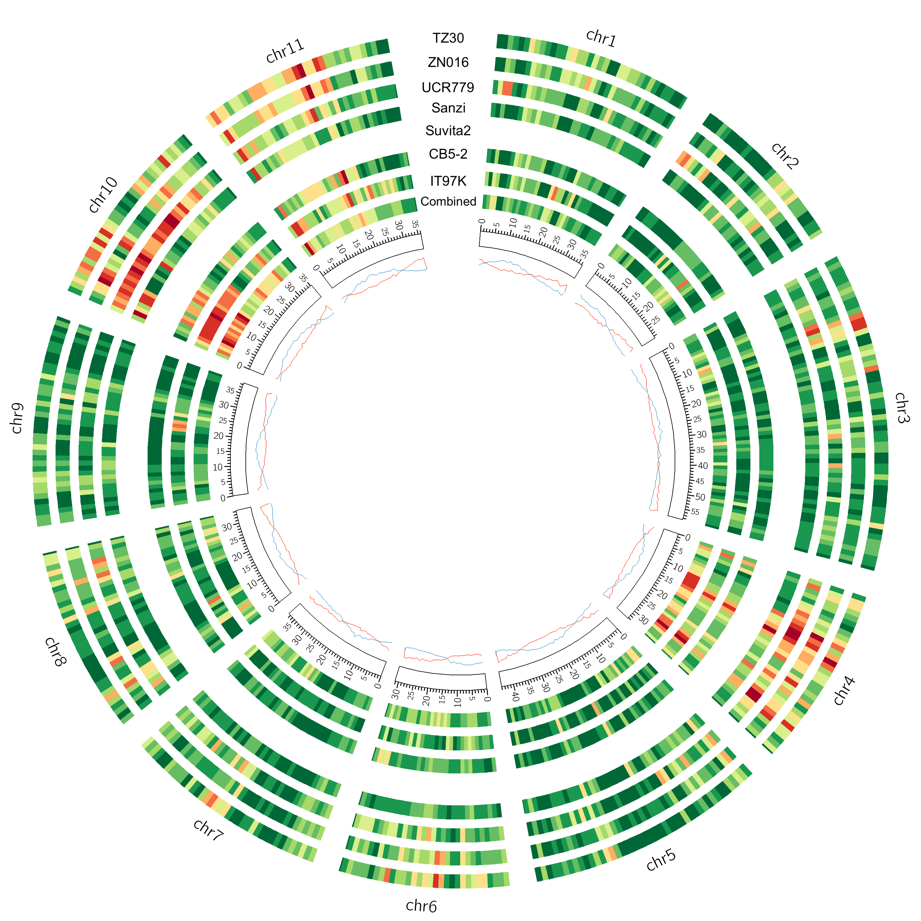

**Supplemental Figure S2C.** **SNP density (number of SNPs per Mb) using Suvita-2 as the “reference”.**

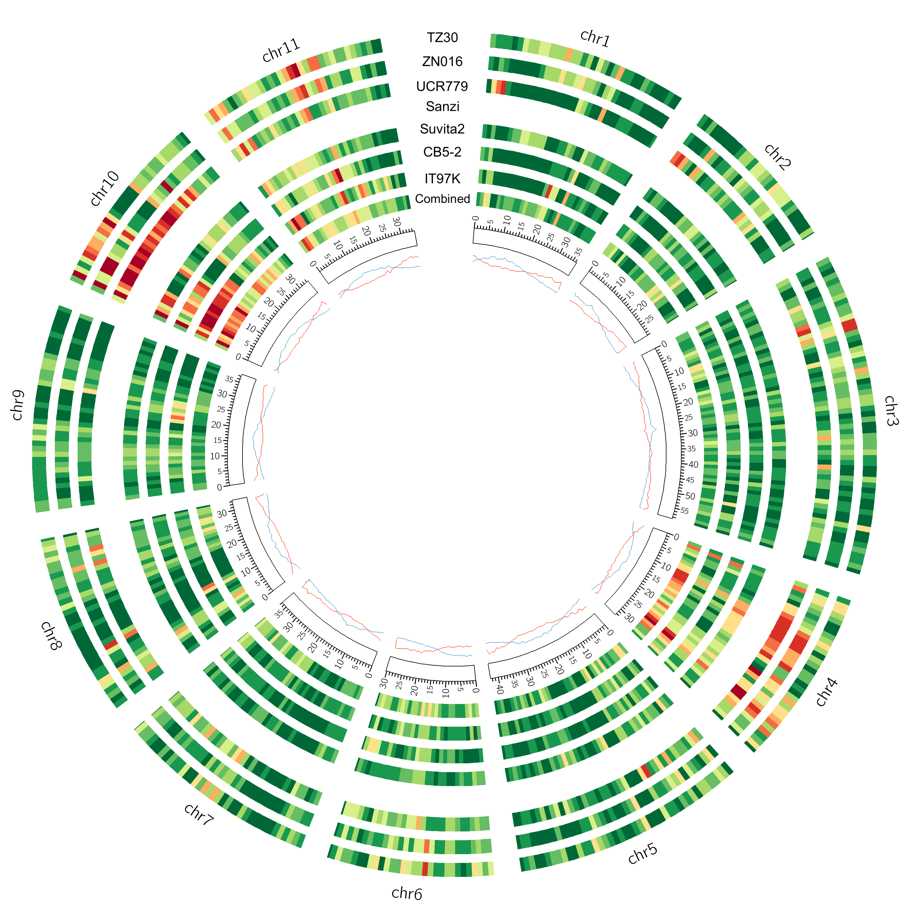

**Supplemental Figure S2D.** **SNP density (number of SNPs per Mb) using Sanzi as the “reference”.**

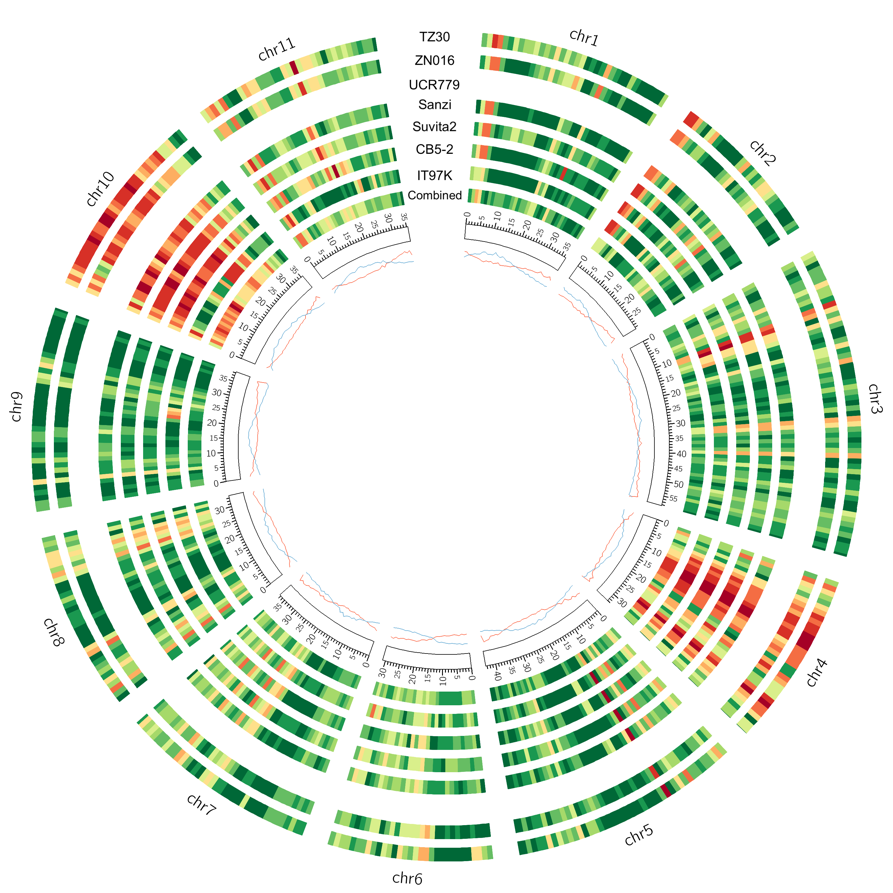

**Supplemental Figure S2E**. **SNP density (number of SNPs per Mb) using UCR779 as the “reference”.**

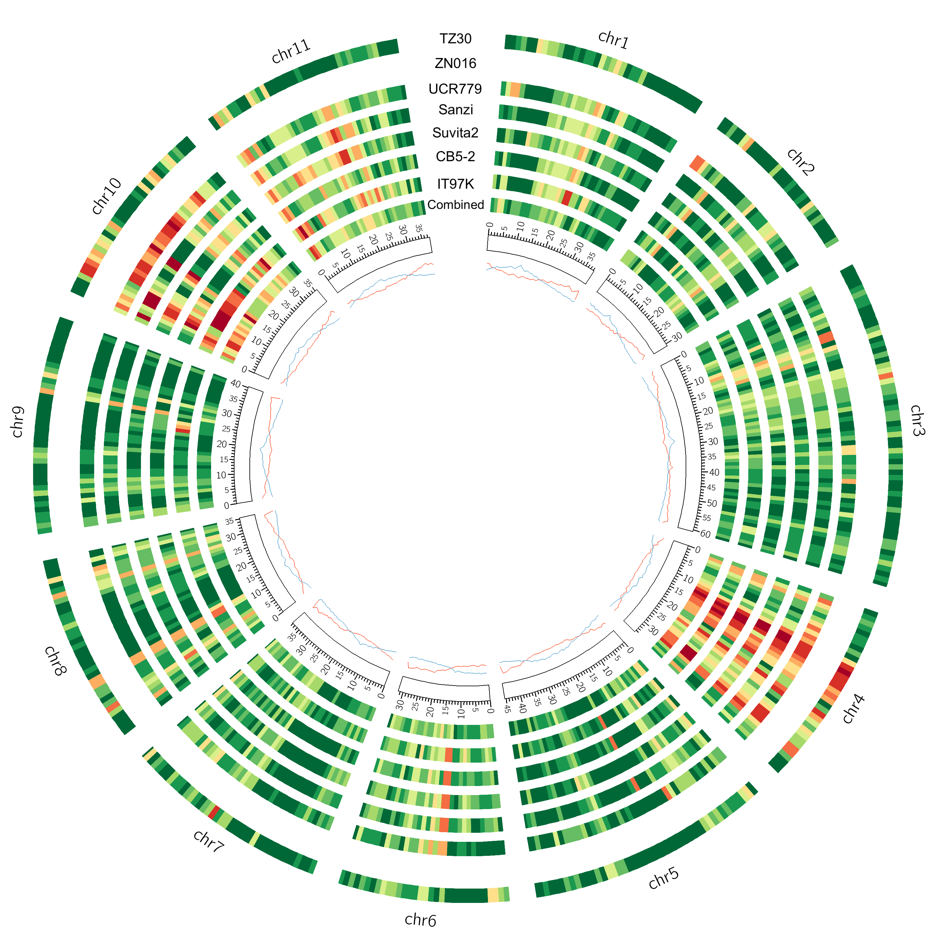

**Supplemental Figure S2F**. **SNP density (number of SNPs per Mb) using ZN016 as the “reference”.**

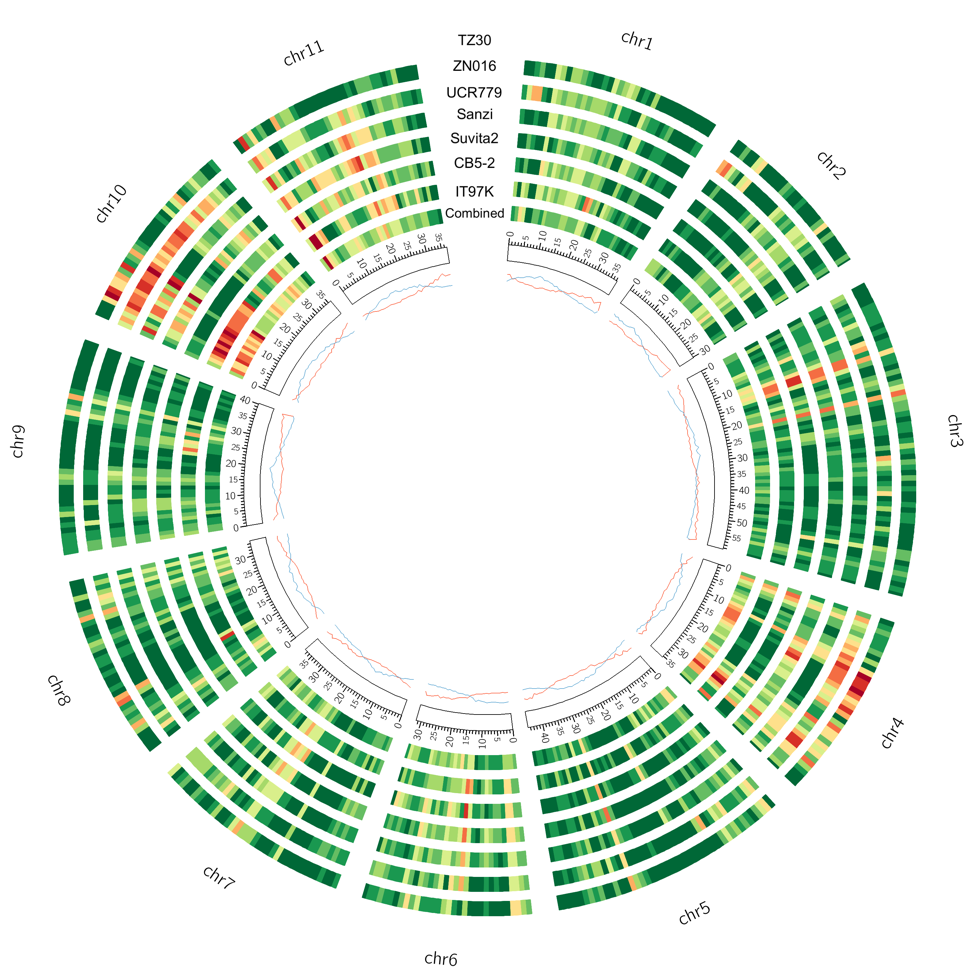

**Supplemental Figure S2G.** **SNP density (number of SNPs per Mb) using TZ30 as the “reference”.**
