## Supplemental Figure S3 for "A view of the pan-genome of domesticated cowpea (*Vigna unguiculata* [L.] Walp.)"

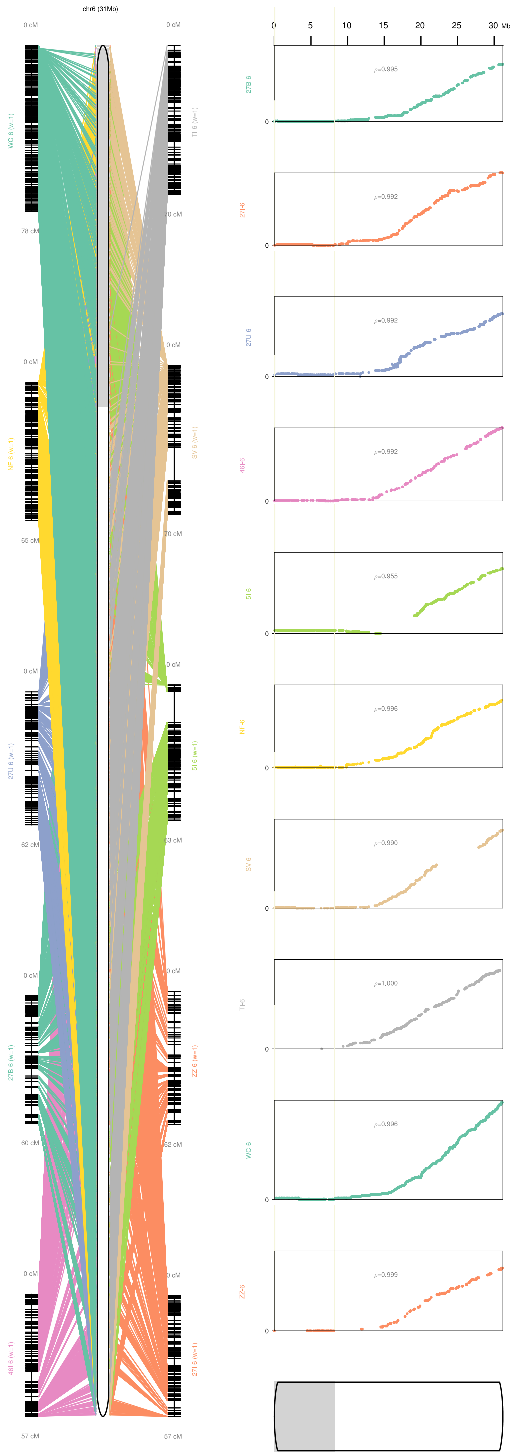


**Supplemental Figure S3. Output of ALLMAPS for chromosome Vu06 for Suvita-2.** Ten genetic maps were used to orient the five Dovetail contigs in Vu06 (two of which are larger than 1Mb – see Supplemental Table S14). The first four were arbitrarily oriented by ALLMAPS due to low recombination in that region, as shown on the graphs on the right, which plot cM position (y-axis) as a function of physical position (x-axis). In particular, the 8.2 Mb contig represented in gray in the bottom left figure, is a region of very low recombination frequency and was likely oriented incorrectly.
