## Supplemental Figure S4 for "A view of the pan-genome of domesticated cowpea (*Vigna unguiculata* [L.] Walp.)"

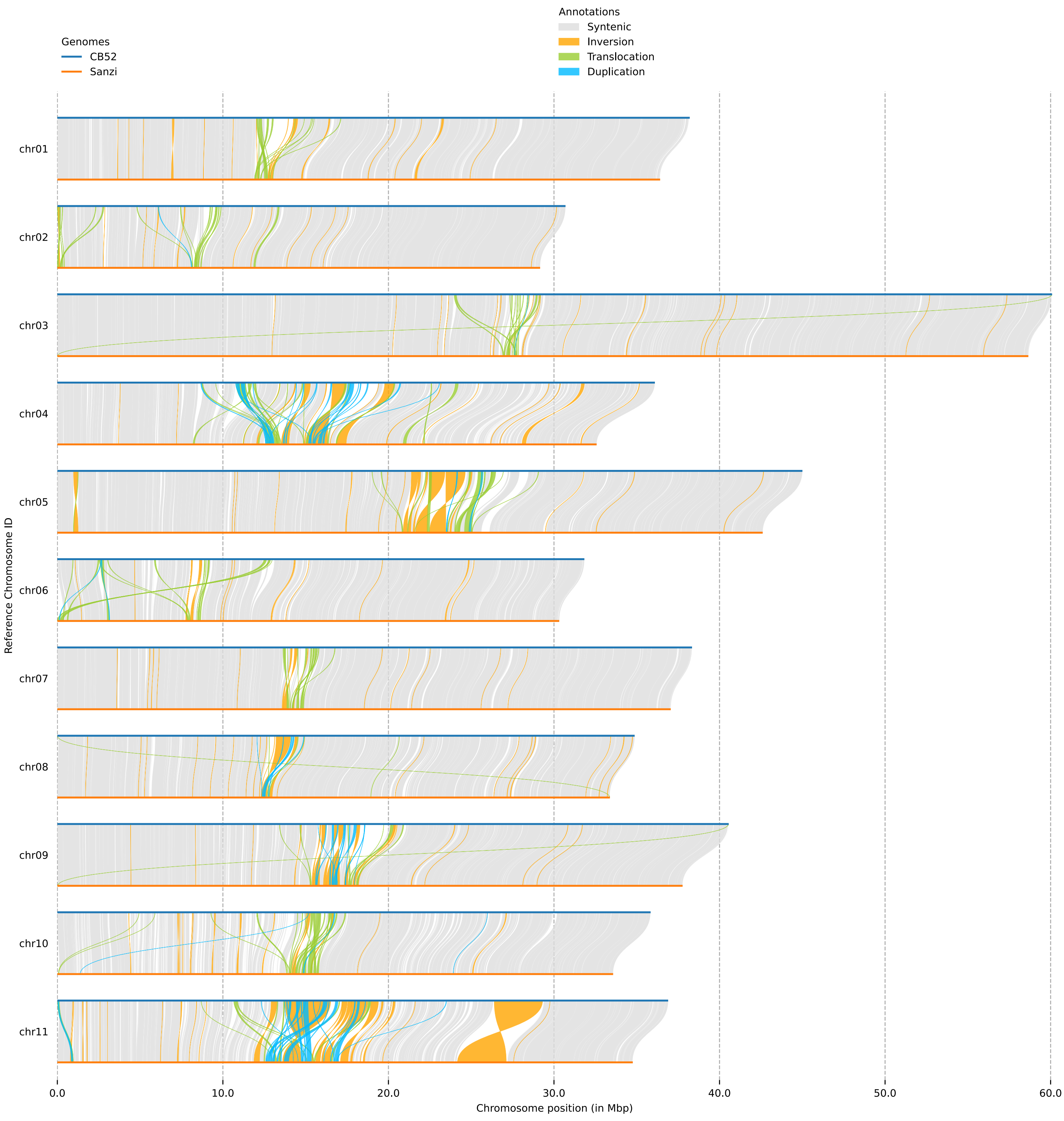


**Supplemental Figure S4A.** Structural variants (of any size) detected by SyRI between the genomes of CB5-2 and Sanzi.

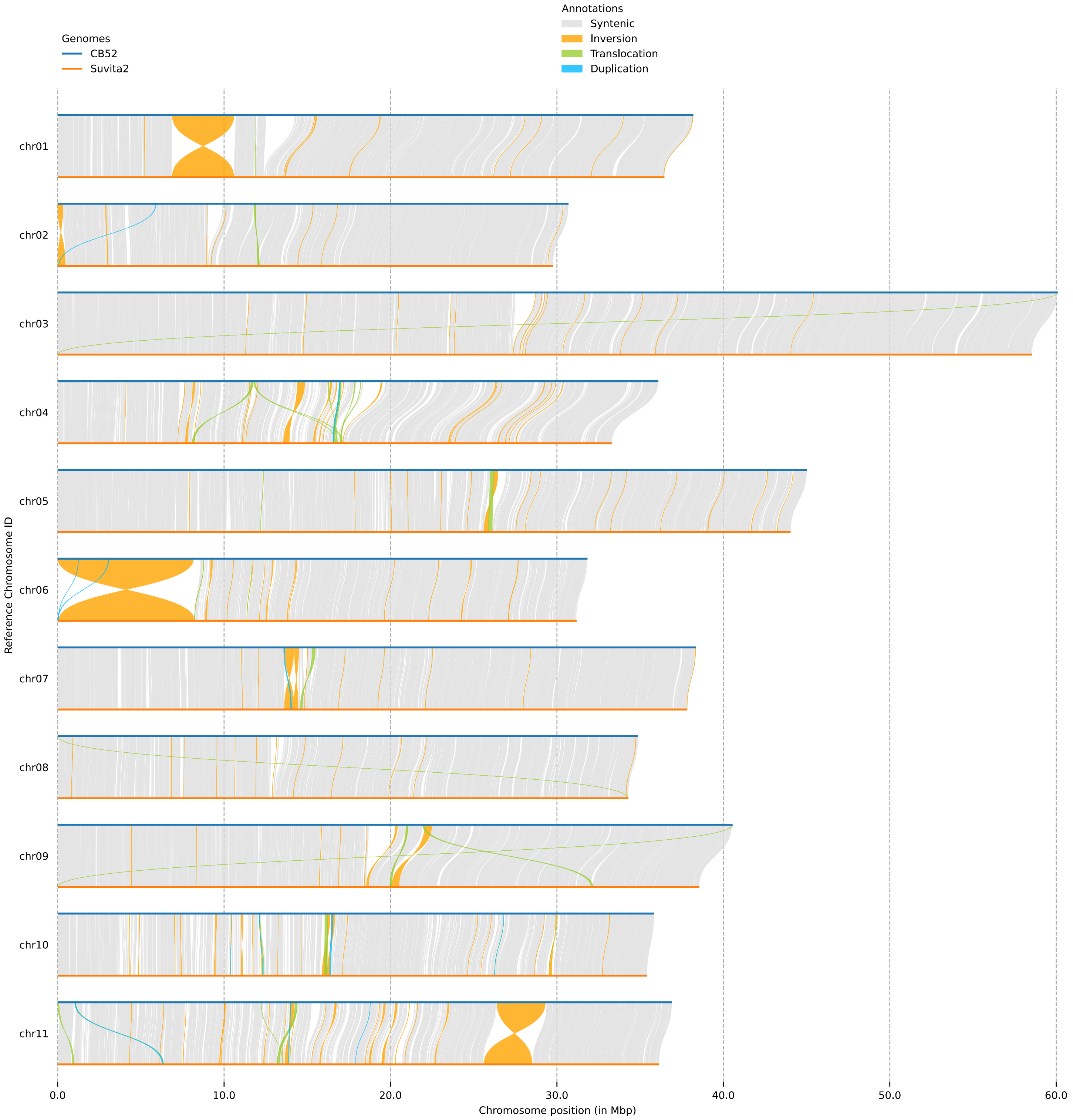


**Supplemental Figure S4B.** Structural variants (of any size) detected by SyRI between the genomes of CB5-2 and Suvita-2.

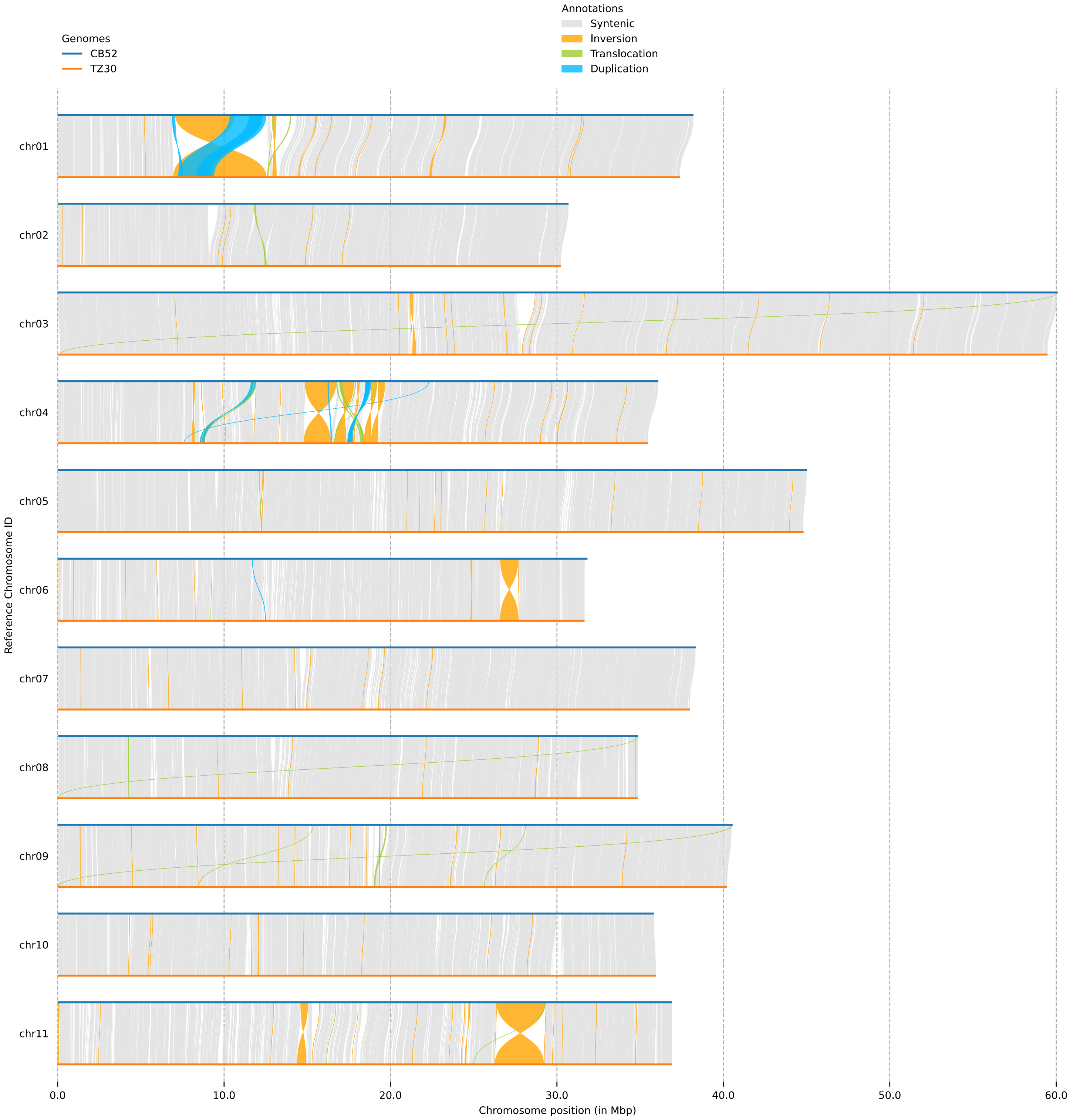


**Supplemental Figure S4C.** Structural variants (of any size) detected by SyRI between the genomes of CB5-2 and TZ30.

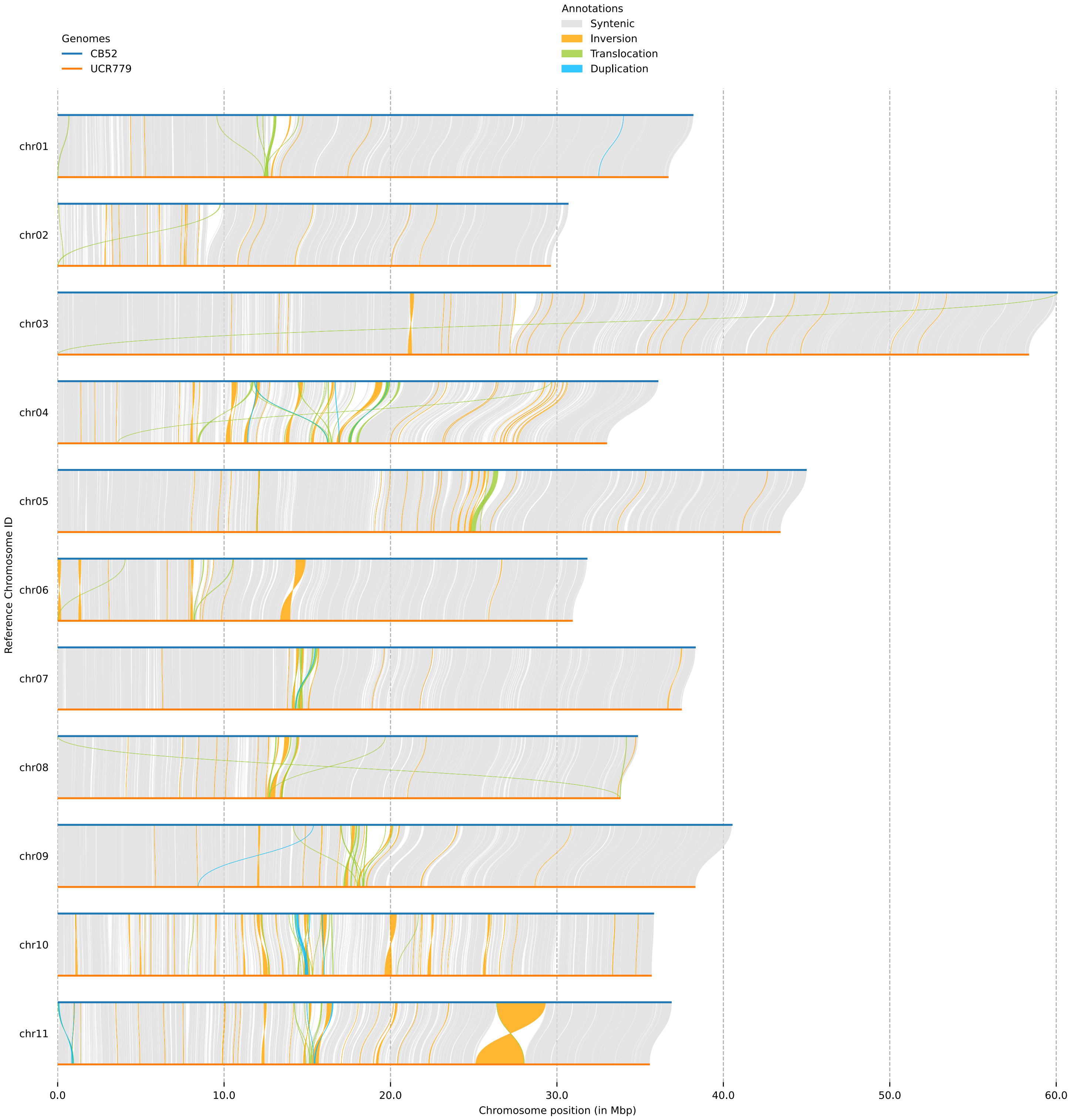


**Supplemental Figure S4D.** Structural variants (of any size) detected by SyRI between the genomes of CB5-2 and UCR779.

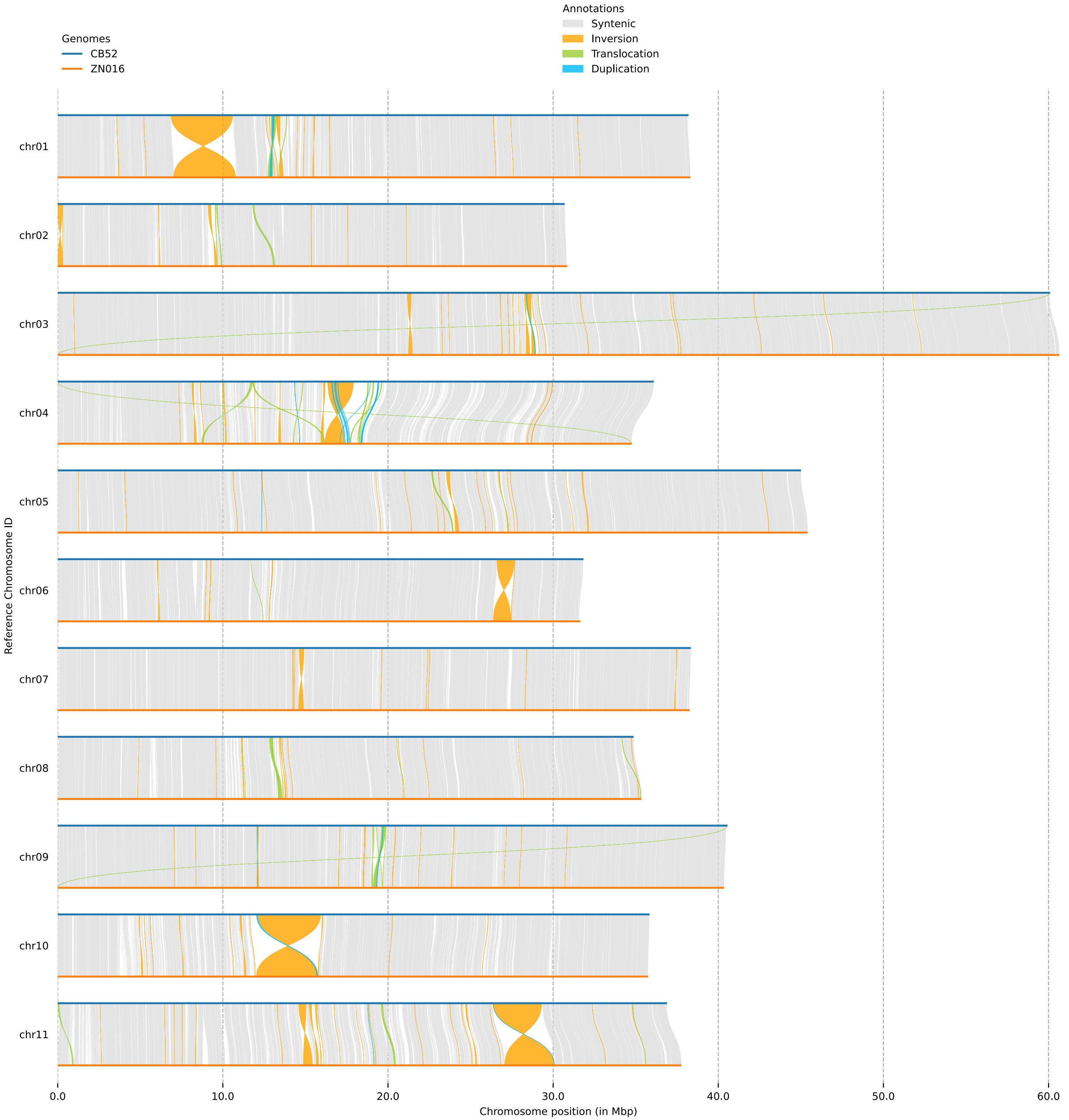


**Supplemental Figure S4E.** Structural variants (of any size) detected by SyRI between the genomes of CB5-2 and ZN016.

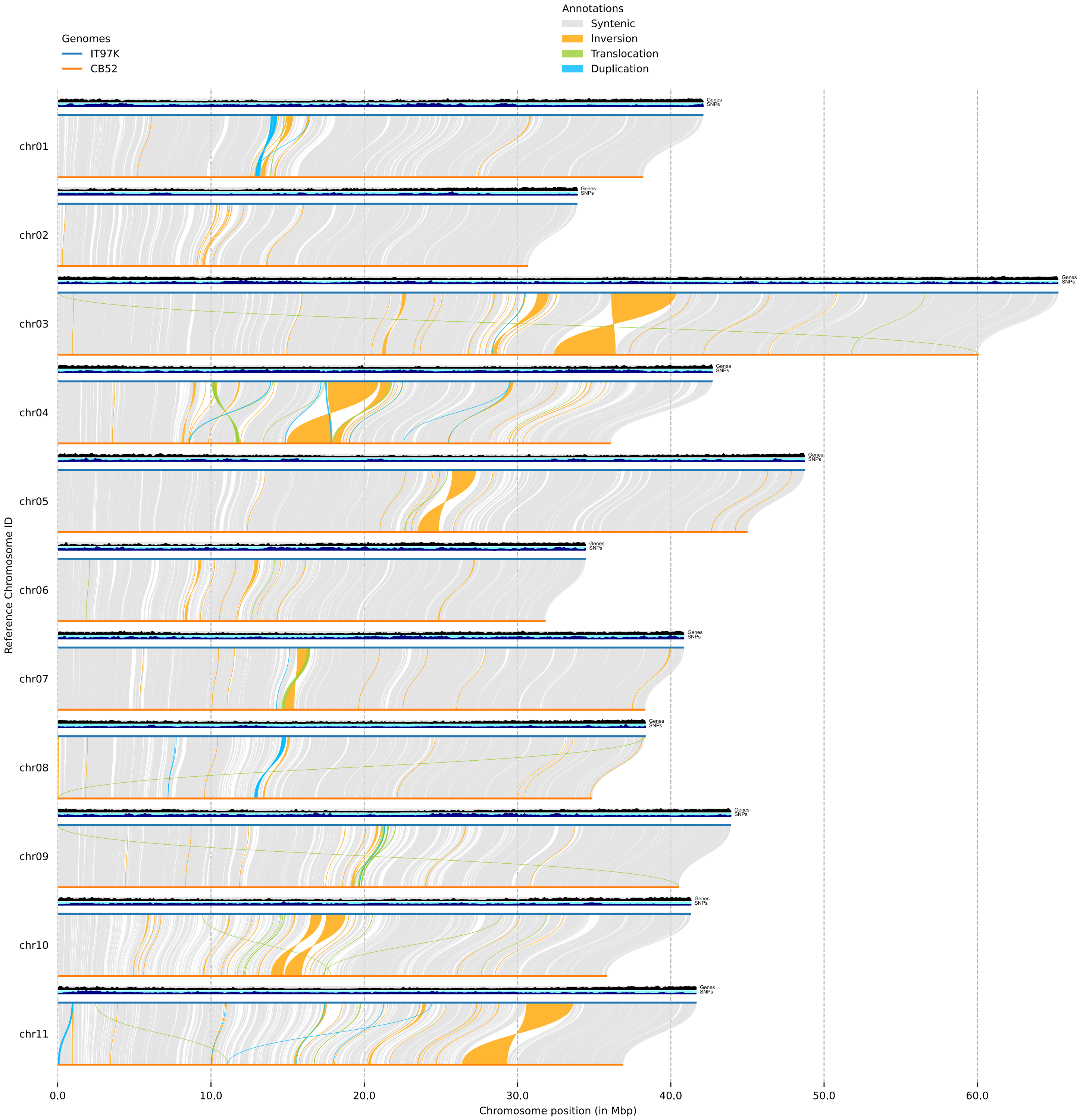


**Supplemental Figure S4F.** Structural variants (of any size) detected by SyRI between the genomes of IT97K and CB5-2.

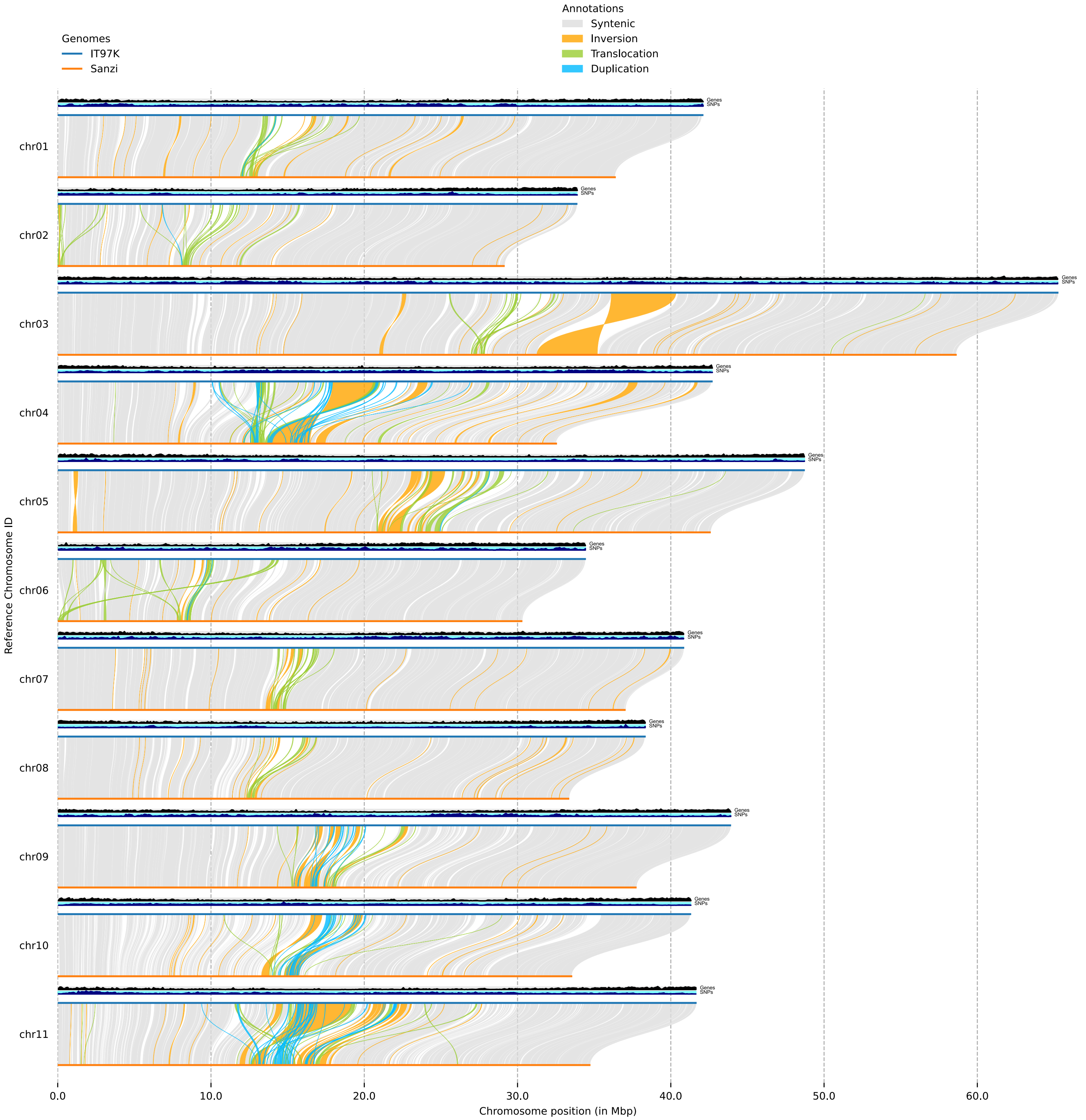


**Supplemental Figure S4G.** Structural variants (of any size) detected by SyRI between the genomes of IT97K and Sanzi.

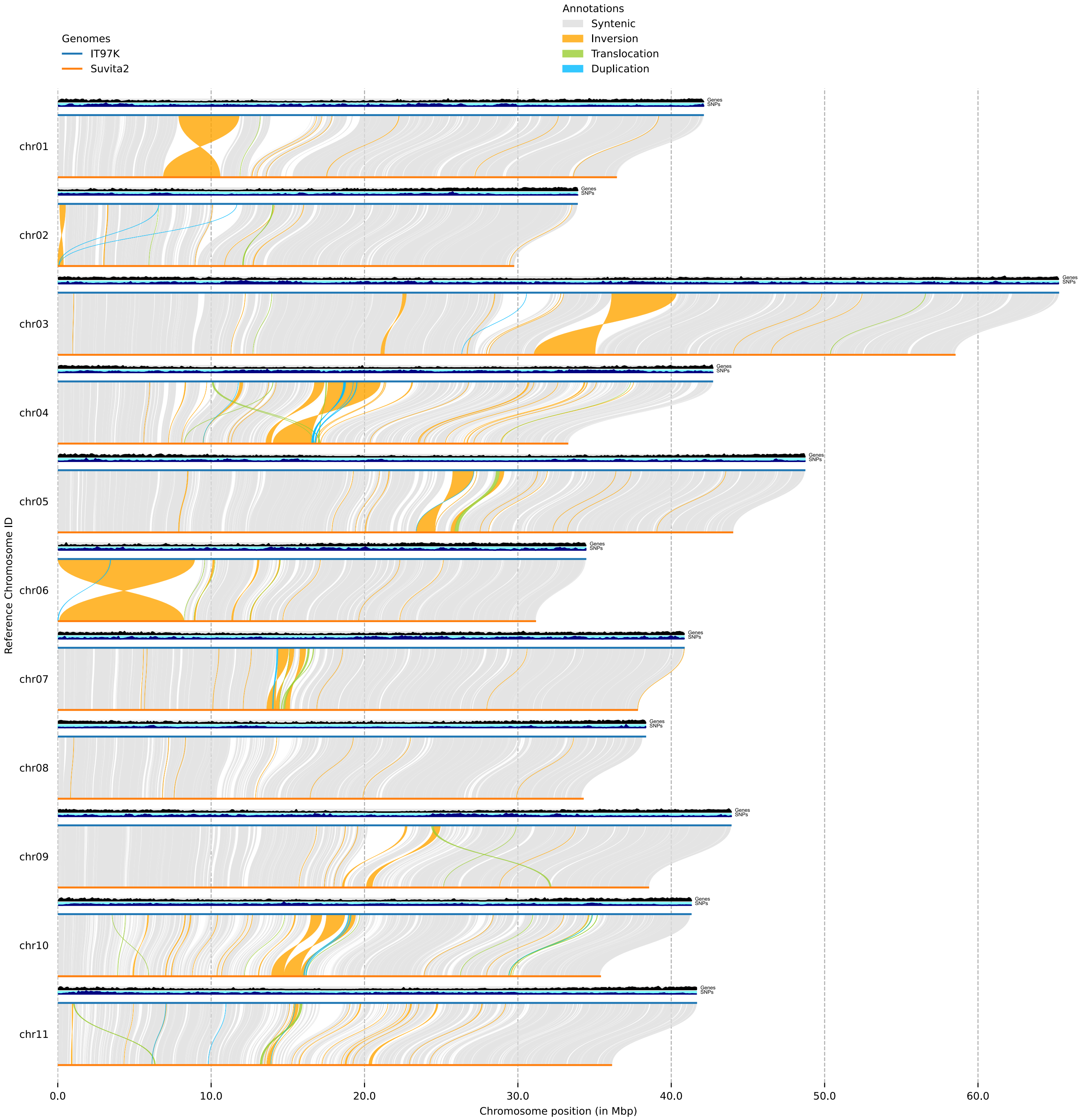


**Supplemental Figure S4H.** Structural variants (of any size) detected by SyRI between the genomes of IT97K and Suvita-2.

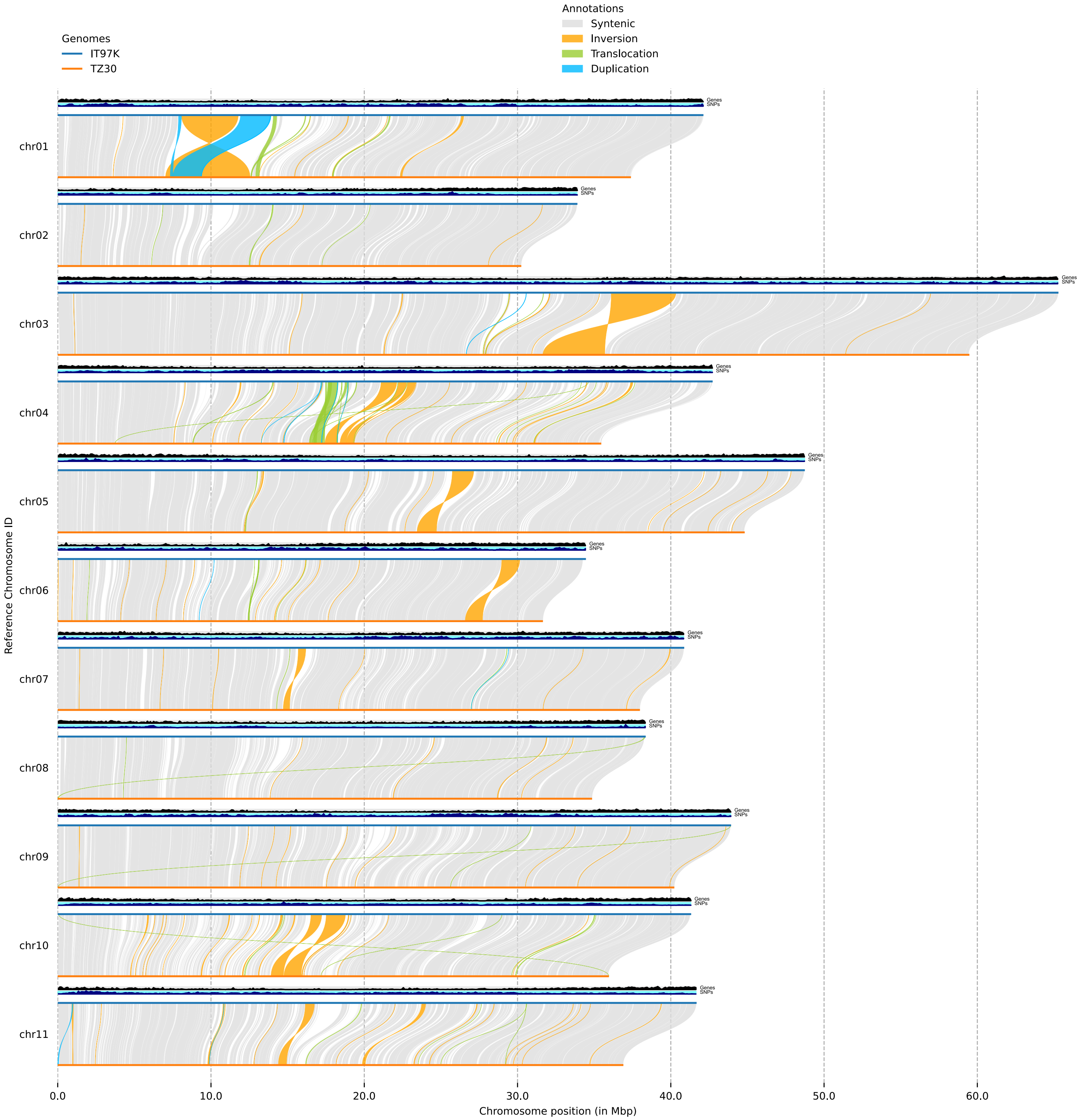


**Supplemental Figure S4I.** Structural variants (of any size) detected by SyRI between the genomes of IT97K and TZ30.

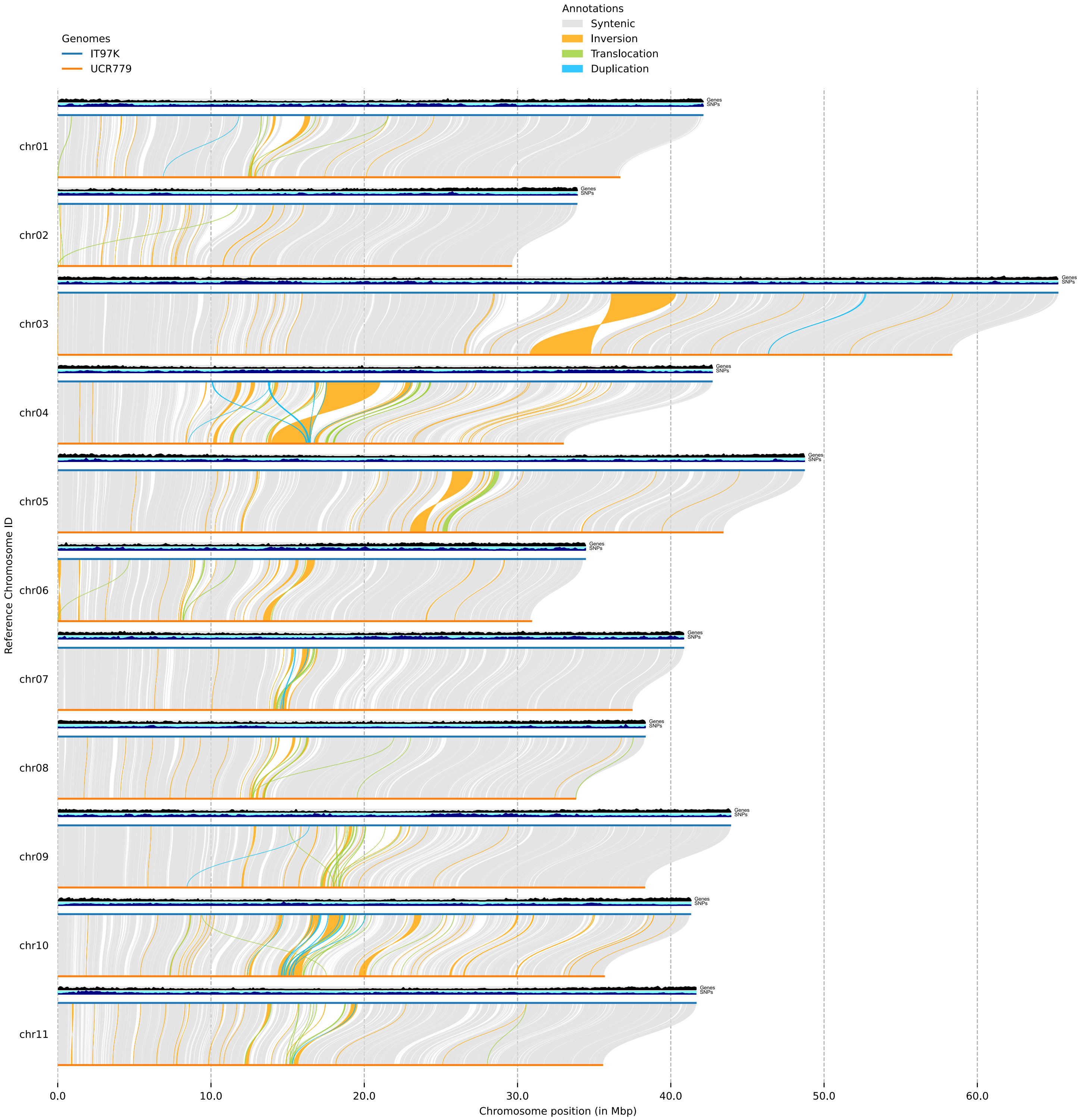


**Supplemental Figure S4J.** Structural variants (of any size) detected by SyRI between the genomes of IT97K and UCR779.

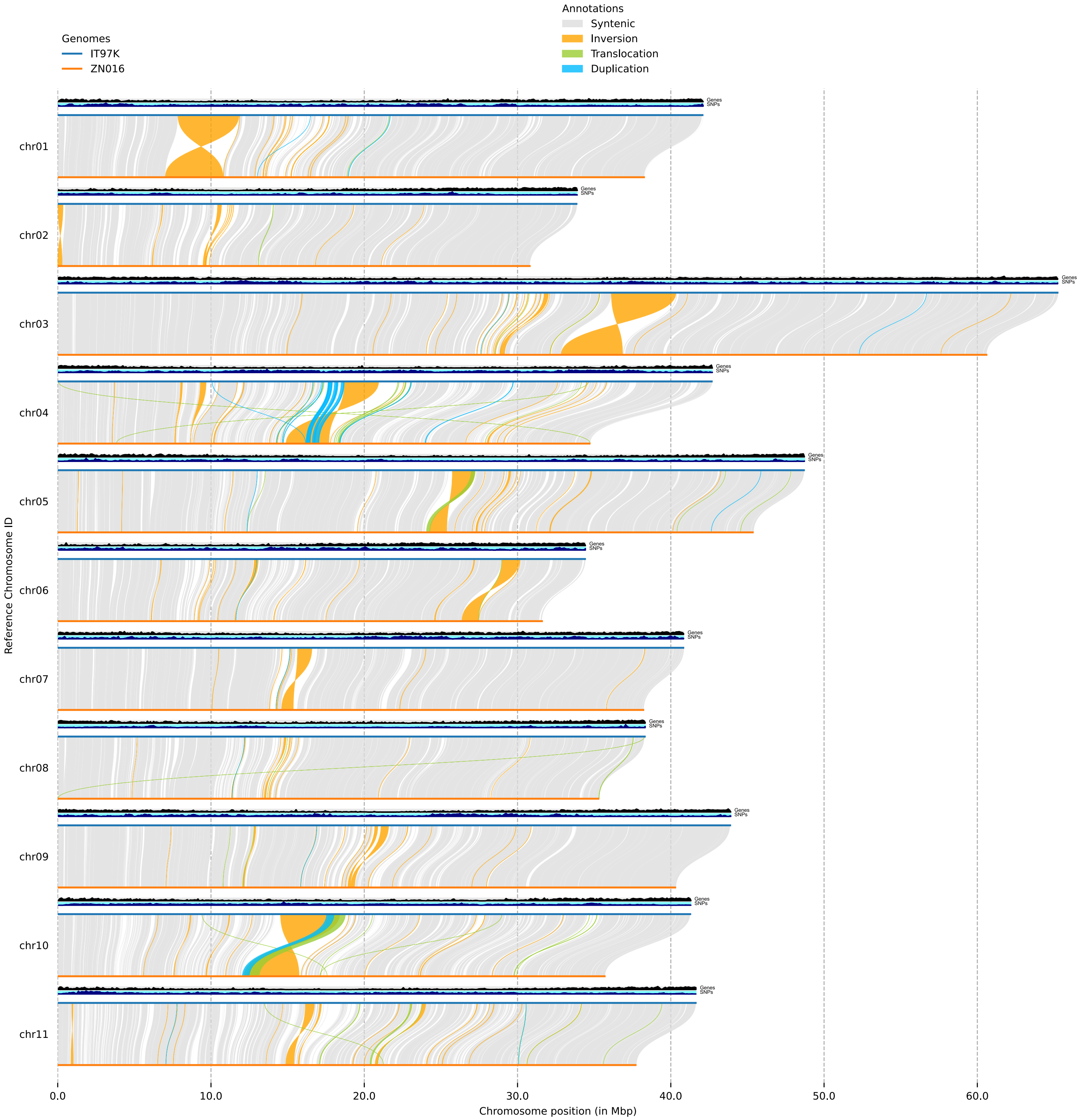


**Supplemental Figure S4K.** Structural variants (of any size) detected by SyRI between the genomes of IT97K and ZN016.

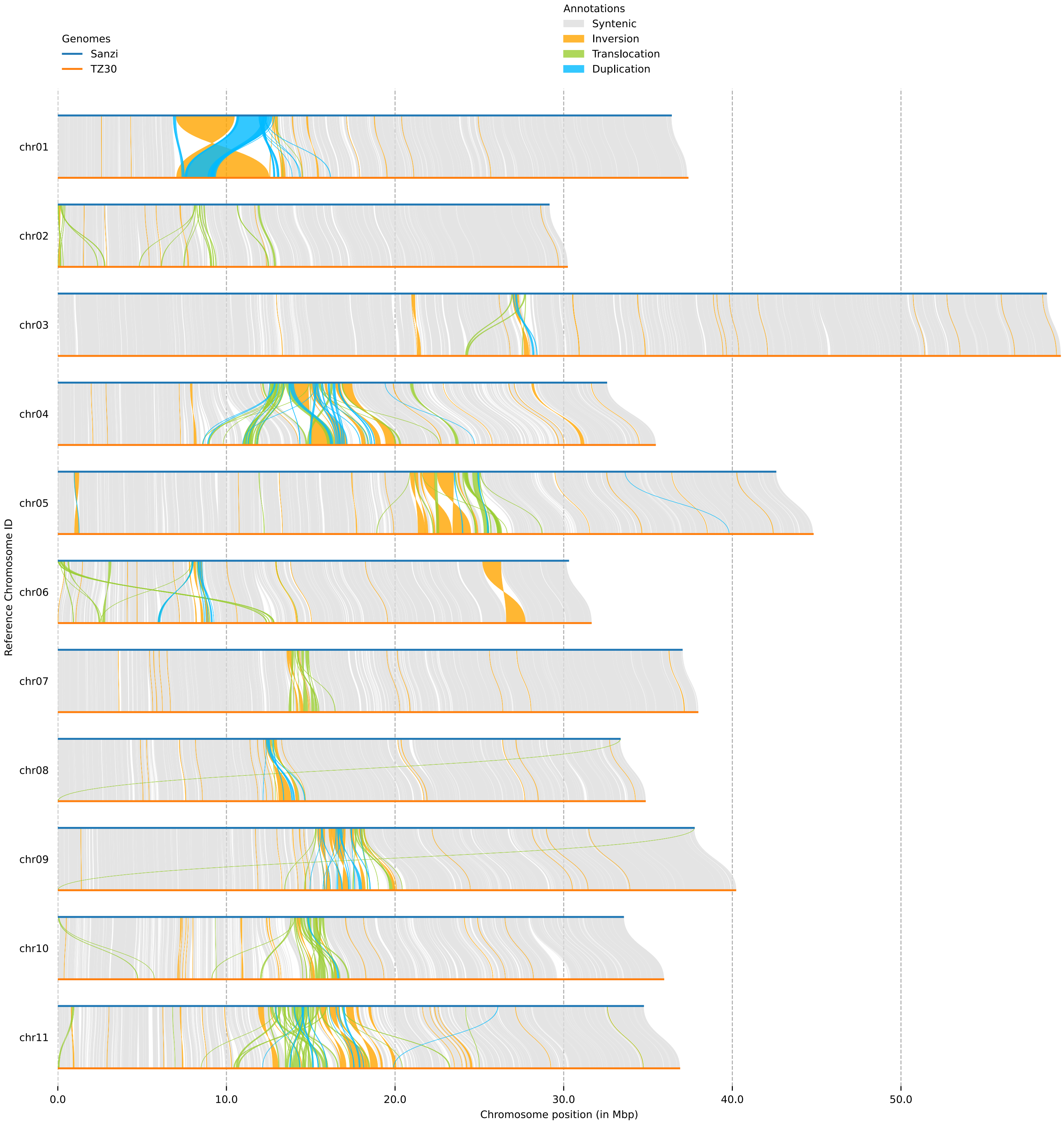


**Supplemental Figure S4L.** Structural variants (of any size) detected by SyRI between the genomes of Sanzi and TZ30.

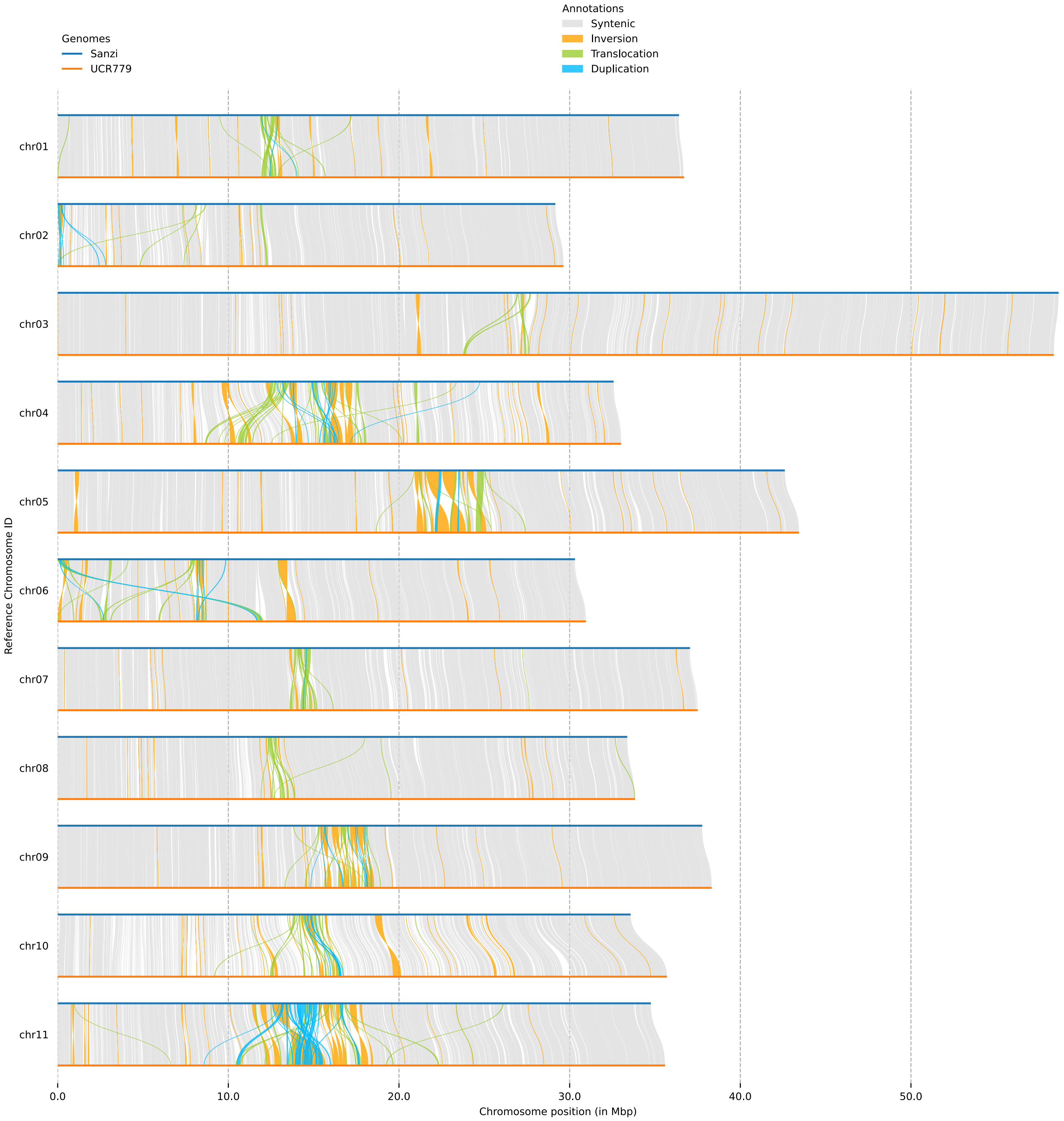


**Supplemental Figure S4M.** Structural variants (of any size) detected by SyRI between the genomes of Sanzi and UCR779.

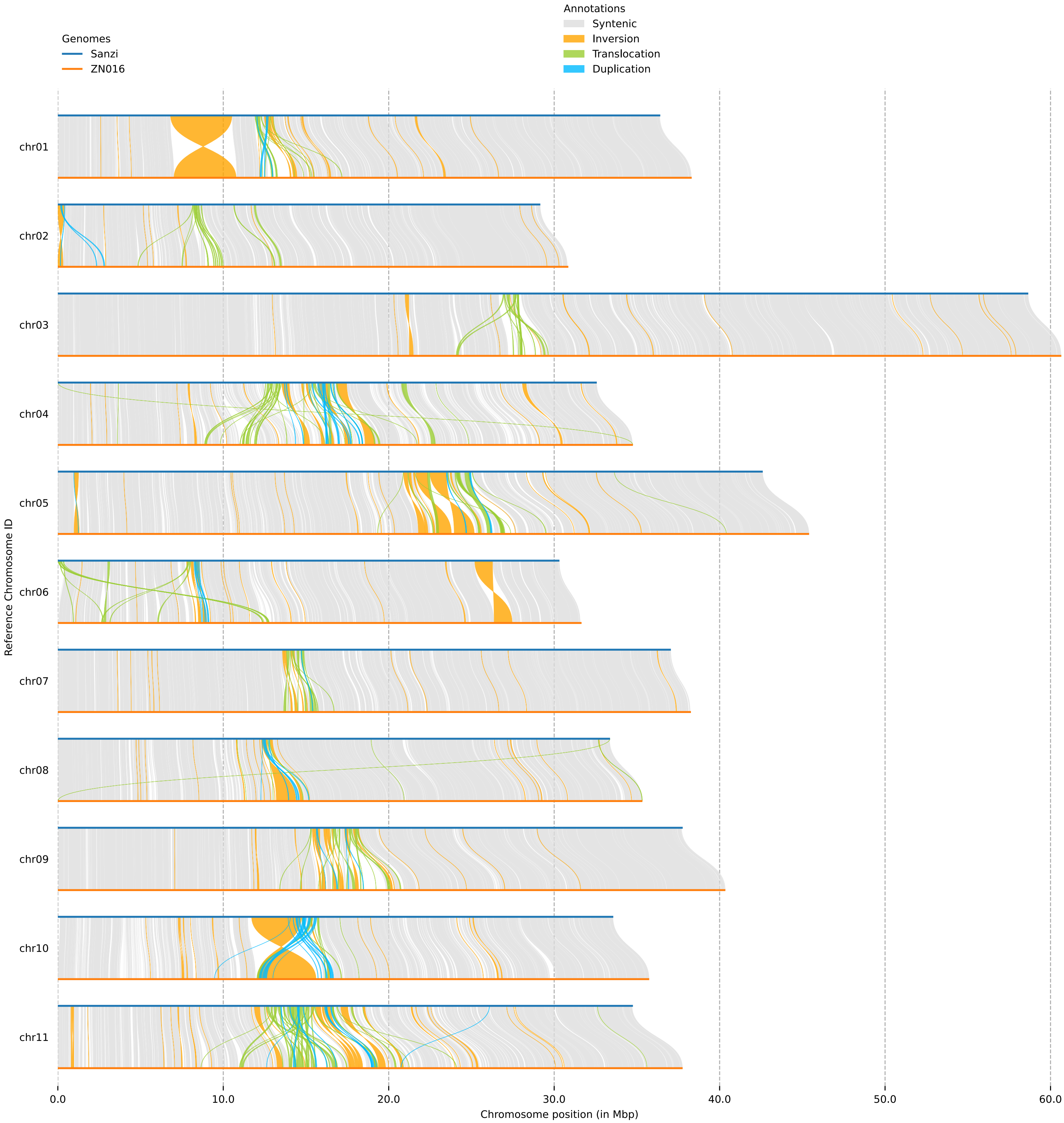


**Supplemental Figure S4N.** Structural variants (of any size) detected by SyRI between the genomes of Sanzi and ZN016.

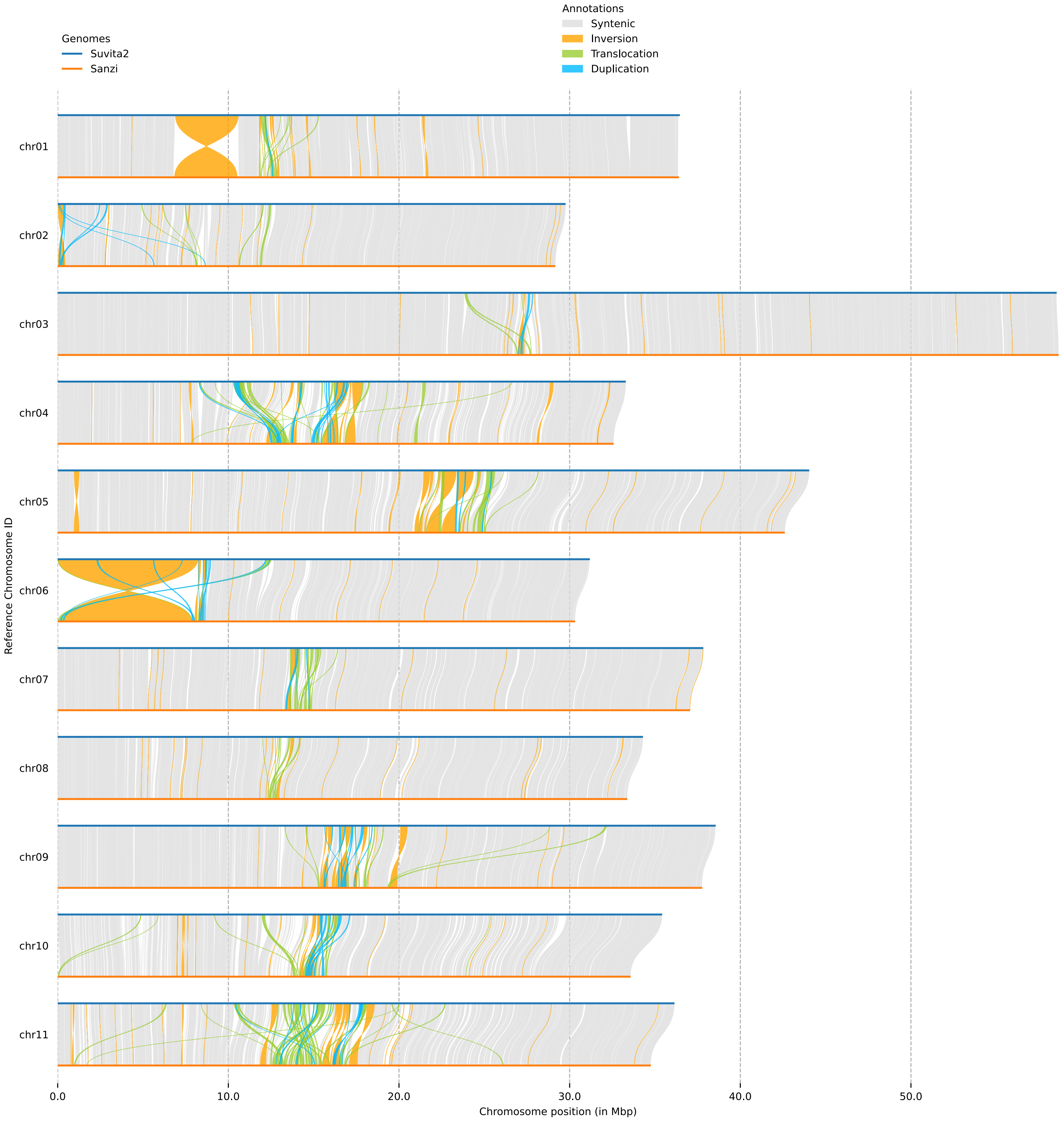


**Supplemental Figure S4O.** Structural variants (of any size) detected by SyRI between the genomes of Suvita-2 and Sanzi.

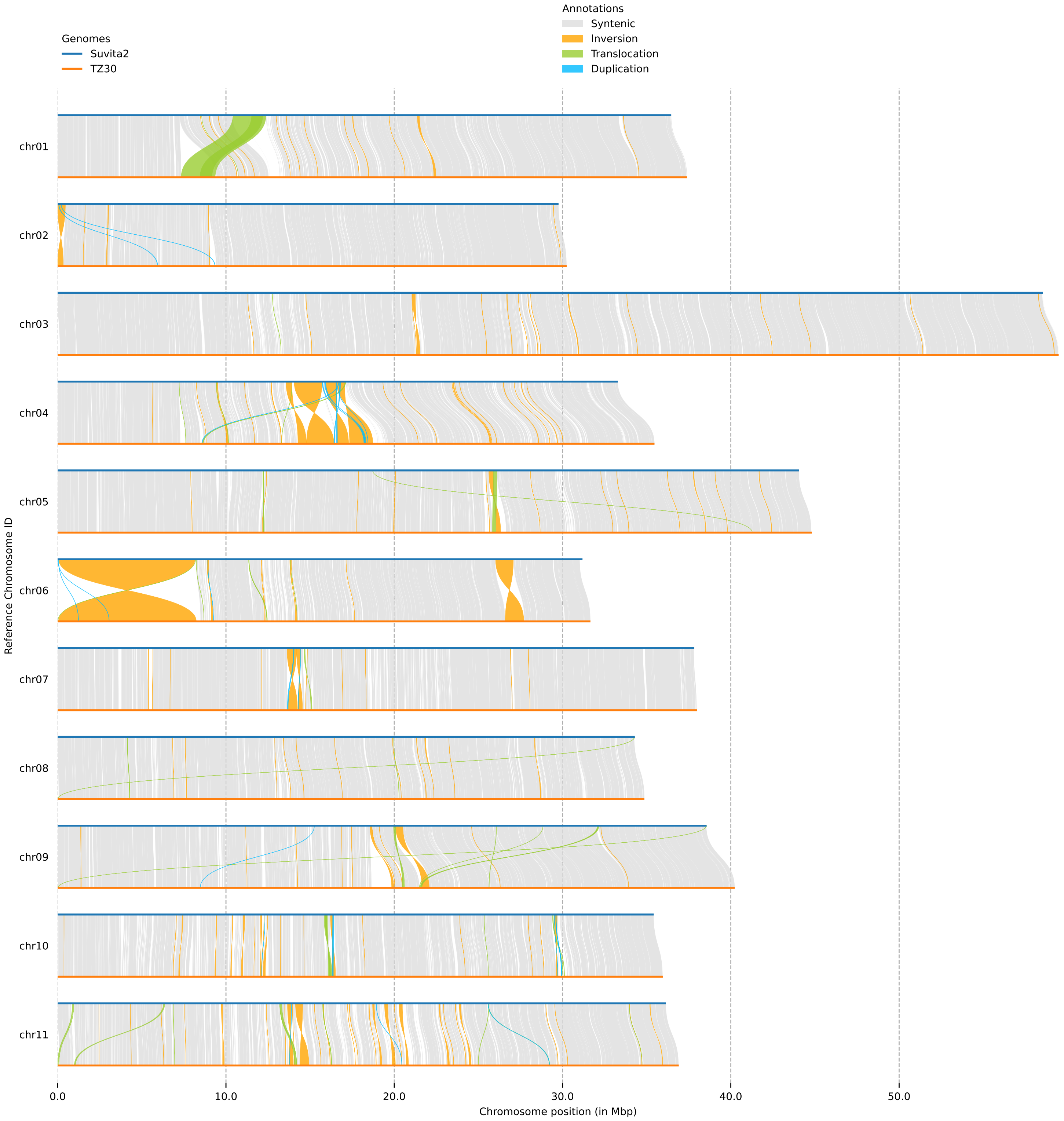


**Supplemental Figure S4P.** Structural variants (of any size) detected by SyRI between the genomes of Suvita-2 and TZ30.

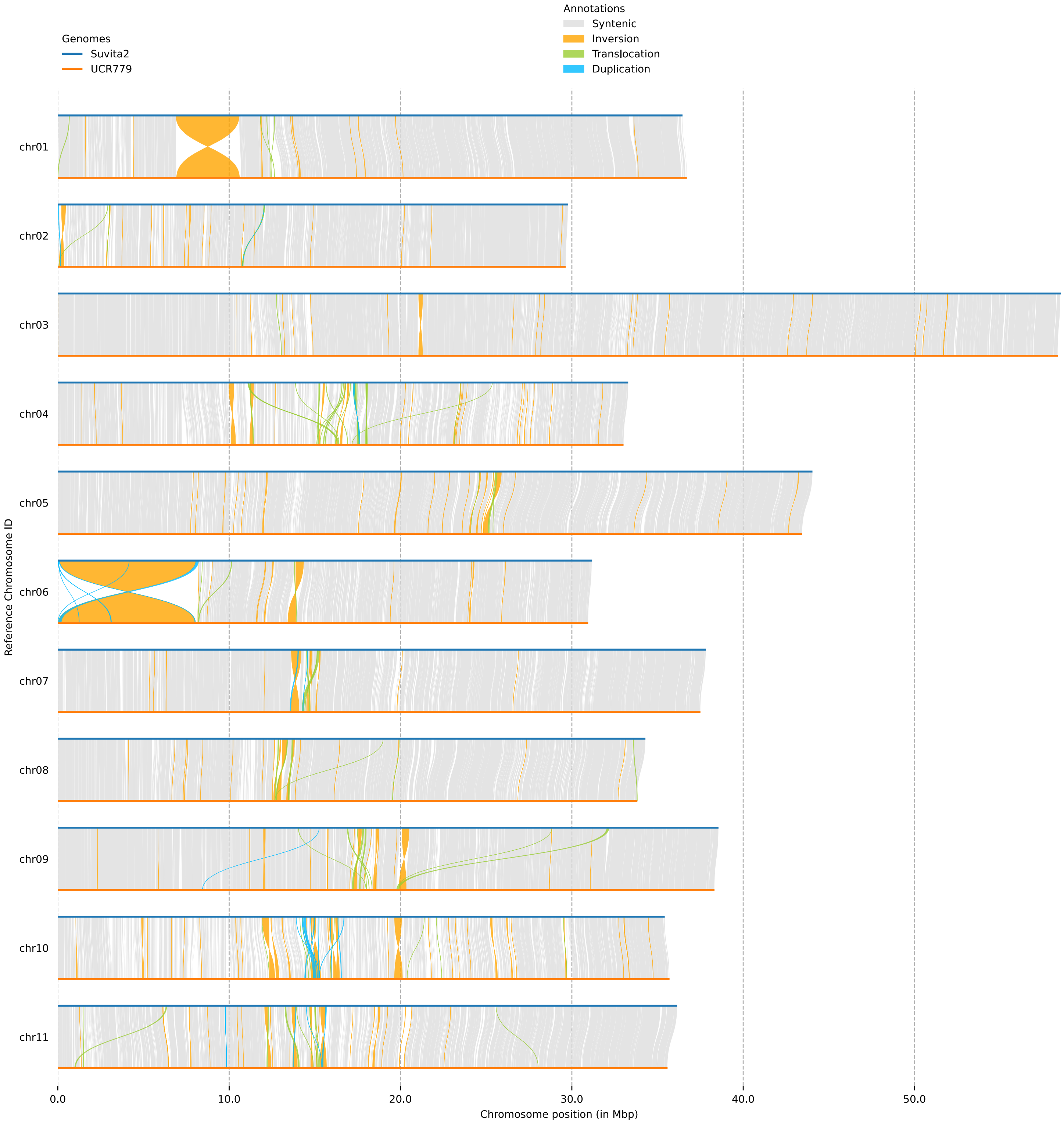


**Supplemental Figure S4Q.** Structural variants (of any size) detected by SyRI between the genomes of Suvita-2 and UCR779.

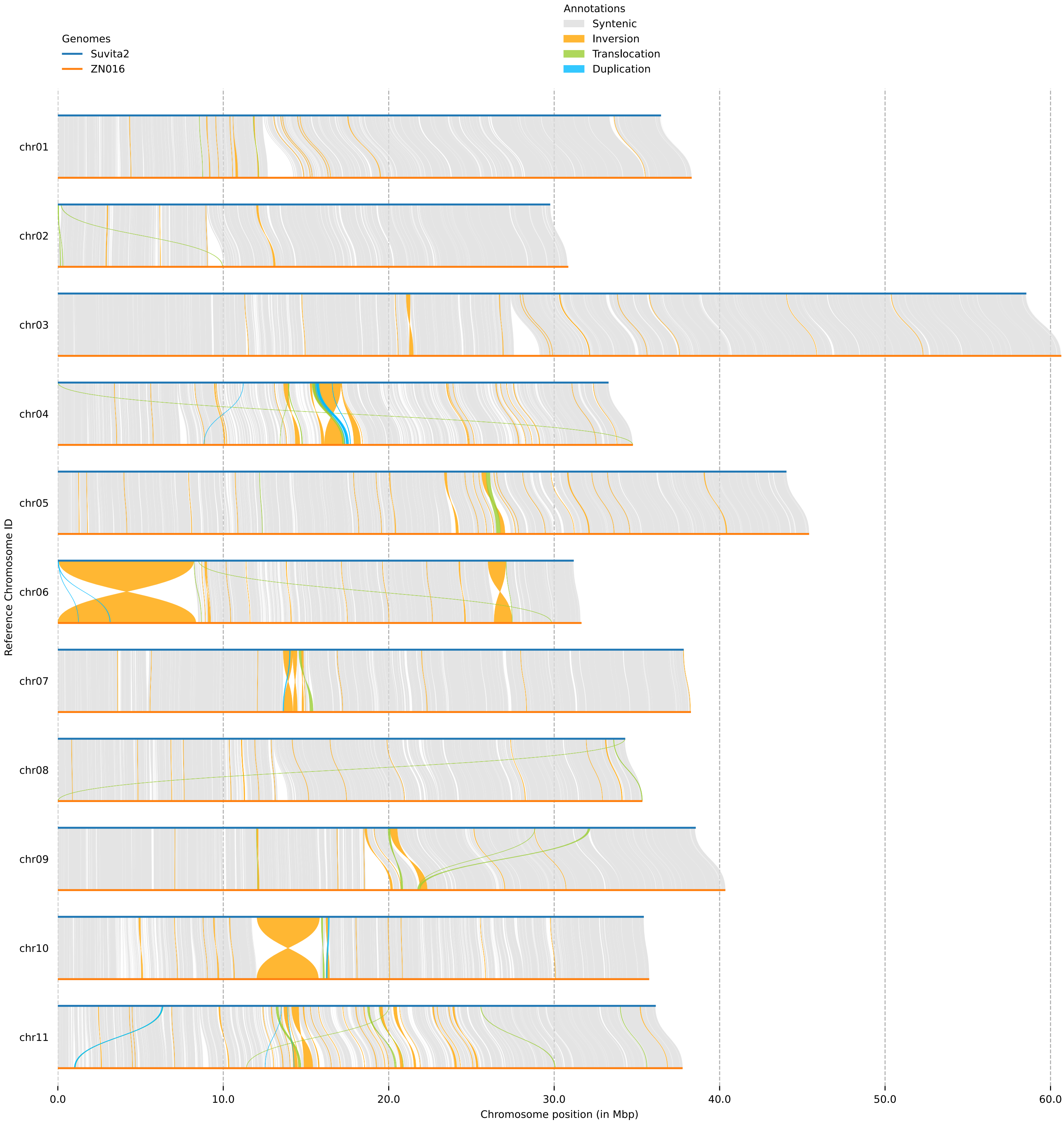


**Supplemental Figure S4R.** Structural variants (of any size) detected by SyRI between the genomes of Suvita-2 and ZN016.

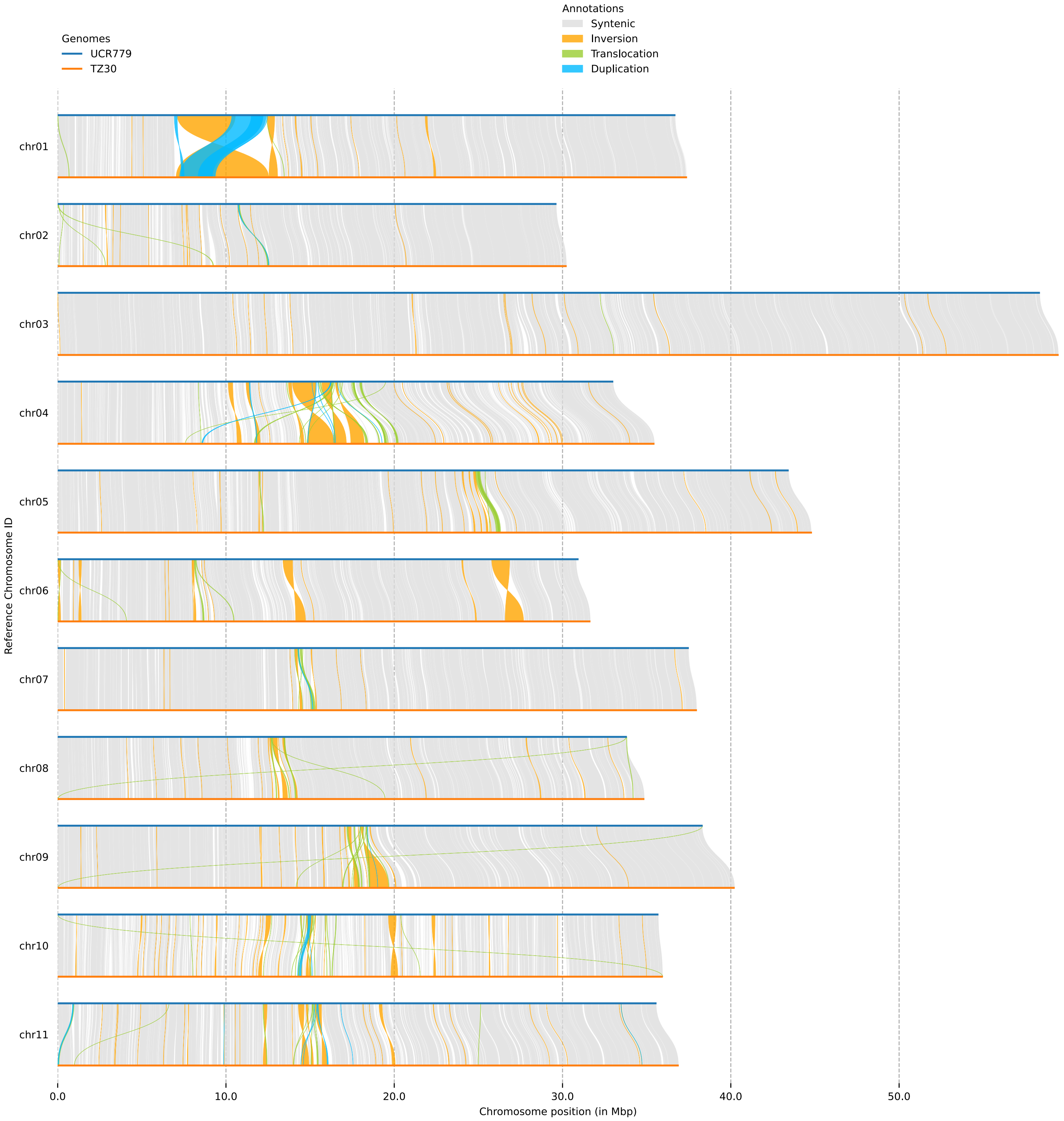


**Supplemental Figure S4S.** Structural variants (of any size) detected by SyRI between the genomes of UCR779 and TZ30.

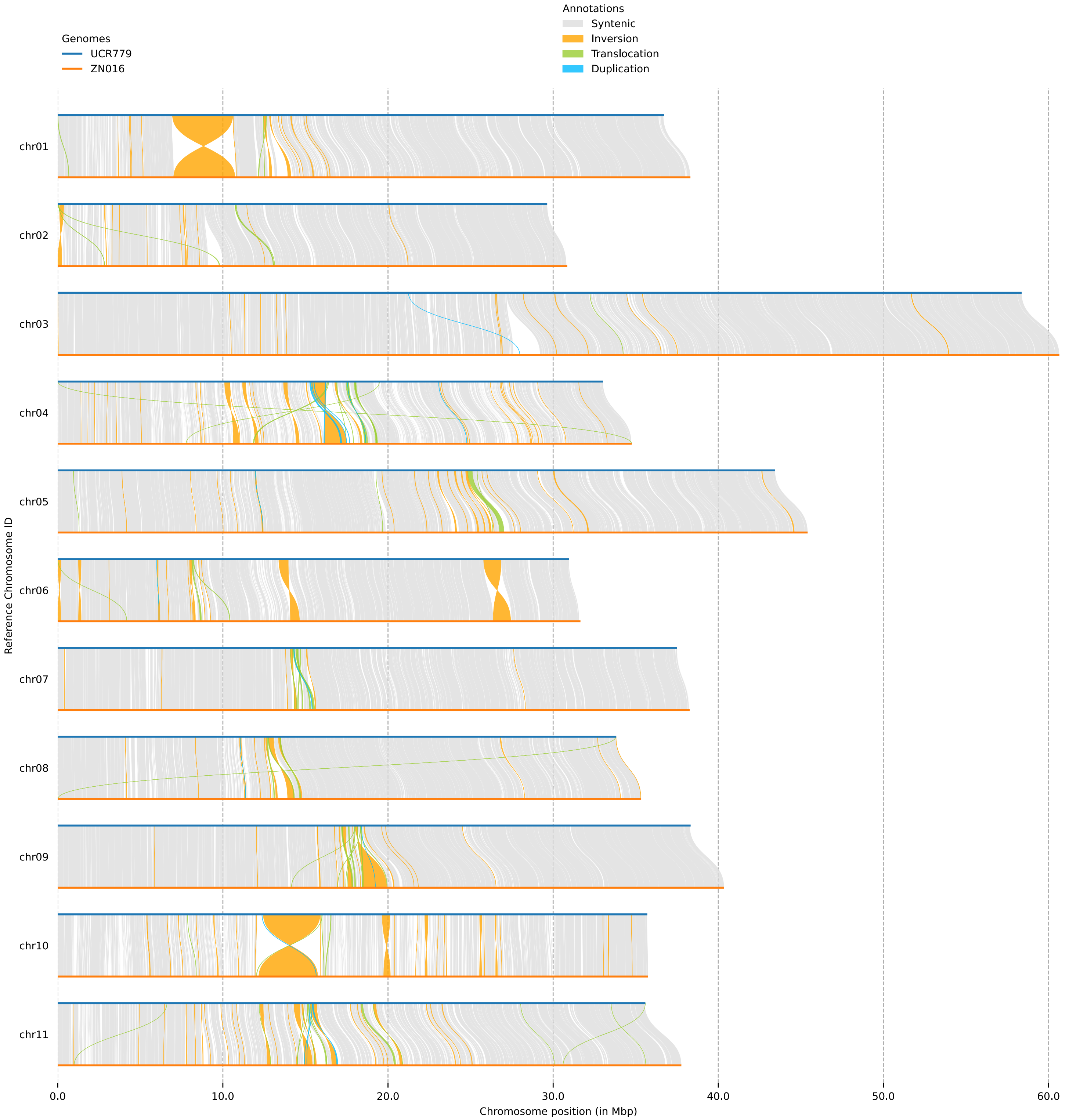


**Supplemental Figure S4T.** Structural variants (of any size) detected by SyRI between the genomes of UCR779 and ZN016.

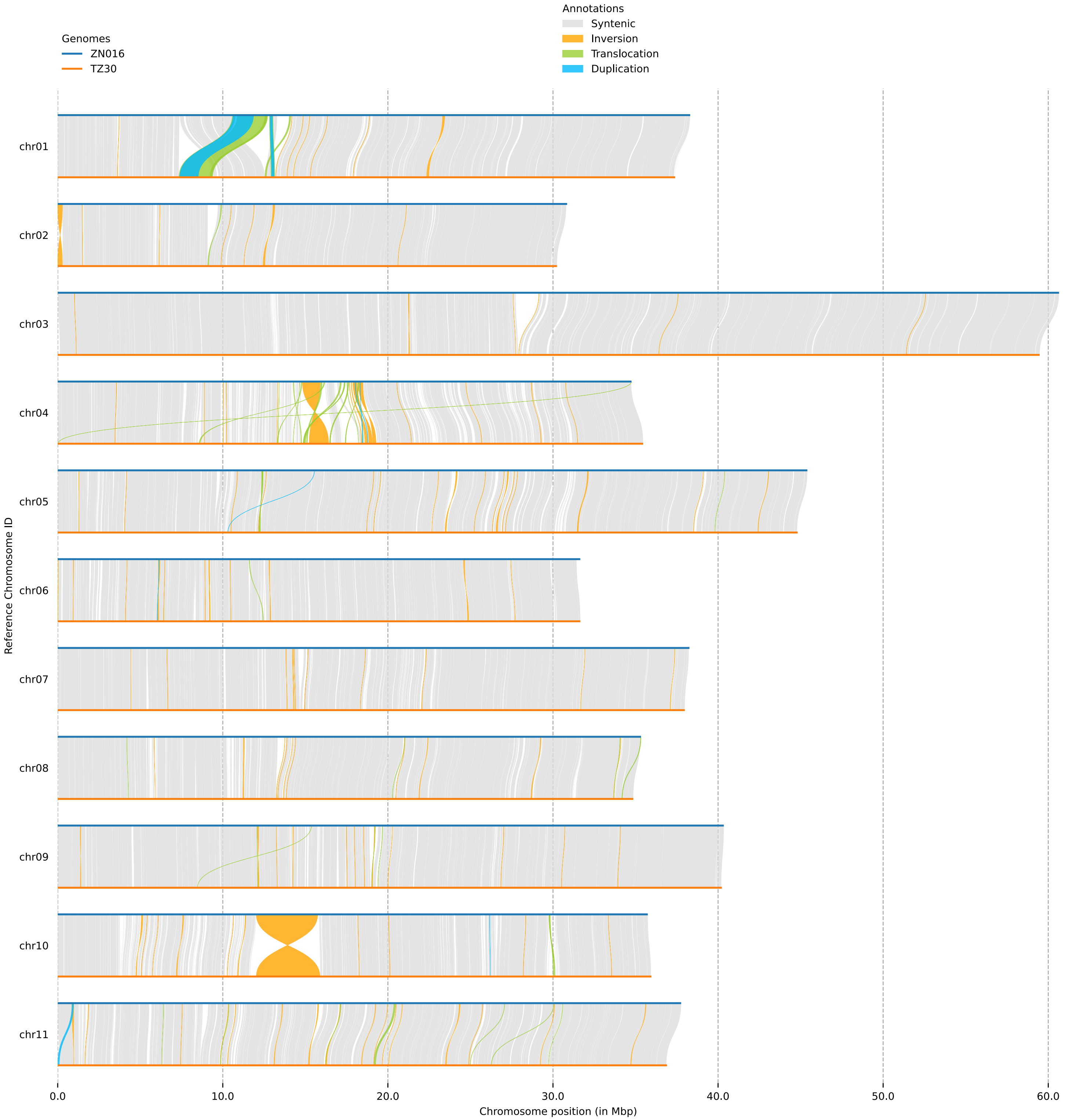


**Supplemental Figure S4U.** Structural variants (of any size) detected by SyRI between the genomes of ZN016 and TZ30.
