## Supplemental Figure S5 for "A view of the pan-genome of domesticated cowpea (*Vigna unguiculata* [L.] Walp.)"

**Supplemental Figure S5A**. **Macrosynteny view with blocks representing regions in the IT97K-499-35 reference cowpea genome with conserved gene order relative to each of the genomes shown as tracks below.** The region from the microsynteny view of Figure 6A is show with a vertical gray bar, and the set of chromosomes displayed is restricted to those showing synteny in that region (i.e., the non-cowpea chromosomes show an apparent lack of synteny on the downstream arm because of genomic rearrangements that have moved corresponding content to other chromosomes than those shown). Various inversions are seen as blocks with orientations opposing those of their neighboring blocks. Gaps in otherwise syntenic regions indicate regions in which gene content diversity outweighs conserved content through presence-absence and copy-number variation.

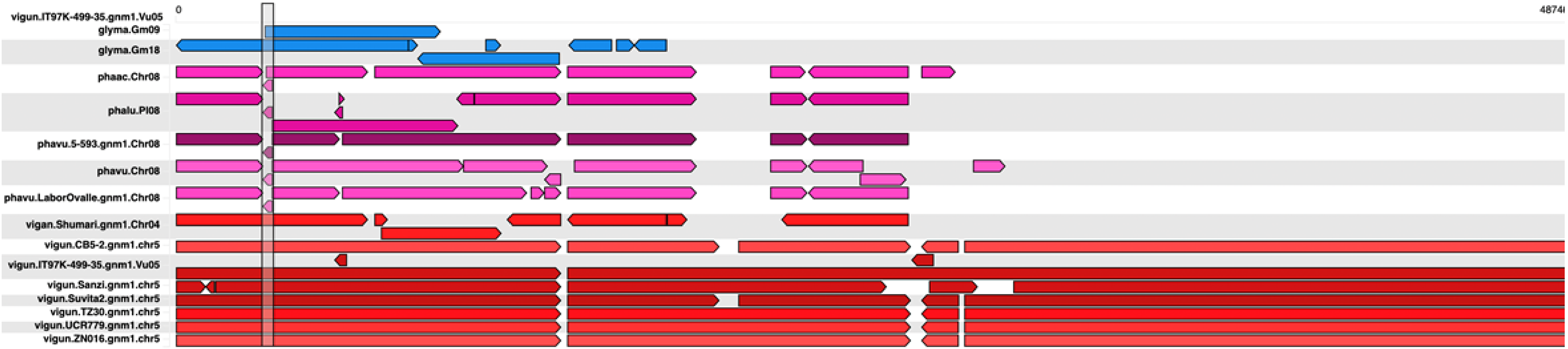

**Supplemental Figure S5B. Counts of genes participating in conserved collinear blocks for all pairwise genome comparisons among the cowpea pangenome members and across representative genomes from several genera in the Phaseoleae tribe.** Self-comparisons are included to illustrate within-species conservation of duplicated content from ancient whole genome duplication (WGD) event shared by subfamily Faboideae species, as well as the more recent WGD in the *Glycine max* genome.

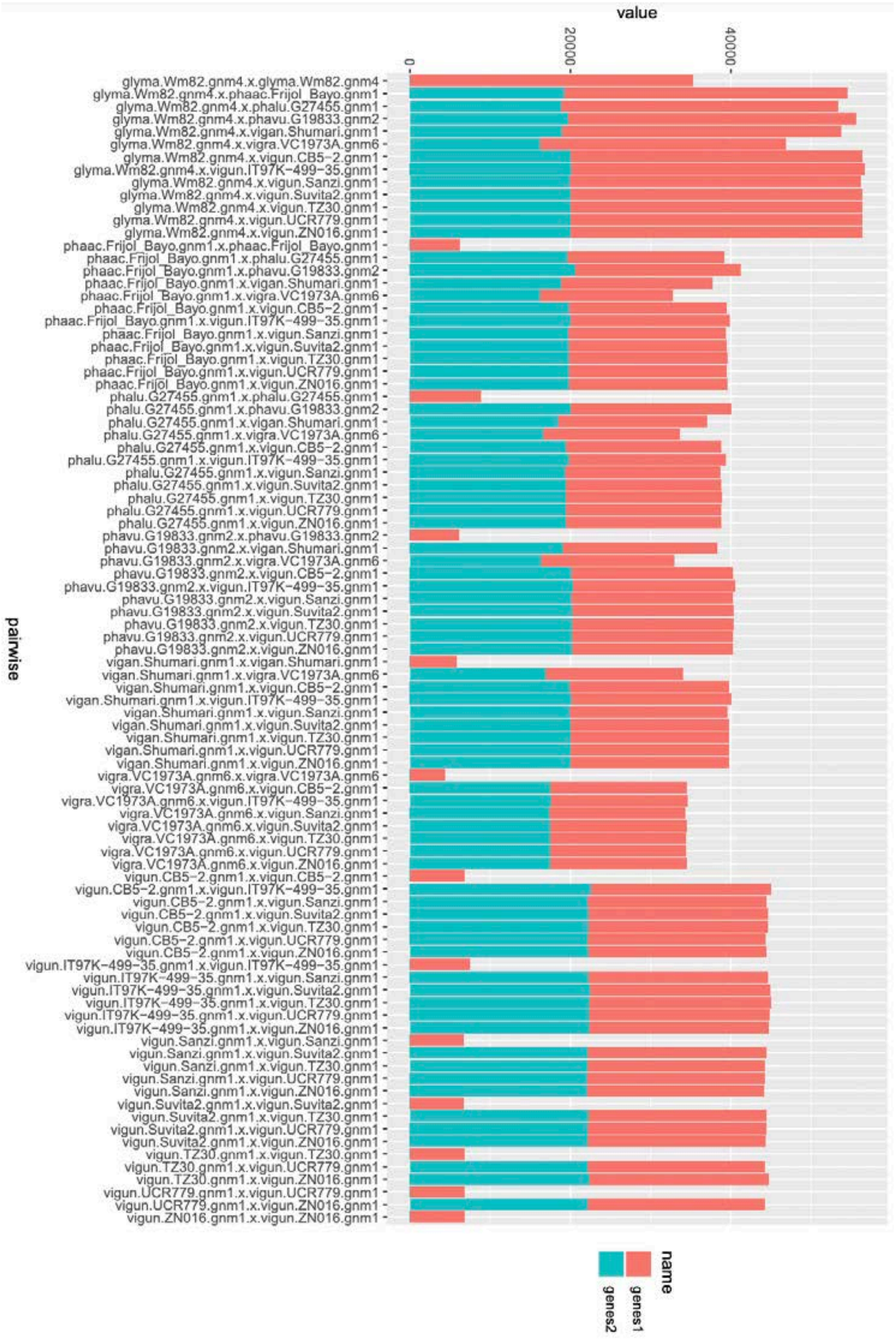
