## Supplemental Table S04 for "A view of the pan-genome of domesticated cowpea (*Vigna unguiculata* [L.] Walp.)"

**Table S04.** **Statistics of the six new assemblies at each step of the Dovetail assembly pipeline.**

|  |  | **CB5-2** | **Suvita-2** | **Sanzi** | **UCR779** | **ZN016** | **TZ30** |
| --- | --- | --- | --- | --- | --- | --- | --- |
| **Meraculous** | assembly (bp) | 445,741,051 | 445,089,603 | 445,080,263 | 451,842,771 | 448,958,014 | 449,031,930 |
|  | # contigs | 29,534 | 34,302 | 33,342 | 34,403 | 29,069 | 31,432 |
|  | N50 (bp) | 39,152 | 30,248 | 31,006 | 33,550 | 42,544 | 34,775 |
|  | longest contig (bp) | 400,619 | 303,123 | 254,506 | 323,431 | 430,667 | 266,922 |
| **Chicago** | assembly (bp) | 447,987,355 | 447,527,988 | 447,138,401 | 453,839,006 | 451,080,434 | 451,434,994 |
|  | # contigs/scaffolds | 6,744 | 9,696 | 12,668 | 14,264 | 7,519 | 7,071 |
|  | N50 (bp) | 5,801,957 | 1,887,887 | 609,507 | 613,651 | 2,390,195 | 3,108,375 |
|  | longest contig (bp) | 15,476,412 | 8,143,501 | 6,203,760 | 8,226,530 | 11,401,244 | 11,167,207 |
| **Hi-C** | assembly (bp) | 448,041,251 | 447581292 | 447,237,961 | 453,955,886 | 451,129,907 | 451,466,780 |
|  | # contigs/scaffolds | 6,559 | 9,162 | 11,661 | 13,085 | 7,041 | 6,790 |
|  | N50 (bp) | 21,318,834 | 34,289,026 | 17,830,733 | 19,877,087 | 35,743,455 | 23,295,971 |
|  | longest contig (bp) | 31,414,800 | 58,539,223 | 30,787,568 | 58,369,212 | 60,653,587 | 59,481,915 |
| **AllMaps** | assembly (bp) | 448,043,751 | 447,585,192 | 447,277,261 | 453,970,486 | 451,130,807 | 451,468,680 |
|  | # contigs/scaffolds | 6,534 | 9,123 | 11,268 | 12,939 | 7,032 | 6,771 |
|  | N50 (bp) | 36,897,245 | 36,142,647 | 34,759,918 | 35,700,653 | 37,764,243 | 36,906,789 |
|  | longest contig (bp) | 60,086,998 | 58,539,223 | 58,655,738 | 58,369,212 | 60,653,587 | 59,481,915 |
