## Supplemental Table S05 for "A view of the pan-genome of domesticated cowpea (*Vigna unguiculata* [L.] Walp.)"

|  | **IT97K-499-35** | | | **CB5-2** | | | **Sanzi** | | | **ZN016** | | | **TZ30** | | |
| --- | --- | --- | --- | --- | --- | --- | --- | --- | --- | --- | --- | --- | --- | --- | --- |
| **Chr.** | **start** | **end** | **size** | **start** | **end** | **size** | **start** | **end** | **size** | **start** | **end** | **size** | **start** | **end** | **size** |
| **1** | 14,698,036 | 16,525,496 | 1,827,460 | 12,796,513 | 14,236,341 | 1,439,828 |  |  | - | 13,289,165 | 17,690,285 | 4,401,120 |  |  | - |
| **2** | 10,238,236 | 14,020,258 | 3,782,022 |  |  | - |  |  | - |  |  | - |  |  | - |
| **3** | 30,476,981 | 31,470,261 | 993,280 |  |  | - |  |  | - |  |  | - |  |  | - |
| **4** | 19,069,641 | 21,130,843 | 2,061,202 | 16,669,712 | 18,502,124 | 1,832,412 | 15,648,675 | 16,247,976 | 599,301 |  |  | - |  |  | - |
| **5** | 25,704,431 | 33,885,354 | 8,180,923 | 26,553,015 | 26,811,298 | 258,283 |  |  | - | 25,404,785 | 27,305,040 | 1,900,255 |  |  | - |
| **6** | 9,156,830 | 9,235,637 | 78,807 |  |  | - |  |  | - |  |  | - |  |  | - |
| **7** | 16,587,031 | 16,604,960 | 17,929 | 14,903,264 | 14,933,285 | 30,021 |  |  | - | 14,573,279 | 14,601,618 | 28,339 | 13,530,446 | 14,858,230 | 1,327,784 |
| **8** | 14,914,119 | 15,164,402 | 250,283 | 12,820,933 | 14,867,106 | 2,046,173 |  |  | - | 13,476,113 | 14,282,079 | 805,966 |  |  | - |
| **9** | 20,802,610 | 22,685,597 | 1,882,987 |  |  | - |  |  | - |  |  | - |  |  | - |
| **10** | 18,917,563 | 19,028,450 | 110,887 |  |  | - |  |  | - |  |  | - |  |  | - |
| **11** | 17,283,961 | 18,283,861 | 999,900 |  |  | - |  |  | - |  |  | - |  |  | - |
| **Total** |  |  | 20,185,680 |  |  | 5,606,717 |  |  | 599,301 |  |  | 7,135,680 |  |  | 1,327,784 |

**Table S05. Putative centromeric region coordinates (all numbers are bp).**
