## Supplemental Table S06 for "A view of the pan-genome of domesticated cowpea (*Vigna unguiculata* [L.] Walp.)"

**Table S06.** **Gene annotation statistics.**

|  | **IT97K-499-35** | **CB5-2** | **Suvita-2** | **Sanzi** | **UCR779** | **ZN016** | **TZ30** |
| --- | --- | --- | --- | --- | --- | --- | --- |
| **Genome size (bp)** | 519,435,864 | 448,043,751 | 447,585,192 | 447,277,261 | 453,970,486 | 451,130,807 | 451,468,680 |
| Primary transcripts (loci) | 31,948 | 28,297 | 28,545 | 28,461 | 28,562 | 27,723 | 27,742 |
| Alternative transcripts | 22,536 | 15,852 | 17,115 | 16,754 | 15,704 | 15,420 | 15,088 |
| Total transcripts | 54,484 | 44,149 | 45,660 | 45,215 | 44,266 | 43,143 | 42,830 |
| **Primary transcripts** |  |  |  |  |  |  |  |
| Average number of exons | 5.2 | 5.4 | 5.4 | 5.4 | 5.4 | 5.4 | 5.4 |
| Median exon length (bp) | 169 | 163 | 164 | 163 | 163 | 162 | 162 |
| Median intron length (bp) | 174 | 178 | 178 | 178 | 178 | 178 | 177 |
| **Gene model support** |  |  |  |  |  |  |  |
| Any EST support (# of gene models) | 28,060 | 23,942 | 24,768 | 24,428 | 24,191 | 21,506 | 21,398 |
| EST support 100% of length | 26,764 | 21,741 | 22,719 | 22,053 | 21,995 | 21,812 | 21,715 |
| EST support >95% of length | 27,109 | 22,378 | 23,339 | 22,754 | 22,630 | 21,923 | 21,823 |
| EST support >90% of length | 27,251 | 22,602 | 23,532 | 23,021 | 22,838 | 22,122 | 22,023 |
| EST support >75% of length | 27,467 | 22,942 | 23,881 | 23,392 | 23,170 | 22,362 | 22,289 |
| EST support >50% of length | 27,654 | 23,325 | 24,236 | 23,759 | 23,550 | 21,506 | 11,763 |
| **Peptide homology coverage of 100%** | 23,431 | 23,264 | 23,466 | 23,236 | 22,889 | 22,515 | 22,532 |
| **Peptide homology coverage >95%** | 27,372 | 25,697 | 25,950 | 25,830 | 25,796 | 23,953 | 23,957 |
| **Peptide homology coverage >90%** | 28,421 | 26,281 | 26,543 | 26,431 | 26,430 | 25,375 | 25,380 |
| **Peptide homology coverage of >75%** | 29,913 | 27,333 | 27,483 | 27,426 | 27,406 | 26,272 | 26,285 |
| **Peptide homology coverage >50%** | 30,839 | 27,930 | 28,071 | 28,034 | 28,073 | 22,515 | 22,532 |
| **Pfam annotation** | 23,798 | 21,759 | 21,845 | 21,827 | 21,888 | 21,665 | 21,701 |
| **Panther annotation** | 28,046 | 25,666 | 25,769 | 25,787 | 25,881 | 25,488 | 25,526 |
| **KOG annotation** | 13,270 | 12,307 | 12,389 | 12,335 | 12,331 | 12,313 | 12,314 |
| **KEGG orthology annotation** | 9,138 | 8,535 | 8,564 | 8,572 | 8,517 | 8,574 | 8,576 |
| **E.C. annotation** | 9,032 | 8,283 | 8,355 | 8,327 | 8,357 | 8,175 | 8,186 |
