## Supplemental Table S07 for "A view of the pan-genome of domesticated cowpea (*Vigna unguiculata* [L.] Walp.)"

**Table S07. BUSCO v4 completeness results.**

|  |  | **IT97K-499-35** | | **CB5-2** | | **Suvita-2** | | **Sanzi** | | **UCR779** | | **ZN016** | | **TZ30** | |
| --- | --- | --- | --- | --- | --- | --- | --- | --- | --- | --- | --- | --- | --- | --- | --- |
| **Genome** | **Complete** | 1595 | 98.8% | 1574 | 97.5% | 1580 | 97.9% | 1581 | 98.0% | 1574 | 97.5% | 1589 | 98.5% | 1583 | 98.1% |
|  | **Complete and single-copy** | 1548 | 95.9% | 1538 | 95.3% | 1547 | 95.8% | 1545 | 95.7% | 1539 | 95.4% | 1552 | 96.2% | 1548 | 95.9% |
|  | **Complete and duplicated** | 47 | 2.9% | 36 | 2.2% | 33 | 2.0% | 36 | 2.2% | 35 | 2.2% | 37 | 2.3% | 35 | 2.2% |
|  | **Fragmented** | 8 | 0.5% | 23 | 1.4% | 18 | 1.1% | 15 | 0.9% | 22 | 1.4% | 10 | 0.6% | 10 | 0.6% |
|  | **Missing** | 11 | 0.7% | 17 | 1.1% | 16 | 1.0% | 18 | 1.1% | 18 | 1.1% | 15 | 0.9% | 21 | 1.3% |
| **Transcripts (primary)** | **Complete** | 1594 | 98.8% | 1570 | 97.3% | 1582 | 98.0% | 1585 | 98.2% | 1581 | 98.0% | 1584 | 98.1% | 1580 | 97.9% |
|  | **Complete and single-copy** | 1546 | 95.8% | 1534 | 95.0% | 1547 | 95.8% | 1550 | 96.0% | 1548 | 95.9% | 1547 | 95.8% | 1547 | 95.8% |
|  | **Complete and duplicated** | 58 | 3.0% | 36 | 2.2% | 35 | 2.2% | 35 | 2.2% | 33 | 2.0% | 37 | 2.3% | 33 | 2.0% |
|  | **Fragmented** | 8 | 0.5% | 24 | 1.5% | 19 | 1.2% | 13 | 0.8% | 14 | 0.9% | 12 | 0.7% | 14 | 0.9% |
|  | **Missing** | 12 | 0.7% | 20 | 1.2% | 13 | 0.8% | 16 | 1.0% | 19 | 1.2% | 18 | 1.1% | 20 | 1.2% |
| **Proteins (primary)** | **Complete** | 1595 | 98.8% | 1569 | 97.2% | 1584 | 98.1% | 1587 | 98.3% | 1585 | 98.2% | 1584 | 98.1% | 1582 | 98.0% |
|  | **Complete and single-copy** | 1545 | 95.7% | 1531 | 94.9% | 1549 | 96.0% | 1550 | 96.0% | 1552 | 96.2% | 1550 | 96.0% | 1551 | 96.1% |
|  | **Complete and duplicated** | 50 | 3.1% | 38 | 2.4% | 35 | 2.2% | 37 | 2.3% | 33 | 2.0% | 34 | 2.1% | 31 | 1.9% |
|  | **Fragmented** | 11 | 0.7% | 31 | 1.9% | 21 | 1.3% | 17 | 1.1% | 18 | 1.1% | 18 | 1.1% | 17 | 1.1% |
|  | **Missing** | 8 | 0.5% | 14 | 0.9% | 9 | 0.6% | 10 | 0.6% | 11 | 0.7% | 12 | 0.7% | 15 | 0.9% |
