## Supplemental Table S11 for "A view of the pan-genome of domesticated cowpea (*Vigna unguiculata* [L.] Walp.)"

**Table S11. Number of SNPs when considering each accession as the “reference” genome, and the resulting union of unique SNPs (merged GVCF) for each accession.**

|  | **IT97K-499-35** | **CB5-2** | **Suvita-2** | **Sanzi** | **UCR779** | **ZN016** | **TZ30** | **Merged (GVCF)** |
| --- | --- | --- | --- | --- | --- | --- | --- | --- |
| **IT97K-499-35** |  | 1,607,267 | 1,660,122 | 1,692,267 | 1,974,970 | 2,092,617 | 1,839,113 | 5,607,053 |
| **CB5-2** | 1,747,504 |  | 1,528,608 | 1,499,900 | 2,501,756 | 1,812,756 | 1,466,499 | 5,480,477 |
| **Suvita2** | 1,766,011 | 1,489,850 |  | 1,512,626 | 2,625,678 | 2,056,752 | 1,847,818 | 5,495,803 |
| **Sanzi** | 1,813,050 | 1,485,875 | 1,539,212 |  | 2,468,787 | 2,002,988 | 1,811,927 | 5,535,733 |
| **UCR779** | 2,029,638 | 2,427,619 | 2,605,364 | 2,417,122 |  | 2,440,589 | 2,424,480 | 5,707,542 |
| **ZN016** | 2,091,894 | 1,692,125 | 1,980,368 | 1,896,387 | 2,382,090 |  | 1,338,143 | 5,419,755 |
| **TZ30** | 1,939,167 | 1,442,388 | 1,865,974 | 1,802,432 | 2,472,527 | 1,422,297 |  | 5,657,433 |
