## Supplemental Table S12 for "A view of the pan-genome of domesticated cowpea (*Vigna unguiculata* [L.] Walp.)"

**Table S12. Number of indels of size 1 to 300 bp when considering each accession as the “reference” genome, and the union set of all indels (merged GVCF) for each accession.**

|  | **IT97K-499-35** | **CB5-2** | **Suvita-2** | **Sanzi** | **UCR779** | **TZ30** | **ZN016** | **Merged (GVCF)** |
| --- | --- | --- | --- | --- | --- | --- | --- | --- |
| **IT97K-499-35** |  | 500,184 | 493,649 | 516,920 | 586,252 | 580,477 | 548,279 | 1,264,602 |
| **CB5-2** | 397,913 |  | 367,662 | 377,767 | 592,079 | 425,572 | 373,435 | 1,145,211 |
| **Suvita2** | 393,125 | 363,343 |  | 372,008 | 619,009 | 472,306 | 446,080 | 1,141,504 |
| **Sanzi** | 413,050 | 373,961 | 372,460 |  | 583,945 | 460,767 | 440,759 | 1,150,696 |
| **UCR779** | 462,494 | 575,960 | 610,235 | 575,146 |  | 556,606 | 572,875 | 1,177,307 |
| **ZN016** | 466,574 | 415,167 | 470,203 | 457,046 | 561,920 |  | 330,037 | 1,131,738 |
| **TZ30** | 435,661 | 364,014 | 442,477 | 436,458 | 579,168 | 331,440 |  | 1,174,076 |
